# The Human Omnibus of Targetable Pockets

**DOI:** 10.1101/2025.09.18.677190

**Authors:** Kristy A. Carpenter, Russ B. Altman

## Abstract

Hundreds of computational methods for predicting ligand binding pockets exist, but the problem of finding druggable pockets throughout the human proteome persists. Different strategies for pocket-finding excel in different use cases. Ensemble models that leverage multiple different pocket-finding strategies can best capture diverse pockets at scale. Despite this, no publicly available human-proteome-wide datasets of pocket predictions from multiple pocket-finding methods exist. We present the Human Omnibus of Targetable Pockets (HOTPocket), a dataset of over 2.4 million predicted pockets over the entire human proteome that utilizes both experimentally-determined and computationally-predicted protein structures. We assembled this dataset by running seven diverse, established pocket-finding methods over all PDB and AlphaFold2 structures of the canonical human proteome. We created a novel pocket scoring method, *hotpocketNN*, which we used to filter candidate pockets and assemble the final proteome-wide dataset. Our *hotpocketNN* method is able to recover known ligand binding pockets, including those which are dissimilar from any pocket seen in its training set. The *hotpocketNN* method outperforms all constituent methods, including P2Rank and Fpocket, when assessing the DCCcriterion on the Astex Diverse Set and PoseBusters dataset. Additionally, *hotpocketNN* was able to identify recently-discovered druggable pockets on KRAS and the mu opioid receptor. We make both the HOTPocket dataset and the *hotpocketNN* method freely available.

**Scientific Contribution:** We introduce HOTPocket, a human proteome-wide dataset of known and predicted binding pockets from a diverse assortment of pocket-finding algorithms. We also introduce *hotpocketNN*, a machine learning model that scores candidate pockets, which we used to filter and harmonize the HOTPocket dataset. Both the data and the model are made publicly available for scientific use.

## Introduction

The time- and cost-intensive nature of the drug development process [1], [2], [3] has motivated widespread effort in developing computational processes, and especially machine learning (ML) models [4], to increase its efficiency. In the earliest stages of drug development, computational models have been developed to design novel druglike molecules [5], [6] and predict their ADMET properties [7], [8], [9]. Additionally, computational models can play a key role in developing drug leads for a target of interest. A common paradigm in structure-based drug discovery is to first identify the active site or other putative binding pocket on the desired target, then to design a small molecule that can bind in the pocket [10], [11]. Pocket identification can be done experimentally, but computational pocket-finding methods are desirable for their higher speed, lower cost, and increased ability to use at scale. There is a plethora of computational pocket-finding methods available [12], [13], [14], [15], [16], [17], underscoring the importance and unsolved nature of the pocket identification task.

There are five typical strategies for pocket-finding; methods can be geometry-based, energy-based, template-based, conservation-based, or ML-based. Geometry-based methods search for concave regions on the protein surface that could accommodate a small molecule. Energy-based methods use probes and physics-based energy functions to search for regions on the protein surface that energetically favor small molecule binding. Template-based methods search for regions of a protein that match sequence or structure motifs from known pockets. Conservation-based methods use the principle that small molecule binding sites are usually critical to protein function and are therefore evolutionarily conserved. ML-based methods use training sets of known binding pocket sequences or structures, as well as a variety of machine learning and deep learning architectures, to learn how to classify pockets on unseen proteins. Each of these pocket-finding strategies has strengths and weaknesses. For example, geometry-based methods by definition can detect any concave pocket but cannot detect cryptic pockets that appear flat in unbound form but undergo conformational change upon ligand binding [18]. Likewise, ML-based methods can implicitly learn a combination of other strategies as well as latent patterns, but can be biased by their training data and are largely uninterpretable. As such, different kinds of pockets, or different downstream use cases, may be best suited by different types of pocket-finding methods, *e.g.* searching for classical small molecule binding cavities may be best served by a geometric method, whereas searching for a cryptic pocket or a macrocyclic peptide binding site may be best served by a conservation-based or ML-based method.

Combining pocket-finding strategies can improve performance, both in terms of being able to capture diverse pockets and increasing confidence in pockets predicted independently by different methods. Ensemble methods have found prior success in pocket-finding. MetaPocket [19] and MetaPocket 2.0 [20] combine predictions from four and eight pocket finding methods, respectively. Both methods achieved higher pocket-finding performance than any individual constituent method, and MetaPocket 2.0 improved performance over the original MetaPocket. TargetCom is another consensus method that combines four existing methods to outperform each individually [21]. SURFNET-Consurf [22], LIGSITEcsc [23], and ConCavity [24] are all consensus methods that combine geometry and conservation to outperform purely geometric methods. More recently, CoBDock, a blind docking method that integrates multiple docking and pocket-finding methods, was shown to outperform individual state-of-the-art docking and binding site prediction methods [25]. Ensembling has also been shown to increase performance over any individual constituent method in the related task of ligand binding affinity prediction [26].

Protein structure prediction models have recently extended to predict the structure of proteins in complex with other proteins, nucleic acids, and small molecule ligands. This framework was introduced by RoseTTAFold All-Atom [27] and AlphaFold3 [28], then followed by several groups aiming to create more openly available reproductions of AlphaFold3 [29], [30], [31], [32] before its authors shared code [33]. These methods purport to predict the structures of protein-ligand interactions without prior knowledge of the binding pocket. While exciting advances in biomolecular complex prediction with potential to aid drug discovery, these methods have not entirely solved the problem of predicting interactions between proteins and small molecules [34], [35], [36], nor have they solved the problem of ligand-agnostic pocket identification. In order to predict a pocket using these methods, one must specify a ligand. Because not all ligands will bind all binding pockets, and because it is infeasible to test all possible protein-ligand combinations across the whole proteome and whole universe of possible ligands, one cannot use AlphaFold3 or similar methods to enumerate all potential pockets in the human proteome. Even at a small scale, a ligand-agnostic pocket-finding method allows medicinal chemists to design a novel molecule that binds the target of interest in a way that AlphaFold3 and similar methods are not suited for.

Despite hundreds of pocket-finding methods having been published, very few make pre-computed pockets accessible. We previously conducted a literature search for databases of ligand-binding pocket structures and identified 58 databases of pocket structures [37]. Of these, only 37 databases (63.8%) were still accessible via website, download, or application programming interface (API) as of 2024. Approximately half of these databases (18, or 48.6%) contain only known pockets extracted from protein-ligand interactions in the Protein Data Bank (PDB) [38] or the scientific literature. Of the remaining 19 databases that contain novel predicted pockets, 8 (42.1%) contain pockets predicted on AlphaFold2-generated [39] protein structures and 11 (57.9%) only contain pockets from experimentally-determined protein structures from the PDB. Because the majority of proteins in the human proteome have not yet had their three-dimensional structures experimentally determined, databases that do not leverage computationally-predicted structures exclude most of the human proteome. All 8 of the databases that include pocket predictions on computationally-predicted structures use a single pocket prediction method to generate their data, *i.e.* there does not yet exist a database of pocket structures that: 1) includes known and predicted pockets; 2) leverages multiple different pocket-finding methods; 3) includes both experimentally-determined and computationally-predicted protein structures; and 4) is publically accessible. Such a database would allow end users to browse diverse pocket knowledge over the entire human proteome, harmonized into a consistent format and in one place, as opposed to wrangling and comparing the outputs of several different resources and tools.

Here, we introduce the Human Omnibus of Targetable Pockets (HOTPocket), a dataset of over 2.4 million pocket predictions from an array of established, diverse pocket-finding methods over the entire human proteome. Pockets are filtered and scored using *hotpocketNN*, a neural network that uses per-residue protein language model embeddings as features. We compare the performance of *hotpocketNN* to that of each of its constituent pocket-finding methods on a subset of BioLiP, the Astex Diverse Set, and PoseBusters. We demonstrate that *hotpocketNN* is able to recover recently-discovered druggable pockets on two high-interest targets: KRAS and the mu opioid receptor. The main contributions of this work are twofold:

1. A novel dataset (HOTPocket) of diverse, harmonized, filtered, and scored binding pocket predictions across the entire human proteome for use in drug development; and
2. A novel method (*hotpocketNN*) for scoring and filtering predicted binding pockets. Both of these contributions are made freely available for scientific use at github.com/Helix-Research-Lab/HOTPocket.

## Methods

### Assembly of human proteome structures

We obtained the canonical sequences of the human reference proteome from UniProt (version September 2023) [40]. We obtained all experimentally-determined protein structures corresponding to these UniProt accessions from the RCSB Protein Data Bank (PDB) (version September 2023) [38]. We obtained all computationally-predicted protein structures corresponding to these UniProt accessions from the AlphaFold Protein Structure Database (version July 2021) [41].

### Selection of pocket-finding methods

We sought to leverage the power of ensembling by compiling proteome-wide pocket predictions from several different pocket-finding methods. We conducted a broad literature search for methods that identify ligand binding sites from static protein structures or single protein sequences. While impossible to exhaustively identify all pocket-finding methods ever published due to the large number of such methods, we assembled a list of over 130 pocket-finding methods (**Table S1**). Of these, we selected methods for downstream use based on three criteria: 1) selected methods must have code, packages, APIs, or large-scale pre-computed predictions available; 2) selected methods should show exemplary performance and/or have seen wide use in the community; and 3) the set of selected methods should span a diverse range of pocket-finding strategies.

### Assembly of proteome-wide pocket predictions and ground truth pockets

#### AutoSite

We obtained AutoSite pocket predictions by running the AutoSite package^1^ (version 1.0, build 5) accompanying the AutoSite publication [42] on all assembled human proteome structures. To prepare the input PDB files, we removed any non-ATOM lines, then used Open Babel (version 2.4.0) [43], with the pH set to 7.4 for hydrogen addition and the rigid molecule flag, to convert to PDBQT format. We ran AutoSite on the PDBQT files using default parameters. The AutoSite program outputs the coordinates of points making up the predicted pocket space. We created predicted pockets by taking all residues within 5A of any predicted pocket space point.

#### CASTp

We obtained CASTp pocket predictions from the CASTp website^2^ [44]. Because the CASTp website only contains precomputed predictions for structures from the PDB, and because the code is not made publicly available, we could not generate predicted pockets for AlphaFold2 structures.

#### CAVITY

We obtained CAVITY pocket predictions from the CavitySpace website^3^. We downloaded all AlphaFold and PDB cavity predictions (version June 2022) and extracted all pockets for all assembled human proteome structures.

#### Fpocket

We obtained Fpocket pocket predictions by running the fpocket package^4^ (version 4.1, available on conda) on all assembled human proteome structures. We stripped any heteroatoms from the input PDB file before running. We ran the program with default parameters.

#### LIGSITEcs

We obtained LIGSITEcs pocket predictions by running the lcs package (from BALL^5^ version 1.1.1) accompanying the LIGSITEcsc publication [23] on all assembled human proteome structures. We stripped any heteroatoms from the input PDB file before running. We ran the program with default parameters. The LIGSITEcs program outputs the center of mass of the predicted pocket space. We expanded each pocket center to a full predicted pocket of comparable size to pockets from other methods by adding all residues within 8A of the central point, following the pocket-building method detailed in [23].

#### P2Rank

We obtained P2Rank pocket predictions by running the P2Rank package^6^ (release 2.5) accompanying the P2Rank publication [45] on all assembled human proteome structures. We used the default configuration for experimentally-determined structures and used the alphafold configuration for AlphaFold2-predicted structures.

#### PocketMiner

We obtained PocketMiner pocket predictions by running the PocketMiner code provided in the GitHub repository^7^ (cloned July 2024) accompanying the PocketMiner publication [46] on all assembled human proteome structures. We stripped any heteroatoms from the input PDB file before running. PocketMiner outputs per-residue cryptic pocket scores and does not cluster hotspot residues into pockets. When applying their method, the PocketMiner authors averaged the PocketMiner scores of a residue and its 10 neighbors to obtain a hotspot score. A neighborhood of residues was classified as a predicted cryptic pocket if it had a hotspot score of 0.7 or higher [46]. We leveraged this strategy to obtain predicted pockets from the single-residue Pocketminer scores.

#### “Ground truth” pockets

We selected the BioLiP2 database [47] as our source of experimentally-determined pockets with which to evaluate predictions. We obtained BioLiP2 annotations for all assembled human proteome structures from the redundant ligand-protein interaction dataset (version September 2023) downloaded from the BioLiP website^8^, which accompanies the BioLiP [48] and BioLiP2 [47] publications.

### Naive pocket ensemble prediction

We assessed how well the proteome-wide predicted pockets recovered known binding pockets. We assembled a dataset of known pockets of druglike ligands by filtering our “ground truth” dataset, BioLiP2, for proteins included in the Therapeutic Target Database (TTD) [49] to select for human proteins known to bind drugs. We filtered the resulting structures to select only pockets with an organic small molecule ligand (**Text S1**). This process resulted in a validation dataset of 778 pockets from 482 unique PDB structures.

We introduced a naive consensus approach for pocket identification: taking the intersection or union of the sets of residues predicted to be in pockets by each of the included pocket prediction methods. We selected any residue that was predicted to be part of a pocket by at least *k* of the pocket-finding methods to be a hotspot residue, with *k* ranging from 1 to 7. We calculated the ROC curve and AUROC, varying *k*, to assess per-residue prediction performance on the constructed BioLiP-TTD dataset. We compared the naive ensemble approaches to each of the seven constituent methods individually. In addition to predicting consensus pockets from all predictions of all methods, we repeated the process only considering predicted pockets that passed a simple filtering scheme (**Text S2**) in an attempt to remove low-confidence pocket predictions.

### Machine learning-based pocket ensemble prediction

We developed an ML model to discard unlikely predictions and enrich the final predicted pocket dataset for the most plausible pockets. We tested three different classes of ML model: logistic regression, fully-connected neural network (NN), and convolutional neural network (CNN).

The input to each ML filtering model was a proposed pocket in the form of an D X D matrix, where D is the number of residues and D is the number of features. We set D to be 15 as this was slightly larger than the overall average predicted pocket size (11.58 amino acids). The matrix representation of any pocket with fewer than 15 residues was padded with zeros to size D X D. Any pocket with more than 15 residues was randomly truncated to a residue subset of size 15.

We tested three different schemes to featurize each of the *N* residues in the proposed pocket: Feature Set A: the per-residue pocket predictions from each of the constituent methods; Feature Set B: the per-residue ESM2 [50] embeddings; and Feature Set C: both the per-residue pocket predictions and per-residue ESM2 embeddings concatenated together. Each feature set is applied for only the residues in the proposed pocket, not all residues in the protein.

With Feature Set A, D is equal to the number of constituent pocket-finding methods (7). Element (DD_D_, D_D_) is equal to 1 if the Dth amino acid is predicted to be in a pocket by the Dth pocket-finding method, and is otherwise equal to 0. With Feature Set B, D is equal to the embedding size (2560). The Dth row is equal to the ESM2 embedding for the Dth amino acid. With Feature Set C, D is equal to the sum of the number of constituent pocket-finding methods and the embedding size (2567), with the features for the Dth row are the predictions for the Dth amino acid from the 7 constituent methods and the ESM2 embedding for the Dth amino acid, concatenated. The CNN architecture accepted two-dimensional inputs; the logistic regression architecture and the NN architecture required the D X D matrix to be flattened into a one-dimensional vector of length D * D. The output of each model is a score between 0 and 1 representing the probability that the input proposed pocket is an actual pocket.

We created a dataset with positive and negative pocket examples to train, validate, and test the ML models. We partitioned the proteins from the BioLiP2/TTD dataset (described in “Naive pocket ensemble prediction”) based on global sequence identity; no structure in any partition had more than 40% sequence identity with any structure in any other partition. We split the data using a 80:10:10 ratio for training, validation, and test splits. All protein chains present in a single structure were kept in the same split. To create inputs for the ML models, we augmented the BioLiP2-annotated data by generating multiple “positive examples” (residue sets labeled as pockets) and multiple “negative examples” (residue sets labeled as non-pockets) for each pocket reported in BioLiP2.

The number of positive examples generated per BioLiP2 pocket depended upon the pocket’s size: 2 positives for a BioLiP2-annotated pocket of fewer than 10 residues; 3 positives for a BioLiP2-annotated pocket of 10 to 24 residues; 4 positives for a BioLiP2-annotated pocket of 25 or more residues. The number of negative examples generated for a given BioLiP2 pocket was twice the number of positive examples for that same pocket. We created positive examples of binding pockets in an iterative process. We sampled a random residue annotated by BioLiP2 as being part of a binding site and created a “pocket sphere” by selecting all residues within a 5A radius. To ensure that the pocket sphere is not too different from the actual binding pocket, we only added the pocket sphere as a positive example if either >50% of the included residues were annotated by BioLiP2 as being part of the binding site, or that the set of included residues contained >80% of all BioLiP2-annotated residues for the binding site. To avoid redundancy, we asserted that no two positive examples should have 90% or more residue overlap. We continued sampling residues from the set of BioLiP2-annotated residues until we had the desired number of positive examples or all BioLiP2-annotated residues were examined as potential pocket centers. We repeated this process for each non-redundant BioLiP2 pocket of each structure in our dataset. We created negative examples using the assumption that the vast majority of randomly-sampled patches of a protein surface that do not overlap with known pockets will not be pockets. For each structure, we took all residues within a 5A radius of a randomly-selected residue, and kept it as a negative example if none of the residues contained in the sphere were labeled by BioLiP2 as part of a binding pocket. We again asserted that no two negative examples should have 90% or more overlap. This process (**Figure S1**) yielded a total of 1357 positive examples and 2577 negative examples.

We performed hyperparameter tuning on each model. For the logistic regression models, we varied the penalty (no penalty, L1, L2, or elastic net) and regularization weight, and trained the model until convergence or a maximum of 500 iterations. We selected the best hyperparameter setting for each feature set by validation AUC. For the NN models, we varied the learning rate, dropout, batch size, optimizer, and number of dense layers. For the CNN models, we varied the filter size, pooling size, number of convolutional layers, and all previously stated NN hyperparameters. We trained each model for 50 epochs in triplicate, with the same three random seeds used for each hyperparameter setting. We selected each model’s best epoch and best set of hyperparameters according to the AUC on the validation set. We averaged the best validation AUC for each model across the three random seeds, and used this measure to select the best hyperparameter setting for each feature set.

We evaluated different manners of ordering the amino acid representations within each pocket: either always putting the amino acids in N-terminus to C-terminus order, or shuffling the order. Shuffling the order enabled us to augment our dataset by including up to ten permutations of amino acid order for each pocket, yielding a nearly 10x increase in dataset size. We compared the validation set performance of the best NN and CNN models trained with each amino acid ordering representation: N-to-C (1x data with consistent residue order), shuffled (1x data with shuffled residue order), and augmented (10x data with shuffled residue order).

We identified the best model overall for downstream use by validation AUC. We selected a score threshold *t* to discriminate between pockets and non-pockets by maximizing the difference between the true positive rate and the false positive rate. We only evaluated our selected final model on the test set to avoid biasing results.

### Assessment of filtered pockets

We visually examined select protein structures to preliminarily assess the ability of the ML model to improve pocket prediction. Selected structures included a random protein from each of the validation and testing sets, as well as two random proteins from BioLiP with structures experimentally solved for the first time in 2024 (after our cutoff for BioLiP downloads to obtain training, validation, and testing sets) with less than 50% sequence identity to any protein that had a BioLiP entry prior to 2024.

We compared the residue-wise performance of our ensembling and filtering method (henceforth called *hotpocketNN*) to the existing pocket-finding methods with ROC and PRC curves. Each residue of each structure had a true label of whether or not it was included in a known pocket and a predicted label of whether or not a given method predicted it as part of a pocket. To calculate the ROC and PRC curves, we aggregated all residue-wise labels over all structures in the database. We compared two versions of *hotpocketNN* (one with ESM2 embeddings as features, and one with ESM2 embeddings and constituent method prediction labels as features), the naive ensembling method described above, and all seven constituent pocket prediction methods. The *hotpocketNN* score for a given residue was the maximum score generated by the ML filtering model of all candidate pockets containing the residue. Likewise, the Fpocket, P2Rank, and PocketMiner scores for a given residue were the maximum score generated by the respective method of all candidate pockets generated by the respective method containing the residue. As AutoSite, CavitySpace, CASTp, and LIGSITEcs do not generate pocket scores, all residues present in any candidate pocket from each respective method were given a predicted label of 1, and all residues not present in any candidate pocket from each respective method were given a predicted label of 0. We created the ground truth labels by assigning each residue within a radius *r* of the relevant ligand in the experimentally-determined structure a label of 1, and all other residues a label of 0. We calculated the ROC and PRC curves using the pocket radii of 5A and 10A.

We used the DCCcriterion to evaluate performance at the pocket level. The DCCcriterion requires computing the central coordinate of the top-*N* ranked pockets for each structure in the dataset, computing the ligand-to-center distances (defined as the distance between the central coordinate and the nearest atom of the nearest relevant ligand), and computing the fraction of distances that are less than 4A. We define D_D_ as the number of biologically-relevant ligands (as defined by BioLiP) in the structure, and calculate the DCCcriterion when *N* is set to D_D_ and when *N* is set to D_D_ + 2. Using D_D_ demonstrates if the method in question can correctly rank pockets over non-pockets, and the D_D_ + 2 case gives buffer for small ranking mistakes; these settings have precedent in prior work [45], [51].

We visualized the full distribution of ligand-to-center distances for the top-*N* ranked pockets for each structure. We calculated a background distribution by taking *N* random points along the surface of each structure and computing the distance between each point and the nearest atom of the nearest relevant ligand.

To assess generalizability, we conducted this analysis over our whole human BioLiP dataset and not the BioLiP / TTD subset, which was used in training, validation, and testing of the ML filter model. Because many BioLiP structures were used to train and validate both our ML model and most of the constituent pocket prediction methods, we also evaluated on two additional datasets: the Astex Diverse Set [52] and PoseBusters [53]. The Astex Diverse Set was released in 2007 and includes 85 structures of diverse sequence with pharmaceutically-relevant ligands [52]. The PoseBusters dataset was released in 2024 and includes 428 structures released after 2021 of diverse sequence with pharmaceutically-relevant ligands [53]. In addition to the original datasets in their entirety, we also calculated the ROC, PRC, and DCCcriterion on the subset of structures that were not present in the training or validation sets of our ML model to avoid inflating estimated performance.

### Preparation of output proteome-wide prediction set

We computed ESM2 embeddings for all protein sequences in our assembled set of human proteome structures and featurized all candidate pockets proteome-wide. We input all candidate pockets to the ML model to obtain scores and discarded any candidate pockets with a score less than 0.4 (our empirically-determined *t*). We also discarded any candidate pockets on AlphaFold2-predicted structures with an average pLDDT of less than 70, which is the threshold used by the AlphaFold2 Protein Structure Database to classify a residue as “low confidence”. See **Figure 1** for a schematic of the full dataset assembly process.

**Figure 1.**
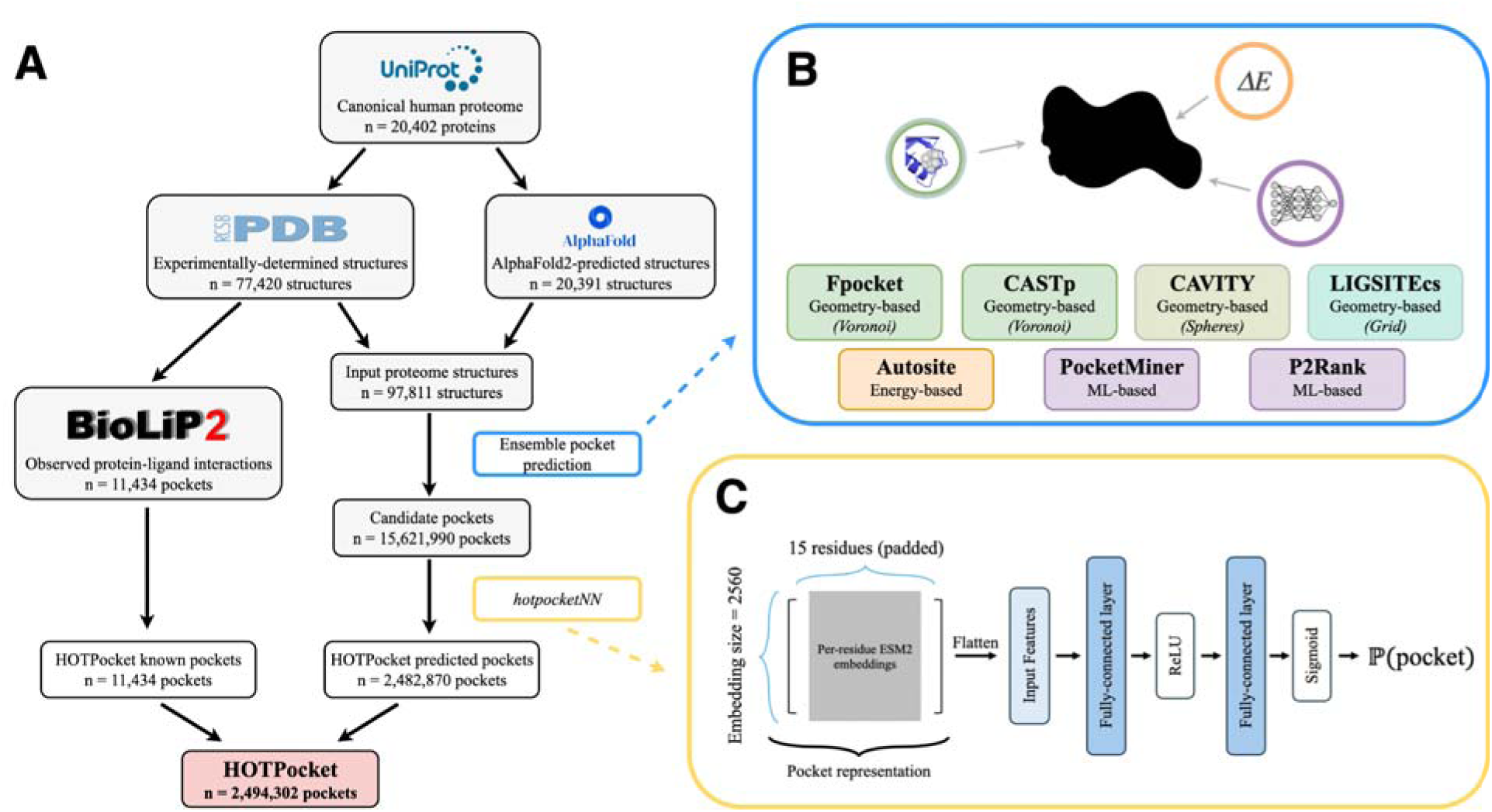
Schematic representation of the HOTPocket dataset assembly process. **a)** We assembled structures of the canonical human proteome from the PDB and AlphaFold2 Protein Structure Database, did ensemble pocket prediction on all structures, then filtered the subsequent candidate pockets using our *hotpocketNN* model. The resulting filtered pockets constitute the predicted pocket portion of the HOTPocket database, which is harmonized with known pockets annotated in the BioLiP2 database. **b)** “Ensemble pocket prediction” step: We applied seven diverse pocket finding methods to each human proteome structure. These methods fall broadly into the categories of geometry-based (shades of green), energy-based (orange), and machine learning-based (purple). **c)** “*hotpocketNN*” step: We used a neural network of the specified architecture to score and filter candidate pockets. Inputs to the model are ESM2 embeddings for each residue in the pocket, with pocket size padded (or truncated) to 15 residues. The output of the model is a score between 0 and 1 representing the probability that the input residue set is a binding pocket.

### Case studies

#### KRAS

KRAS is a target of high interest for cancer therapeutics, but has been long considered an “undruggable” protein [54], [55]. In 2019, Kessler *et al.* discovered a KRAS inhibitor that bound the “switch I/II” pocket [56]. Our first case study was to investigate if *hotpocketNN* could rediscover this pocket.

We defined the switch I/II cryptic pocket residues to be all residues within 5A of the F0K ligand in PDB structure 6gj8, released with the Kessler *et al.* paper. We ran *hotpocketNN* using Feature Set B on 6gj8 to test the easiest case: if *hotpocketNN* can identify the switch I/II pocket on a structure that has a ligand bound in that pocket (without considering the ligand itself during prediction).

We sought a KRAS structure released before 2012, which marked the beginning of a period when multiple papers identified plausible druggable pockets on KRAS [56], [57], [58], [59]. The only such structure that contained a full KRAS protein, as opposed to a peptide, was 3gft. The only biologically-relevant organic ligand in the 3gft structure is a GTP analogue. We ran *hotpocketNN* on 3gft to test if our method can identify the switch I/II pocket on a structure that was solved before the switch I/II pocket or any similar pocket was discovered.

Since KRAS is in the inactive, GDP-bound state in the 6gj8 structure, but is in the active, GTP-bound state in the 3gft structure, we sought additional GDP-bound KRAS structures without a ligand in the switch I/II pocket on which to assess *hotpocketNN*. We identified four such structures: 5uk9, 6bp1, 6quu, and 7lz5. We ran *hotpocketNN* on these structures to test if *hotpocketNN* can identify the switch I/II pocket from an inactive-state structure that does not have a ligand bound in that pocket.

#### Mu opioid receptor

The mu opioid receptor (mOR) is a G-protein coupled receptor (GPCR) that is the primary target for opioids such as fentanyl, codeine, and methadone. In addition to the main active site, an allosteric mOR pocket exists [60], [61]. Two separate groups recently found ligands that bind allosteric pockets on mOR [62], [63] and deposited structures displaying this interaction in the PDB, which were all released in 2024. One allosteric site is located inside the transmembrane helical bundle near the orthosteric site (observed in structure 8k9l). The other is on the outside of the helical bundle (observed in structure 9bjk). We ran *hotpocketNN* on three mOR structures from the PDB: the structure with a ligand bound allosterically outside the helical bundle (8k9l), a structure from the same study with no ligands (8k9k), and a structure released in 2022 of fentanyl bound to mOR with no allosteric ligands (8ef5). We did not run *hotpocketNN* on structure 9bjk because it was not available in the legacy PDB format, which is required by the pocket finding methods used by *hotpocketNN*. We assessed if *hotpocketNN* was able to recover the orthosteric pocket and both allosteric pockets on each of the three structures.

## Results

### Assembly of human proteome structures

We obtained a total of 97,811 protein structures on which to predict pockets. Of these, 77,420 structures were experimentally-determined and originated from the PDB, and 20,391 structures were computationally-predicted and originated from the AlphaFold Protein Structure Database. This set of protein structures represented 20,402 unique human proteins.

### Selection of pocket-finding methods

We selected 7 pocket-finding methods to apply to the human proteome: AutoSite [42], CASTp [44], [64], CAVITY [65], [66], Fpocket [51], LIGSITEcs [23], P2Rank [45], and PocketMiner [46]. Four of these methods employ diverse geometry-based strategies: Voronoi tessellation (Fpocket and CASTp), probe spheres (CAVITY), and grid meshes (LIGSITEcs). Another method is energy-based (AutoSite). The two remaining methods are ML-based – one of which uses a Random Forest (P2Rank) and the other, which specifically focuses on cryptic pockets, uses a graph neural network (PocketMiner).

### Assembly of proteome-wide pocket predictions and ground truth pockets

We compiled a total of 11,434 known pockets and 15,621,990 predicted pockets across the 97,811 protein structures and 20,402 unique proteins of the human proteome (**Table 1**). BioLiP, CASTp, and LIGSITEcs did not produce any pockets for computationally-predicted structures (due to only annotating known pockets from experimentally-determined structures, only having predictions from experimentally-determined structures available, and errors arising when applying the method to AlphaFold2 structures, respectively), resulting in a marked decrease in number of proteins covered when compared to other methods (**Table S2**). AutoSite, CASTp, Fpocket, and PocketMiner tended to predict far more pockets per structure than the other methods, leading to their dominance in the number of pockets predicted overall.

**Table 1.** Summary of assembled proteome-wide pocket dataset. BioLiP pockets are derived from experimentally-determined protein complex structures; all other pockets are computational predictions. Number of proteins represented is determined by the number of unique UniProt IDs. Pocket size is defined as the number of amino acids constituting the pocket.

| Method | # pockets | # proteins | Mean pockets per structure (*std.*) | Median pockets per structure | Mean pocket size (*std.*) |
| --- | --- | --- | --- | --- | --- |
| BioLiP | 11,434 | 4,437 | 2.14 (2.45) | 1 | 8.94 (9.02) |
| AutoSite | 2,303,329 | 20,171 | 33.16 (46.24) | 20 | 16.69 (11.55) |
| CASTp | 2,225,879 | 5,080 | 72.07 (77.58) | 47 | 8.22 (29.94) |
| CAVITY | 88,916 | 14,999 | 4.82 (4.46) | 3 | 36.57 (32.10) |
| Fpocket | 2,899,488 | 20,359 | 39.15 (66.81) | 22 | 10.26 (7.72) |
| LIGSITEcs | 166,916 | 7,083 | 3.17 (0.86) | 3 | 15.55 (4.99) |
| P2Rank | 750,966 | 16,996 | 11.37 (26.75) | 5 | 12.83 (8.00) |
| PocketMiner | 7,186,496 | 20,015 | 119.16 (203.59) | 53 | 11.00 (0.00) |

Average pocket size (measured by number of residues included in the pocket) varied but typically was in the 8-15 residue range. The exceptions to this were CAVITY, which tended to predict fewer and larger pockets than other methods, and PocketMiner, which always generated pockets of size 11 (a central amino acid and its ten nearest neighbors) due to how we converted its high-scoring hotspot residues into predicted pockets (see Methods). We also note that PocketMiner had by far the most number of predicted pockets, both per-structure and overall. As many of the generated pockets are highly redundant, this is indicative of a suboptimal clustering strategy, and should not be taken as evidence of the existence of millions of cryptic pockets throughout the human proteome.

### Naive pocket ensemble prediction

There were 735 PDB structures that had BioLiP2 annotations and at least one predicted pocket from each of the seven constituent pocket prediction methods. We randomly selected two of these structures to visualize (PDB IDs: 1wl4 and 4nst) (**Figure 2**). AutoSite, CASTp, CAVITY, Fpocket, and PocketMiner tended to predict more residues as being part of pockets than LIGSITEcs and P2Rank, but upon applying the filtering scheme the magnitude of this difference decreased. For 1wl4, all seven methods correctly predicted the BioLiP2-annotated binding pocket, albeit with varying levels of precision (**Figure 2a**). In this case, the filtering scheme eliminated many, but not all, false positives. Only AutoSite, CASTp, Fpocket, and P2Rank predicted pockets overlapping with the BioLiP2-annotated binding pocket for 4nst – after filtering, this was reduced to just AutoSite and P2Rank (**Figure 2b**). The filtering scheme excluded all predicted pockets from CAVITY, Fpocket, and PocketMiner.

**Figure 2.**
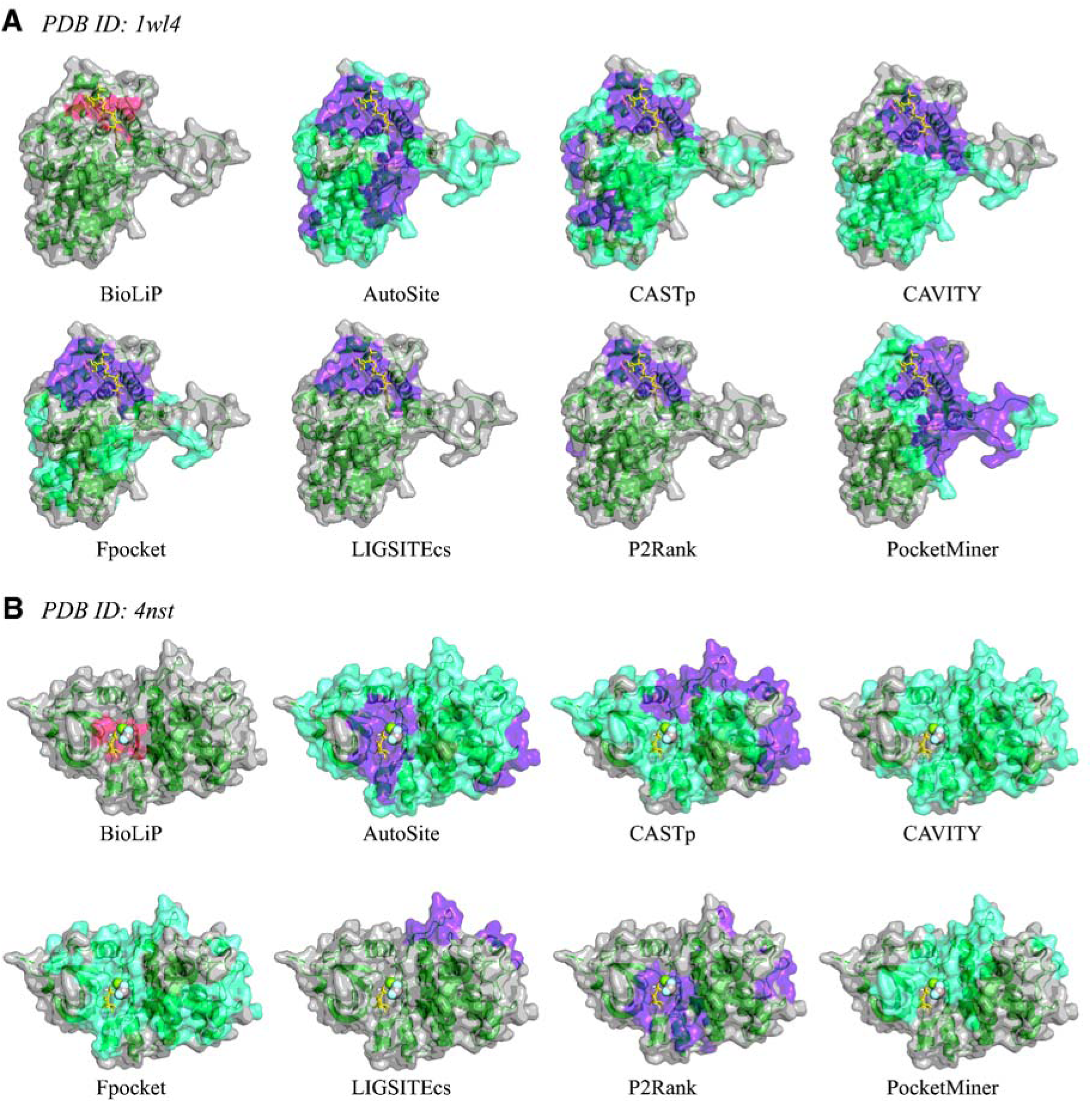
Visualization of predicted pockets from all seven constituent methods as compared to the known binding pockets annotated in BioLiP2 for PDB structures **a)** 1wl4, and **b)** 4nst. Known binding pockets are highlighted in pink; predicted binding pockets that pass the simple filter (see filter description in **Text S2**) are highlighted in purple; all other predicted binding pockets are highlighted in green cyan. Biologically-relevant ions are shown as spheres and biologically-relevant organic small molecules are shown with yellow sticks. Predicted pockets are shown in aggregate, *i.e.* individual pockets are not distinguished from each other and thus may appear as one large mega-pocket.

We examined the overlap between constituent method predictions using a heatmap-style visualization (**Figure 3**). For both the unfiltered and filtered version of the 1wl4 heatmap, the hotspot of residues with 6 or 7 constituent methods predicting pocket membership that emerged is in the general ligand binding location as annotated in BioLiP2 (**Figure 3a**). However, in the 4nst unfiltered heatmap, multiple such hotspots emerged, with only one overlapping with the BioLiP2-annotated pocket (**Figure 3b**). Because not all of the constituent pocket-finding methods produced pockets that passed the simple filter for the 4nst structure, no hotspots remained after filtering. The BioLiP2-annotated pocket was still predicted by two methods after filtering, but the top predicted residues were distal to the binding site. For both 1wl4 and 4nst, we again observed that the simple filter eliminated some, but not all, false positives.

**Figure 3.**
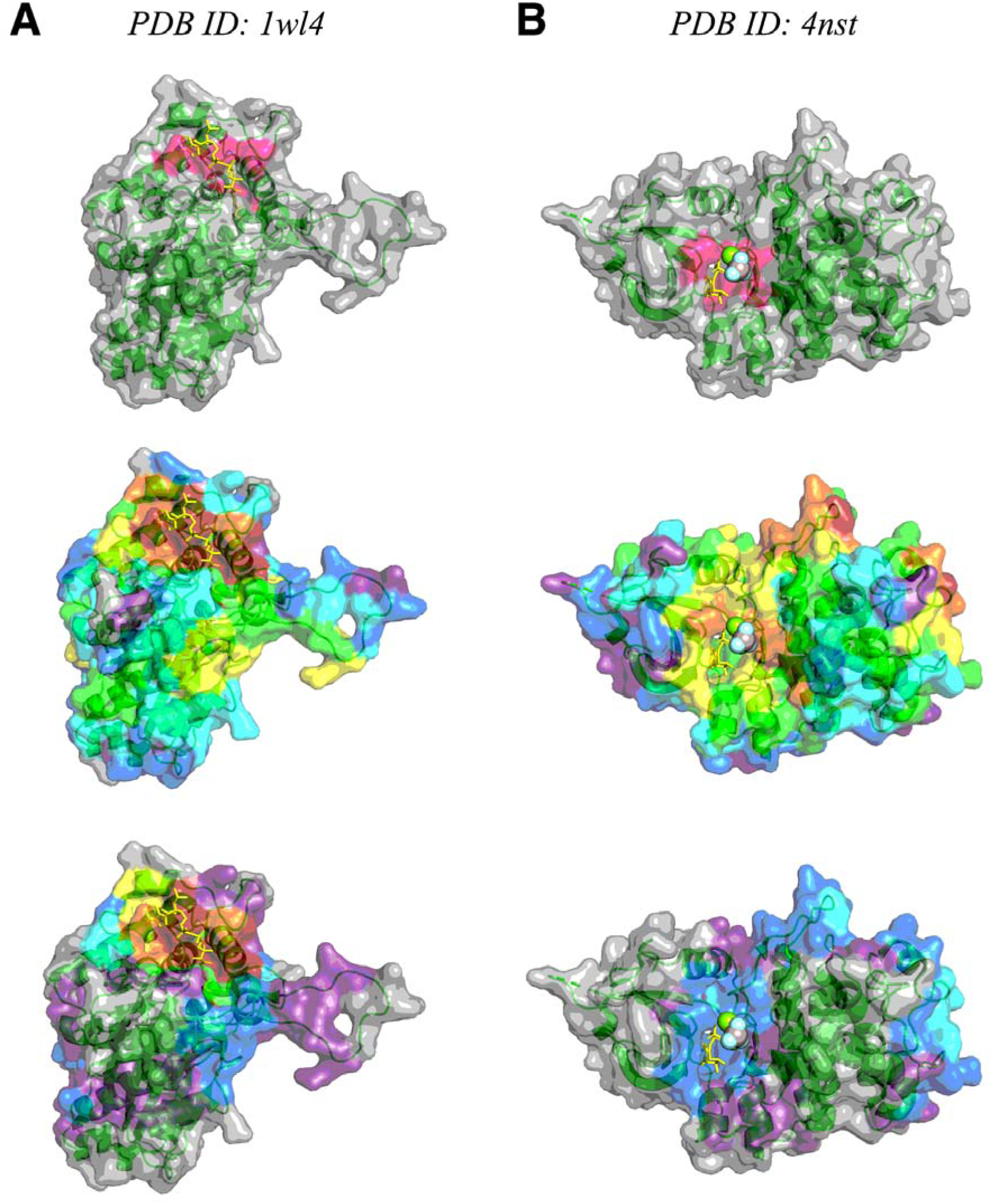
Heatmap visualization of predicted pockets as compared to known binding pockets annotated in BioLiP2 for PDB structures **a)** 1wl4, and **b)** 4nst. The heatmap colors correspond to how many of the constituent methods predicted that a given residue is part of a pocket: gray = 0 methods; purple = 1 method; blue = 2 methods; cyan = 3 methods; green = 4 methods; yellow = 5 methods; orange = 6 methods; red = all 7 methods. Top: BioLiP2 annotated known binding pocket shown in pink. Middle: heatmap of all predicted pockets. Bottom: heatmap of filtered predicted pockets.

When assessing per-residue predicted labels over the BioLiP-TTD dataset, we found that P2Rank performed the best, followed by Fpocket and the naive filtered ensembling approach (**Figure 4**). The unfiltered version of the naive ensembling approach performed poorly, and all individual pocket prediction methods except for CAVITY achieved a higher AUC. This indicates that, while the simple filtering approach described above boosts the ensemble approach’s performance, a better filtering and scoring approach is required in order to better harmonize predictions from constituent methods and outperform each individually.

**Figure 4.**
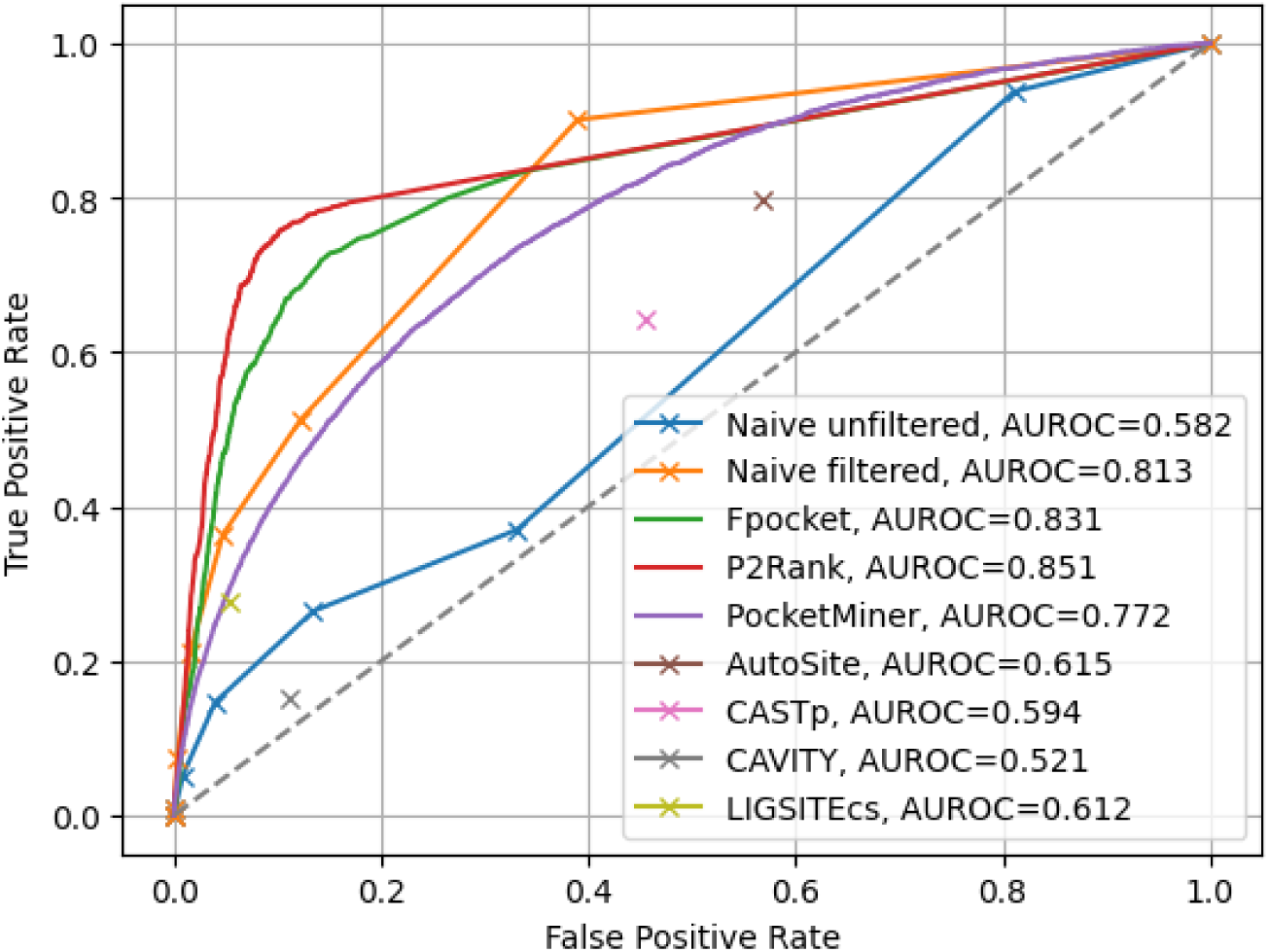
ROC curve for unfiltered and filtered predicted pockets. True positive rate and false positive rate are calculated based on the per-residue labels across all structures in the BioLiP-TTD dataset (N = 482 structures) at each threshold of how many constituent methods need to predict a residue as part of a pocket in order to classify that residue as being part of a pocket. For Fpocket, P2Rank, and PocketMiner, each residue’s predicted score for being a member of a pocket was the maximum of all predicted pockets of which it was a member. AutoSite, LIGSITEcs, CAVITY, and CASTp do not provide pocket scores; for these methods, each residue’s predicted score for being a member of a pocket was 1 if it is part of any predicted pocket for the structure, or 0 otherwise. Ground truth residue labels are based on exact match to annotated binding sites in BioLiP.

### Machine learning-based pocket ensemble prediction

Of the three ML algorithms evaluated, the feedforward neural network performed the best with regard to validation set AUC, followed closely by the convolutional neural network, and the logistic regression models performed the worst (**Table 2, Table S3, Table S4**). Models performed better when provided the ESM2 embeddings as features, and adding the pocket predictions from each of the constituent models as per-residue features slightly boosted performance. The order in which residues were listed in the proposed pockets had little effect on model performance, with the “augmented” framework slightly outperforming the others due to its 10x increase in training data size (**Figure S2**).

**Table 2.** Results of hyperparameter tuning. For each ML model type and feature set pairing, the best validation AUC (averaged over three random seeds) across all hyperparameter combinations is listed. The corresponding training AUC for the epoch in which the best validation AUC was achieved for the best hyperparameter setting, averaged over three random seeds, is also listed. Feature set A is the per-residue pocket predictions from each of the constituent methods; Feature set B is the per-residue ESM2 embeddings; Feature set C is both the per-residue pocket predictions and per-residue ESM2 embeddings concatenated together.

| Model type | Feature set | Best mean train AUC (*std.*) | Best mean val AUC (*std.*) |
| --- | --- | --- | --- |
| Logistic regression | A | 0.7779 (0.0000) | 0.7823 (0.0000) |
| Logistic regression | B | 0.9894 (0.0000) | 0.8468 (0.0000) |
| Logistic regression | C | 0.9843 (0.0000) | 0.8520 (0.0000) |
| NN | A | 0.8582 (0.0107) | 0.8745 (0.0005) |
| NN | B | 0.9963 (0.0048) | 0.9208 (0.0018) |
| NN | C | 0.9995 (0.0002) | 0.9344 (0.0019) |
| CNN | A | 0.8803 (0.0076) | 0.8748 (0.0010) |
| CNN | B | 0.9966 (0.0022) | 0.8907 (0.0145) |
| CNN | C | 0.9964 (0.0026) | 0.9072 (0.0028) |

Based on the hyperparameter tuning results, we selected two models with which to proceed: the NN with Feature Set B (ESM2 embeddings only) and the NN with Feature Set C (ESM2 embeddings and constituent method pocket predictions). The former used a single feedforward hidden layer, a batchsize of 32, a dropout probability of 0.5, a learning rate of 0.001, and the AdamW optimizer; the latter used a single feedforward hidden layer, a batchsize of 64, a dropout probability of 0.4, a learning rate of 0.01, and the Adam optimizer (**Table S4**). Both models had a test set AUC of about 0.91 (**Table 3**). The threshold *t* that maximized the difference between the true positive rate and the false positive rate was 0.4 for both models.

**Table 3.** Performance of final models with which downstream analysis is conducted. These were the only models for which the test set AUC was calculated. Feature set B is the per-residue ESM2 embeddings; Feature set C is both the per-residue pocket predictions and per-residue ESM2 embeddings concatenated together.

| Model type | Feature set | Train AUC | Val AUC | Test AUC |
| --- | --- | --- | --- | --- |
| NN | B | 0.9807 | 0.9259 | 0.9146 |
| NN | C | 0.9994 | 0.9344 | 0.9132 |

### Assessment of filtered pockets

When visually assessing select experimentally-determined and computationally-predicted protein structures, the *hotpocketNN* model correctly identified the area around the binding pocket and rejected candidate pockets far from the true binding site **(Figure 5**, **Figures S3-S6**). Disordered regions due to low AlphaFold2 confidence were almost entirely predicted as candidate pockets and frequently (but not always) rejected by *hotpocketNN* (**Figure 5, Figures S5-S6**). Filtering results from the *hotpocketNN* model that used Feature Set B (ESM2 embeddings alone) (**Figure 5, Figure S3, Figure S5**) and those from the *hotpocketNN* model that used Feature Set C (ESM2 embeddings and constituent pocket finding method predictions) (**Figure S4, Figure S6**) were similar, with the *hotpocketNN* model that used Feature Set C appearing to identify slightly larger and more pockets than that which used Feature Set B. The most pronounced difference between the two *hotpocketNN* versions was that in the case of low-confidence AlphaFold2 structures, the *hotpocketNN* model that used Feature Set C predicted many more low-confidence disordered regions as being part of pockets (**Figures S7-S8**).

**Figure 5.**
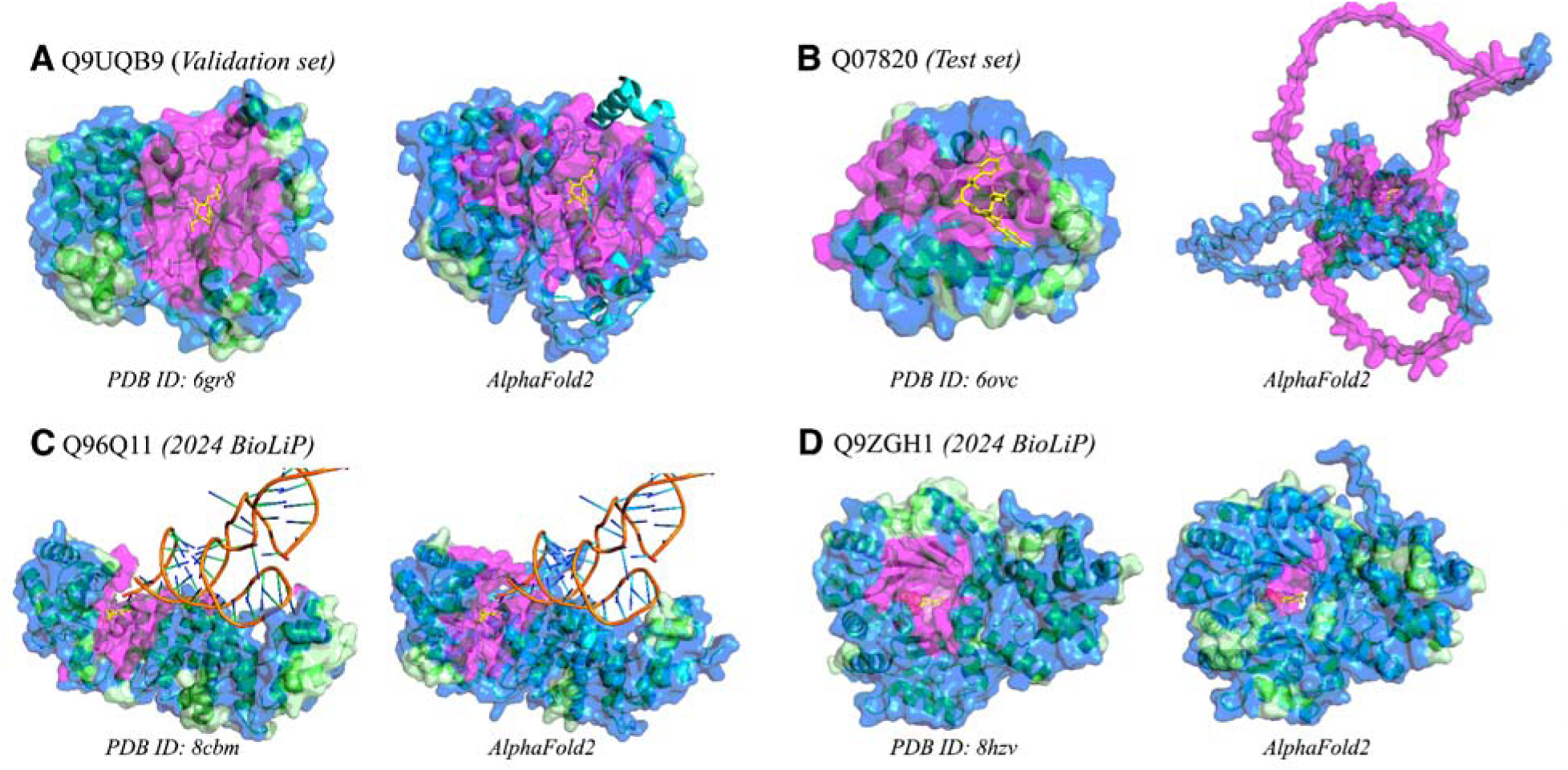
Visualizations of accepted and rejected candidate pockets on select protein structures, using ESM2 embeddings as features for *hotpocketNN* ensembling and filtering method (Feature Set B). Structures are from: **a)** the *hotpocketNN* validation set, **b)** the *hotpocketNN* test set, and **c-d)** the set of human protein structures first released in 2024 that have BioLiP annotations and have low sequence identity to pre-2024 structures. For each protein, both experimentally-determined structures from the PDB (left) and AlphaFold2-predicted structures (right) are shown. The surface of all protein structures is colored as follows: magenta for residues that are part of an accepted candidate pocket accepted by *hotpocketNN*, blue for residues that are part of a candidate pocket but not an accepted candidate pocket, and light green for residues that are not part of any candidate pockets. Biologically-relevant ligands are visualized in the structure as yellow sticks; non-biologically-relevant ligands are omitted. AlphaFold2-predicted structures (green ribbons) are shown aligned with their experimentally-determined PDB counterparts (cyan ribbons). Only the surface of the predicted structure is shown and pocket predictions were made on the predicted structure. The aligned ligands from the experimentally-determined structure are shown in yellow; these ligands are not a part of the AlphaFold2-predicted structures. For each structure, we show only a single biological assembly; for the “2024 BioLiP” structures, we show only a single chain to better see the novel protein-ligand interaction.

The distribution of distances between the pocket center of mass and the closest ligand atom for both versions of *hotpocketNN* had a clear peak at the lowest bin of the histogram, with exponential-like decay and a long tail for greater distances (**Figure 6**). The distributions for each constituent method, with the exception of PocketMiner, also had peaks at the lowest bin of the histogram, though the distributions are bimodal and have a peak near the peak of the reference distribution. Both versions of *hotpocketNN* and all constituent pocket-finding methods were significantly different from the reference distribution according to the Kolmogorov-Smirnov test (p < 1e-88). P2Rank had by far the largest proportion of distances within the lowest bin of the histogram; it accordingly also had the best performance out of all methods for the Human BioLiP dataset per the DCCcriterion (**Table 4**). For this dataset, both versions of *hotpocketNN* performed comparably to top-scoring constituent methods, though did not outperform many. Since P2Rank clearly outperformed all other constituent methods, we used *hotpocketNN* to re-score only candidate pockets from P2Rank; this led to DCCcriterion scores that were almost as high as those of P2Rank. The Feature Set B (ESM2 embeddings only) version of *hotpocketNN* outperformed all other methods, including P2Rank, on the Astex Diverse Set and PoseBusters dataset in terms of DCCcriterion (**Table 4**). We could not plot the distance distribution or calculate the DCCcriterion for the pre-*hotpocketNN* ensemble predictions as there is no way to rank a set of pockets from different methods.

**Figure 6.**
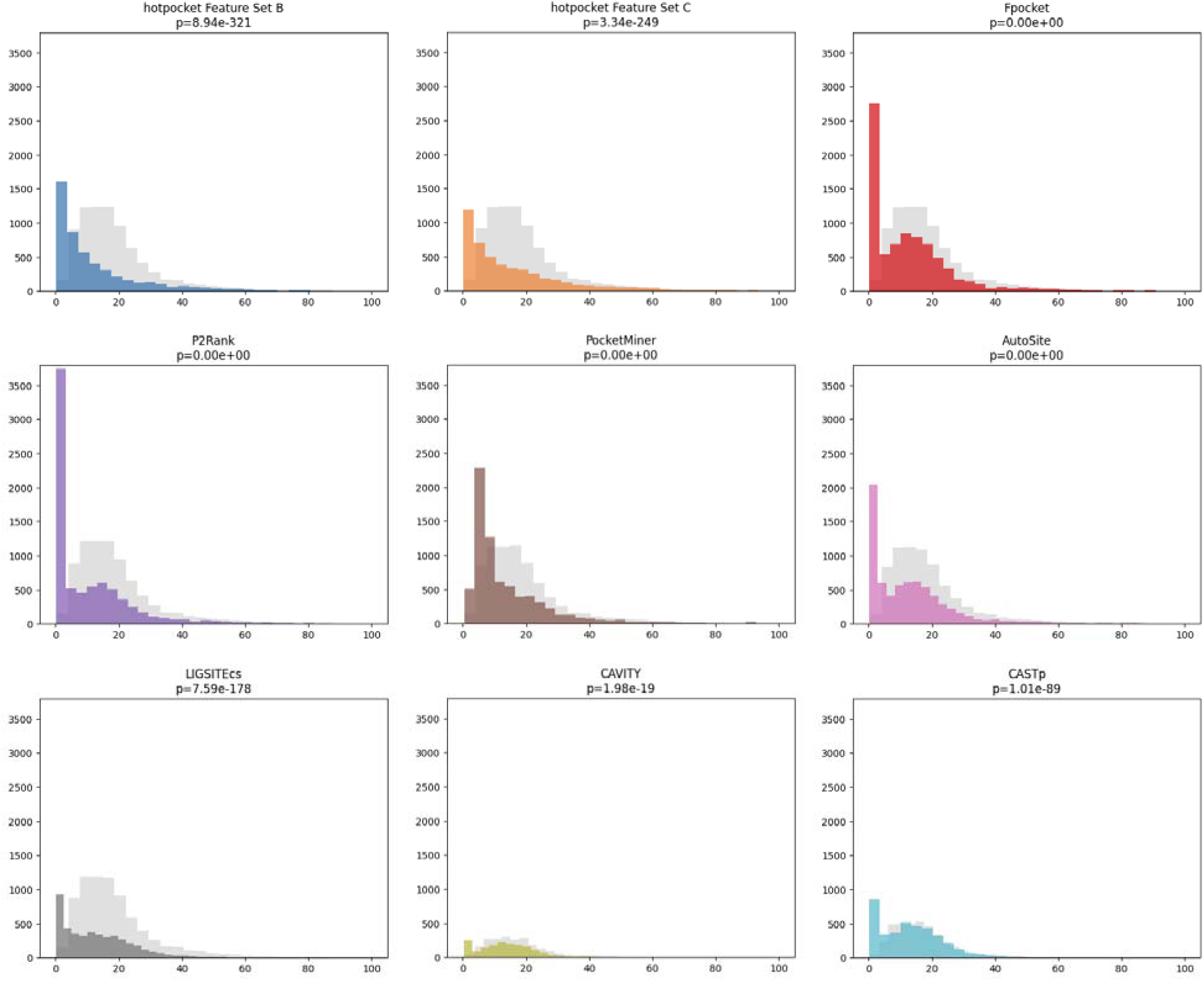
Histograms of the distance between the pocket center of mass and the nearest atom of a relevant ligand, for the top-N pockets of each structure in the Human BioLiP dataset (where N is the number of biologically-relevant ligands in the structure). Both *hotpocketNN* versions and all seven constituent methods are shown separately. Background distributions are shown in light gray. The p-value for the two-sided Kolmogorov-Smirnov test, comparing the distance distribution from each method with its relevant background distribution, is given.

**Table 4.** DCCcriterion using a threshold of 4A for top D_D_ and top D_D_+2 pockets generated by each method, where D_D_ is the number of biologically-relevant ligands in each structure. DCCcriterion is defined as the percentage of pockets that have a center of mass within the threshold distance (here, 4A) of a biologically-relevant ligand; its values range from 0 to 100, with higher values indicating better performance. Any structures present in the *hotpocketNN* neural network training or validation sets were omitted (401 structures removed from Human BioLiP dataset, 1 structure removed from Astex Diverse Set, 0 structures removed from PoseBusters dataset).

|  | Astex Diverse Set |  | PoseBusters |  | Human BioLiP |  |
| --- | --- | --- | --- | --- | --- | --- |
|  | Top N | Top N+2 | Top N | Top N+2 | Top N | Top N+2 |
| *hotpocketNN*, ESM embs | **53.1** | **50.4** | **53.7** | **51.0** | 33.7 | 29.0 |
| *hotpocketNN*, combined | 42.6 | 40.8 | 42.4 | 39.2 | 27.5 | 24.1 |
| *hotpocketNN*, ESM embs, P2Rank only | 41.2 | 32.4 | 47.4 | 37.8 | 46.5 | 36.5 |
| *hotpocketNN*, combined, P2Rank only | 39.9 | 31.7 | 45.3 | 36.9 | 45.0 | 35.8 |
| AutoSite | 35.3 | 27.6 | 34.0 | 25.3 | 33.9 | 26.5 |
| CASTp | 26.7 | 21.1 | - | - | 25.6 | 20.3 |
| CAVITY | 28.6 | 16.7 | - | - | 17.4 | 13.2 |
| Fpocket | 41.3 | 29.8 | 40.7 | 30.2 | 35.8 | 27.2 |
| LIGSITEcs | 45.8 | 34.4 | 38.9 | 30.1 | 25.7 | 21.1 |
| P2Rank | 46.4 | 34.7 | 47.6 | 36.9 | **50.6** | **38.4** |
| PocketMiner | 5.4 | 5.4 | 9.0 | 8.1 | 8.8 | 8.2 |

Following the suggestion of recommendation of Utgés and Barton to increase the DCCcriterion threshold beyond 4A [67], we also calculated the DCCcriterion for all three datasets at thresholds of 8A and 10A (**Tables S5-S6**). As the distance threshold increased, the gap between *hotpocketNN* performance and P2Rank performance increased, *i.e. hotpocketNN* outperformed P2Rank to a greater extent (in the case of Human BioLiP, the gap narrowed, such that P2Rank outperformed *hotpocketNN* to a lesser extent). At the 10A threshold, the *hotpocketNN* re-scoring of P2Rank outperformed all other models on the Human BioLiP task, including P2Rank.

We evaluated per-residue scoring performance with ROC curves and the AUROC (**Figure 7, Figure S9**). When using a threshold of 5A from the relevant ligand to set ground truth labels, performance of both *hotpocketNN* models and the best-performing constituent methods was overall close, with one or both *hotpocketNN* versions beating nearly all constituent methods for each dataset. At a 10A threshold, both *hotpocketNN* versions were consistently the top two methods by AUC. PocketMiner, Fpocket, and P2Rank were consistently among the top models by AUC. CAVITY and LIGSITEcs were consistently the worst models by AUC, likely due to their naive implementation of per-residue scoring. Similar trends appeared in the PRC analysis (**Figure 8, Figure S10**). For the Astex Diverse Set and PoseBusters dataset,

**Figure 7.**
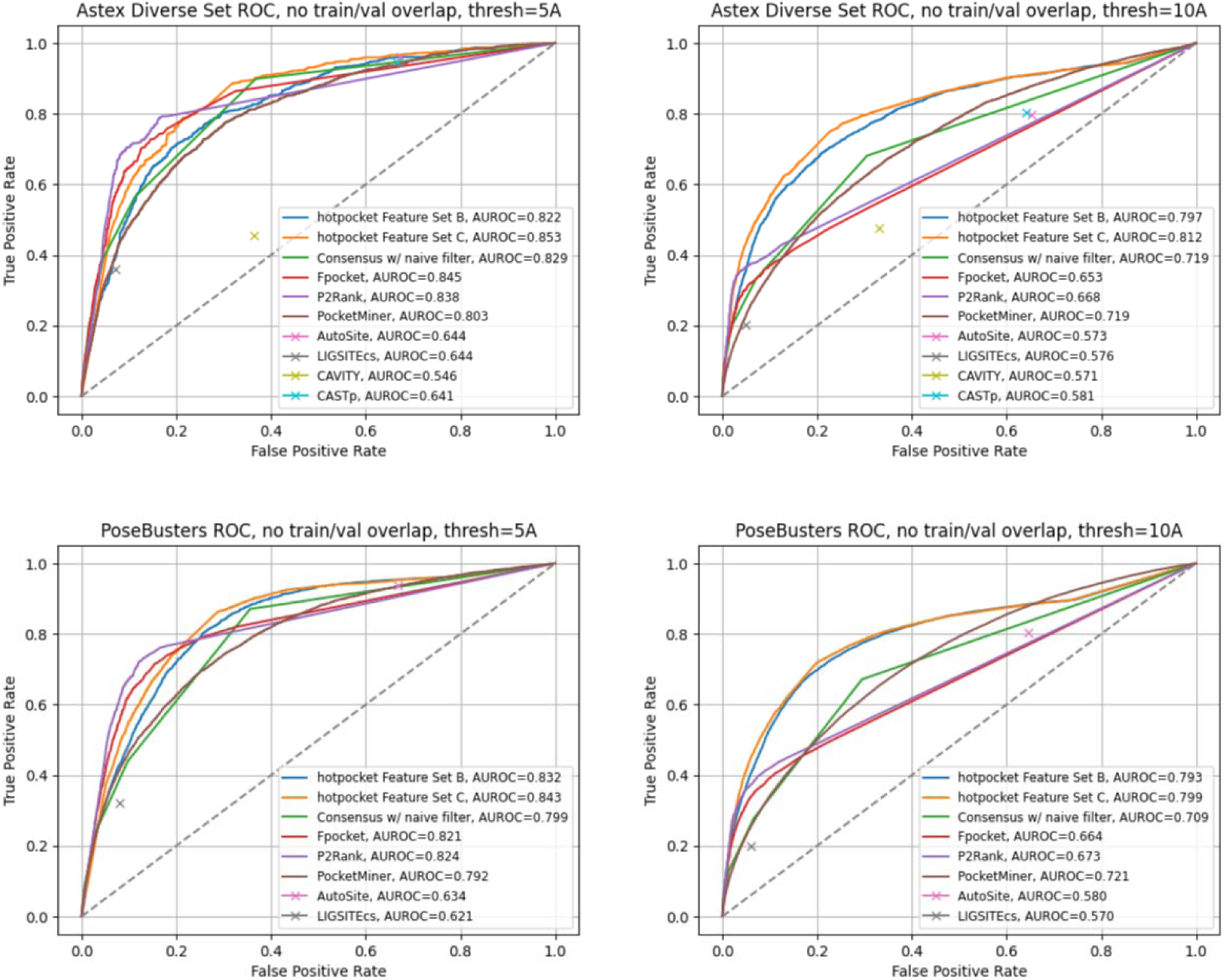
ROC curves and AUROCs for per-residue scoring performance of *hotpocketNN*, naive filter across union of constituent methods, and each constituent pocket-finding method individually across the Astex Diverse Set (top) and PoseBusters dataset (bottom). Each residue was labeled as a binding residue if it was within 5 angstroms (left) or 10 angstroms (right) of a relevant ligand. For the *hotpocketNN* models, Fpocket, P2Rank, and PocketMiner, each residue’s predicted score for being a member of a pocket was the maximum of all predicted pockets of which it was a member. AutoSite, LIGSITEcs, CAVITY, and CASTp do not provide pocket scores; for these methods, each residue’s predicted score for being a member of a pocket was 1 if it is part of any predicted pocket for the structure, or 0 otherwise. One structure from the Astex Diverse Set (and zero structures from the PoseBusters dataset) overlapped with the training/validation sets for *hotpocketNN* and was omitted. CAVITY and CASTp did not have predictions available for any PoseBusters structures. Feature Set B is the per-residue ESM2 embeddings; Feature Set C is both the per-residue pocket predictions and per-residue ESM2 embeddings concatenated together.

**Figure 8.**
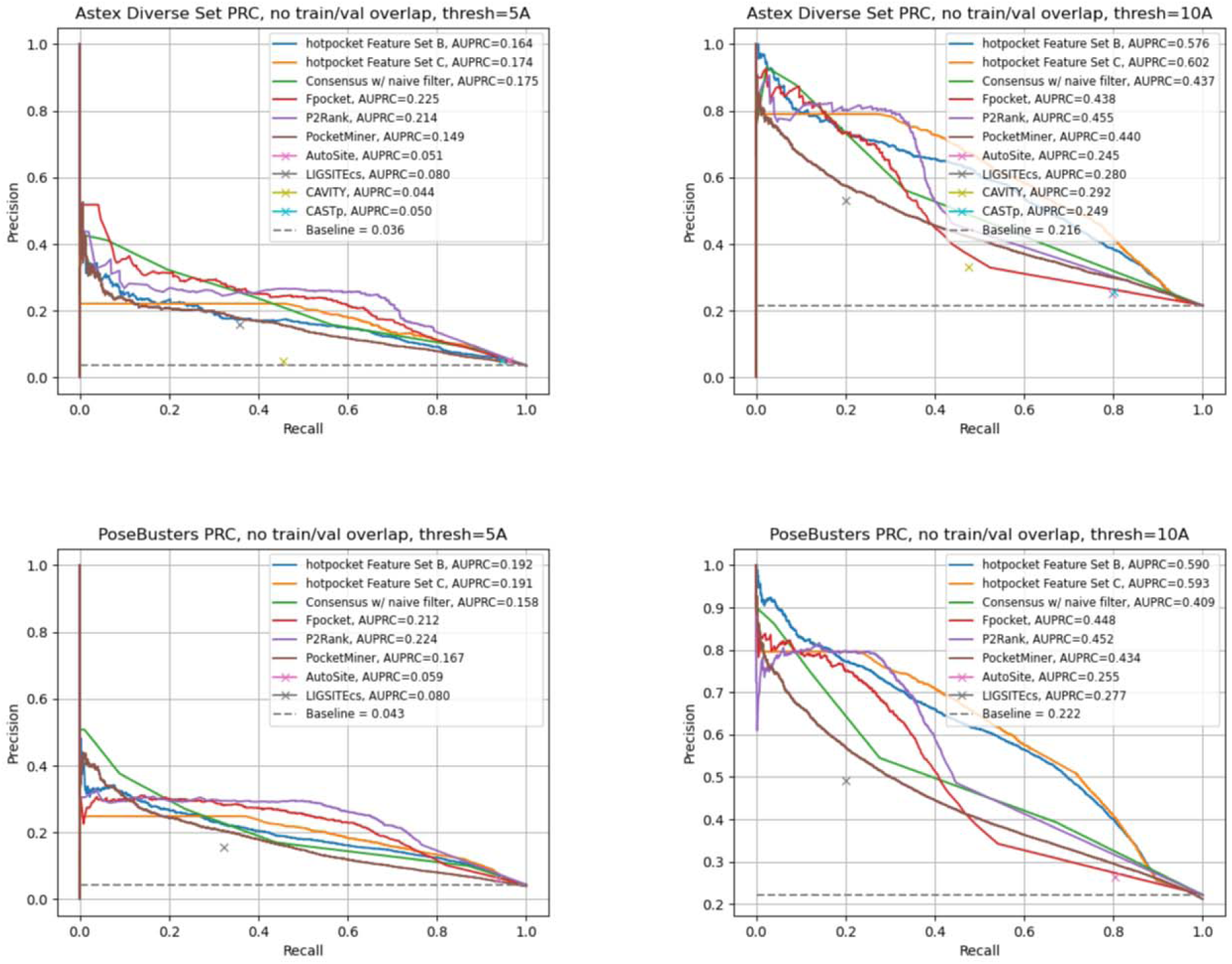
PRC curves and AUPRCs for per-residue scoring performance of *hotpocketNN*, naive filter across union of constituent methods, and each constituent pocket-finding method individually across the Astex Diverse Set (top) and PoseBusters dataset (bottom). Each residue was labeled as a binding residue if it was within 5 angstroms (left) or 10 angstroms (right) of a relevant ligand. For the *hotpocketNN* models, Fpocket, P2Rank, and PocketMiner, each residue’s predicted score for being a member of a pocket was the maximum of all predicted pockets of which it was a member. AutoSite, LIGSITEcs, CAVITY, and CASTp do not provide pocket scores; for these methods, each residue’s predicted score for being a member of a pocket was 1 if it is part of any predicted pocket for the structure, or 0 otherwise.One structure from the Astex Diverse Set (and zero structures from the PoseBusters dataset) overlapped with the training/validation sets for the *hotpocketNN* and was omitted. CAVITY and CASTp did not have predictions available for any PoseBusters structures. Feature Set B is the per-residue ESM2 embeddings; Feature Set C is both the per-residue pocket predictions and per-residue ESM2 embeddings concatenated together.

P2Rank and Fpocket outperformed *hotpocketNN* with a threshold of 5A, but *hotpocketNN* outperformed all constituent methods with a threshold of 10A. For the Human BioLiP dataset, *hotpocketNN* outperformed all constituent methods at both thresholds. In both the ROC and PRC analyses, we observed that the consensus method with the naive filter as described previously was almost never better than the best *hotpocketNN* version and never better than the best constituent method, but was always better than the worst constituent method.

### Preparation of output proteome-wide prediction set

We used both Feature Set B and Feature Set C versions of *hotpocketNN* to make pocket score predictions for the entire human proteome. After filtering out candidate pockets with low *hotpocketNN* scores and low AlphaFold2 confidence scores, we had proteome-wide pocket sets comprising 2,482,870 and 3,475,632 pockets total, respectively for the Feature Set B version (**Table 5**) and the Feature Set C version (**Table S7**) (15.89% and 22.25% of the original predicted pocket set, respectively). While for both versions of *hotpocketNN*, the number of proteins with pockets predicted by each constituent method and the mean/median pockets per structure all decreased, the mean pocket size stayed roughly the same. The different constituent methods had different rates of candidate pocket acceptance, with P2Rank having the highest and CAVITY having the lowest for both *hotpocketNN* versions. The Feature Set C version had higher candidate pocket acceptance rates than the Feature Set B version for all constituent methods. The relative ordering of constituent method candidate pocket acceptance rates was nearly identical between the two versions.

**Table 5.**
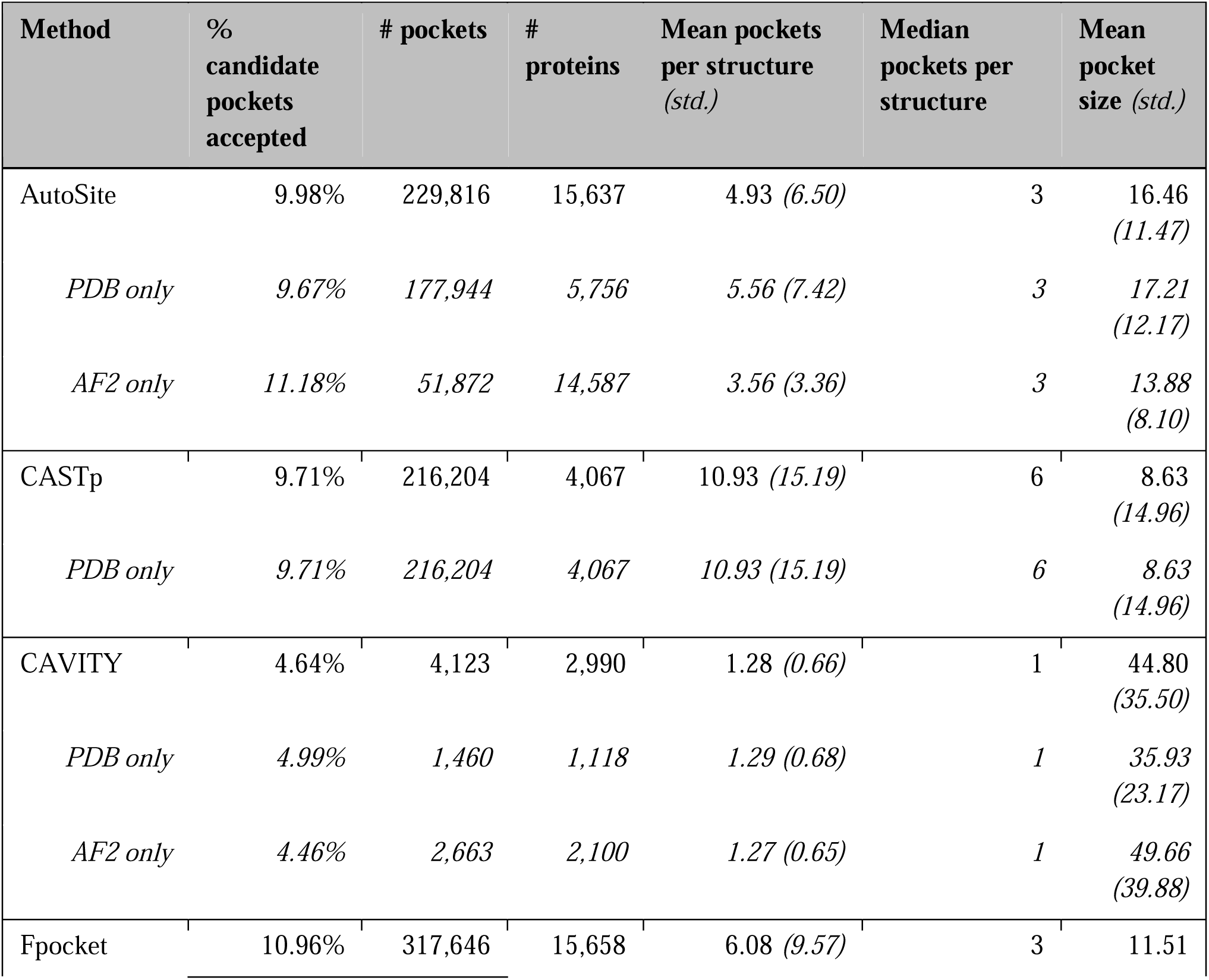
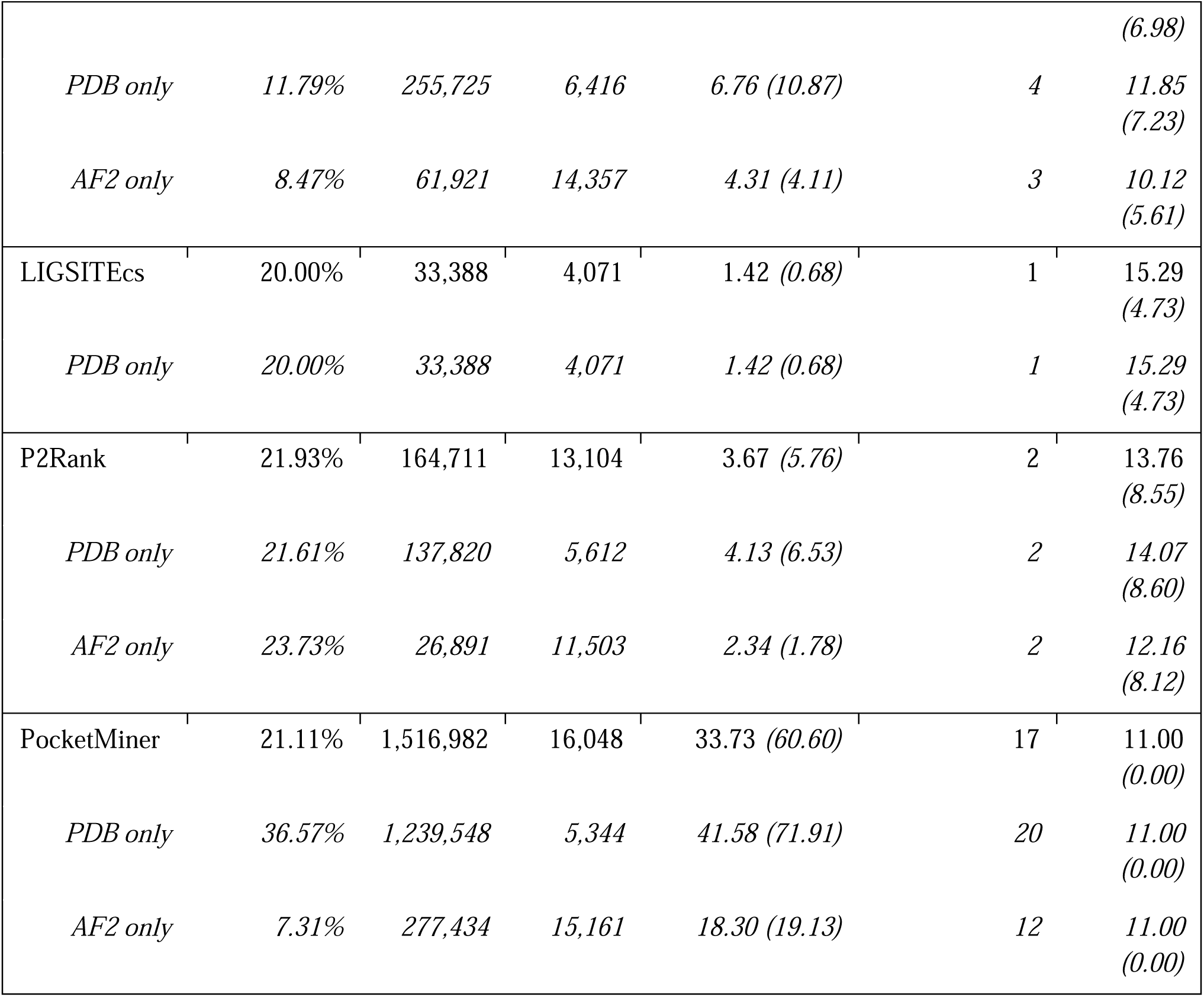
Summary of proteome-wide pockets after passing through the *hotpocketNN* filter (using Feature Set B: ESM2 embeddings only) and the low-confidence AlphaFold2 filter.

### Case studies

#### KRAS

We used the Feature Set B (ESM2 embeddings only) version of *hotpocketNN* to predict pockets on various KRAS PDB structures (**Figure 9**). The *hotpocketNN* model correctly identified both the GDP/GTP binding pocket and the cryptic switch I/II pocket in the 6gj8 structure, which includes a ligand bound to the switch I/II pocket (**Figure 9a**, **Table S8**). The *hotpocketNN* model failed to identify the switch I/II pocket on 3gft, which is GTP-bound (**Figure 9b**, **Table S8**). The remaining four KRAS structures we analyzed were all GDP-bound with no other ligand present. For two of them (5uk9 and 6quu), *hotpocketNN* correctly identified the switch I/II pocket despite no ligand being bound to it (**Figure 9c**, **Figure 9e**, **Table S8**). The other two structures (6bp1 and 7lz5) had about half of the switch I/II pocket highlighted by *hotpocketNN* (**Figure 9e**, **Figure 9f**, **Table S8**).

**Figure 9.**
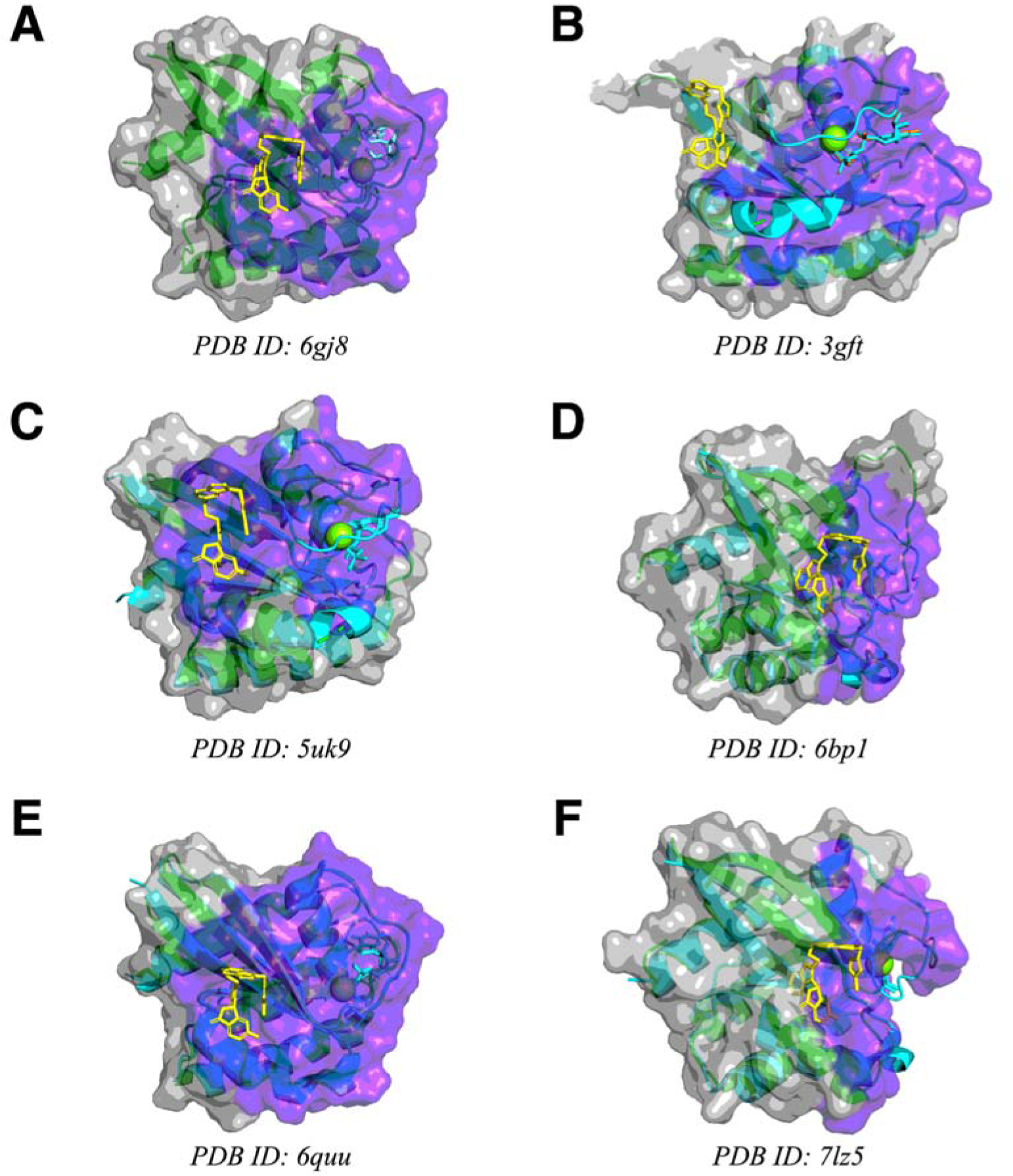
*hotpocketNN* predicted pockets (purple highlight) for various KRAS structures (green ribbons and gray surface): **a)** 6gj8 (GDP-bound), **b)** 3gft (GTP-bound); **c)** 5uk9 (GDP-bound); **d)** 6bp1 (GDP-bound); **e)** 6quu (GDP-bound); **f)** 7lz5 (GDP-bound). In subfigures **b-f**, the 6gj8 structure (cyan ribbons, no surface) is shown aligned with the main structure. The cryptic pocket ligand from the 6gj8 structure is shown in yellow sticks, GDP analogues are shown in cyan sticks, and GTP analogues are shown in orange sticks.

#### Mu opioid receptor

We used the Feature Set B (ESM2 embeddings only) version of *hotpocketNN* to predict pockets on various mOR PDB structures (**Figure 10**). For all three structures, *hotpocketNN* recovered both the orthosteric pocket (recall ranging from 0.67 to 0.93) and the “inside” allosteric pocket (recall ranging from 0.47 to 0.74) (**Figure 10**, **Table S9**). The *hotpocketNN* model recovered the “outside” allosteric pocket in one structure (8ef5), but not the other two (**Figure 10**, **Table S9**).

**Figure 10.**
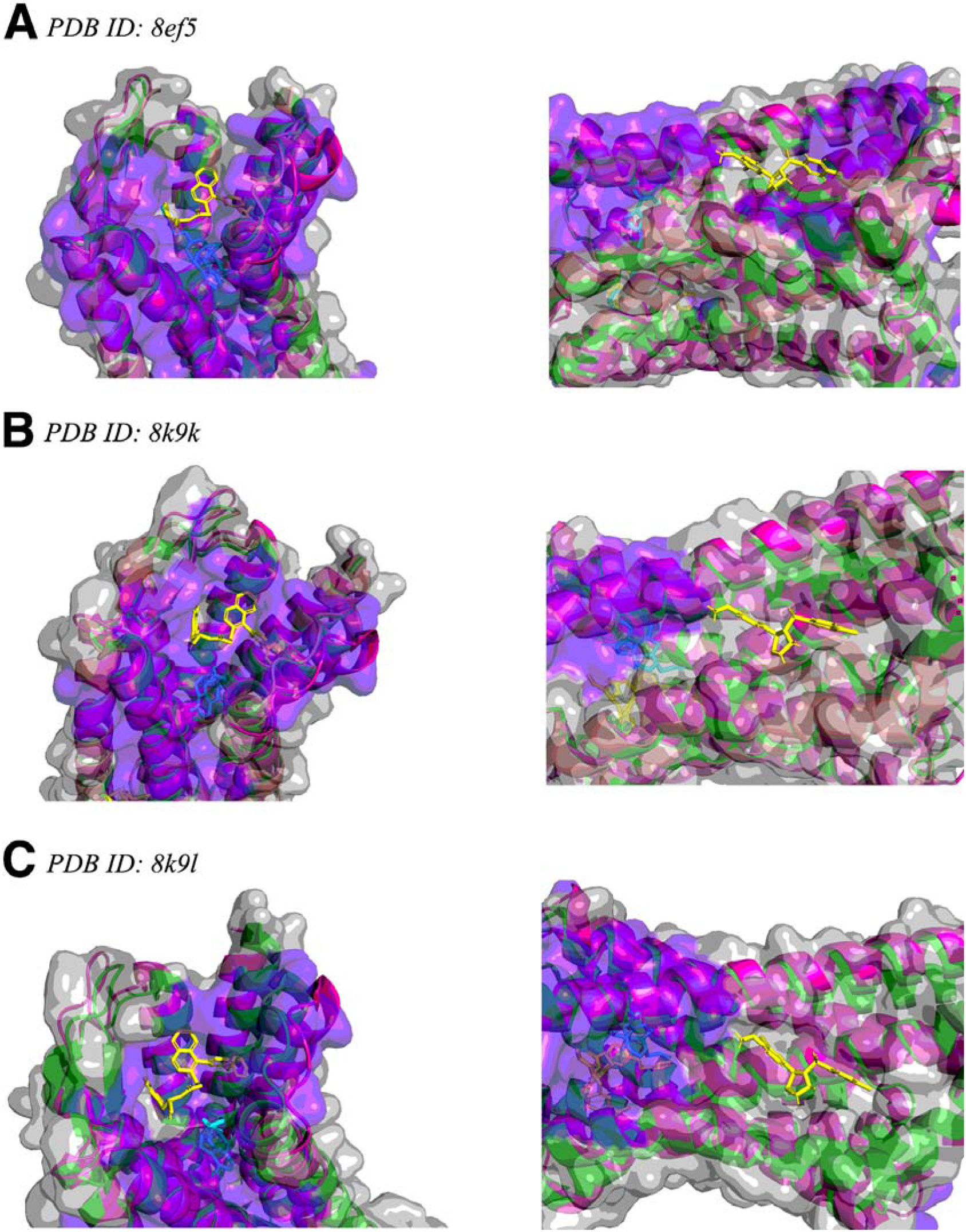
*hotpocketNN* predicted pockets (purple highlight) for various mu opioid receptor structures (green ribbons and gray surface): **a)** 8ef5 (with fentanyl as ligand), **b)** 8k9k (without ligands); **c)** 8k9l (with “outside” allosteric ligand). Left: “inside” allosteric pocket. Right: “outside” allosteric pocket. In all subfigures, the 9bjk structure (pink ribbons, no surface) is shown aligned with the main structure. In subfigures **a** and **b**, the 8k9l structure (salmon ribbons, no surface) is shown aligned with the main structure. The allosteric ligands from the 8k9l and 9bjk structures are shown in yellow sticks and the orthosteric ligands (fentanyl in 8ef5, naloxone in 9bjk) are shown in cyan sticks.

## Discussion

We have introduced a novel dataset, HOTPocket, which encompasses harmonized binding pocket predictions from an ensemble of prediction methods across the whole human proteome. While a simple intersection of different methods’ predictions appeared to work well sometimes (**Figure 3**), we observed that performance did not exceed that of Fpocket or P2Rank individually upon larger scale evaluation (**Figure 4**). We created *hotpocketNN*, a neural network model that uses ESM2 embeddings of candidate pocket residues to generate a pocket likelihood score. Scoring candidate pockets with *hotpocketNN* allows for direct comparison and ranking between pockets produced by different pocket-finding methods. We also retain the name of the original pocket-finding method that produced each accepted candidate pocket to enhance interpretability (*e.g.* one knows which pocket-finding method originally generated the pocket and can then investigate method-specific metrics) and to allow for downstream filtering according to use case (*e.g.* only considering pockets from certain pocket-finding methods to yield pockets with desired properties). The *hotpocketNN* method was able to recover known binding pockets, outperformed constituent pocket-finding methods on the Astex Diverse Set and PoseBusters dataset with respect to DCCcriterion, and consistently performed well with respect to per-residue AUROC and AUPRC.

We tested three different featurization schemes for *hotpocketNN*. Both schemes that included the ESM2 embeddings performed well; Feature Set B included only these as features, while Feature Set C also included per-residue scores from each of the constituent pocket-finding methods. ESM2 embeddings likely worked well as features because they leverage latent patterns observed in millions of proteins on which the ESM2 model was trained, and are simply higher dimension and more sophisticated than the Feature Set A scheme of encoding per-residue predictions from the seven constituent pocket-finding methods. As neither *hotpocketNN* version clearly outperformed the other, we have provided assessments of both and have also used both to create two HOTPocket datasets for public use. However, we expect that Feature Set B (ESM2 embeddings only) may be more useful for predictions on novel protein structures that may not have predictions available from the constituent pocket-finding methods.

Previous research has found that using ESM2 embeddings, and embeddings from other protein language models, as features for protein function prediction and structural annotation models yields excellent performance [68], [69], [70], [71]. Some existing methods have used ESM2 embeddings in a manner very similar to *hotpocketNN* for binding site prediction. Wang *et al.* used ESM2 embeddings to predict binding site probability scores over the whole protein [72]; *hotpocketNN* is distinct in that it is also an ensemble model, lending more interpretability and confidence. Additionally, we distinguish our work by providing a dataset of precomputed predictions. Škrhák *et al.* also introduced a model with very similar architecture to *hotpocketNN*, but theirs is only focused on cryptic pockets [73].

We observed that the pockets predicted by *hotpocketNN* tended to be relatively large. Despite larger size, these predicted pockets still tended to center on the ligand of interest (**Figure 6**). Highlighting a larger pocket may not be problematic as ligands can vary in size. The per-residue AUROC and AUPRC assessment required selecting a threshold radius around the ligand of interest for which to define ground truth pocket residues. At a threshold of 5A, some of the outer residues in the pocket predicted by *hotpocketNN* may erroneously be labeled as false positives. These residues would be labeled as true positives with a threshold of 10A. We observed that *hotpocketNN* performance was close to that of the top-performing constituent methods at a threshold of 5A, and surpassed all constituent models at a threshold of 10A (**Figures 7-8**). This shows that comparing methods with the per-residue ROC/PRC metric is difficult as it is dependent on pocket size, with smaller predicted pockets leading to better performance with smaller thresholds and larger predicted pockets leading to better performance with larger thresholds. A related consideration is raised by Utgés and Barton [67]: because many different ligands of different sizes could bind a single binding site, a DCCcriterion threshold of 4A may be too conservative. When we increased the DCCcriterion threshold, we found that *hotpocketNN* outperformed constituent models to a greater extent (when it originally was the best method) or was outperformed by constituent models to a lesser extent (when it originally was not the best method).

We observed that P2Rank and Fpocket outperformed *hotpocketNN* in the DCCcriterion analysis on the Human BioLiP dataset, but not on the Astex Diverse Set or PoseBusters dataset. This is likely due to overlap between the Human BioLiP dataset and the P2Rank and Fpocket training datasets. The Astex Diverse Set is commonly used as a held-out validation or test set, and the PoseBusters dataset was constructed so as to only include structures that were released after the training of both of these methods, making them fairer datasets for evaluation.

Computationally-predicted protein structures are now as accurate as experimentally-determined protein structures for many proteins, with a few notable exceptions [74], [75], [76], [77]. It is possible to use computationally-predicted structures for drug discovery when a high-quality experimentally-determined structure does not yet exist. For this reason, we evaluated *hotpocketNN* both on established benchmark datasets of experimentally-determined structures and on AlphaFold2-predicted structures. We found that *hotpocketNN* could recover known binding pockets from AlphaFold2-predicted structures, even when the predicted structure was generated before the structure with the known binding pocket was experimentally resolved (**Figure 5**). The *hotpocketNN* model generally discarded candidate pockets predicted on low-confidence regions; we also filtered out low-confidence pocket regions when compiling the HOTPocket dataset.

When selecting pocket-finding methods to include as in our ensemble model, it became apparent that we could not include many models with reported excellent performance because their source code or precomputed predictions were not available (*e.g.* if the method was only available as a web server for small-scale use) or their code was no longer usable due to lack of maintenance. This underscores the importance of making code and data available and regular website and package maintenance. Motivated by this, we have made the HOTPocket dataset freely available and the *hotpocketNN* method open source.

Our assessment of novel BioLiP interactions discovered in 2024 showed that *hotpocketNN* can recover recently-discovered pockets not present in its training data, demonstrating real-world utility. Similarly, in the KRAS case study, *hotpocketNN* fully or partially recovered a recently-discovered druggable pocket on all structures – including structures from before the pocket was known – except for a structure in which KRAS was in the GTP-bound state. It is possible that this pocket is not present or less accessible in the GTP-bound state. The mOR case study was less conclusive, with one allosteric pocket being easy to predict due to its proximity to the orthosteric pocket (and presence inside the helical bundle) and the other only being recovered in one of three structures. It is possible that the “outside” allosteric pocket is weaker, leading to it not always being recovered. When searching for weak pockets, an end user may elect to lower the score threshold. Supplementing *hotpocketNN* analysis with molecular dynamics may also yield further insight.

One limitation of the HOTPocket dataset is that it may eventually become outdated as constituent pocket-finding methods get updated and more structures of human proteins are deposited in the PDB. In between updates to HOTPocket, end users may still use the *hotpocketNN* code to generate predictions on new structures or use the provided code base to update their version of HOTPocket locally. Inclusion of a new pocket-finding method only requires the ability to run the new method over HOTPocket’s assembled proteome structures; the Feature Set B version of *hotpocketNN* can be run on any predicted pocket regardless of method of origin. A limitation of the *hotpocketNN* method is that there are some cases in which one constituent method (*e.g.* P2Rank) is clearly superior and the other methods add noise that decreases *hotpocketNN* performance. In those cases, end users may choose to only include candidate pockets from the top-performing constituent method, which we have shown boosts performance of *hotpocketNN*.

## List of abbreviations

AF2: AlphaFold2
API: Application programming interface
AUC: See “AUROC”
AUPRC: Area under the precision-recall curve
AUROC: Area under the receiver operating characteristic
CNN: Convolutional neural network
GDP: Guanosine diphosphate
GPCR: G-protein coupled receptor
GTP: Guanosine triphosphate
HOTPocket: The Human Omnibus of Targetable Pockets
ML: Machine learning
mOR: Mu opioid receptor
NN: Neural network
PDB: Protein Data Bank
PRC: Precision-recall curve
ROC: Receiver operating characteristic
TTD: Therapeutic Target Database

## Declarations

### Availability of data and materials

Code and data are available at github.com/Helix-Research-Lab/HOTPocket.

## Authors’ contributions

KAC: Conceptualization; Methodology; Software; Validation; Investigation; Data Curation; Writing - Original Draft; Writing - Review & Editing; Visualization; Funding acquisition

RBA: Conceptualization; Resources; Writing - Review & Editing; Supervision; Project administration; Funding acquisition

## Acknowledgements

We thank Alexander Derry, Alp Tartici, Gowri Nayar, and the Helix Group at large for insightful discussions that improved the work. Most of the computing for this project was performed on the Sherlock cluster. We would like to thank Stanford University and the Stanford Research Computing Center for providing computational resources and support that contributed to these research results.

## Supplement

**Text S1.** List of excluded BioLiP ligand identifiers for “druglike ligand” filter to include only organic small molecules.

dna, rna, peptide, ACT, CA, CO, CO3, CU, FE, K, MG, MN, PO4, SO3, SO4, ZN

**Text S2.** Naive pocket filtering criteria. <u>AutoSite</u>

Built-in score: None provided besides ranked pocket order. Filter: Include only first five pockets.

### CASTp

Built-in score: None provided besides ranked pocket order. Filter: Include only first five pockets.

### CavitySpace

Built-in score: Categorical pocket druggability classes.

Filter: Include only pockets labeled with “Strong” druggability.

### Fpocket

Built-in score: Pocket druggability score ranging from 0 to 1. Filter: Include only pockets with druggability score of at least 0.5.

### LIGSITEcs

Built-in score: None provided besides ranked pocket order. Filter: Include only first five pockets.

### P2Rank

Built-in score: Pocket probability scores ranging from 0 to 1. Filter: Include only pockets with probability score of at least 0.5.

### PocketMiner

Built-in score: Per-residue pocket likelihood score ranging from 0 to 1; aggregated score cutoff of 0.7 already accounted for in method for generating predicted pockets.

Filter: Include only pockets with aggregated pocket likelihood score of at least 0.8.

---

**Table S1.** Non-comprehensive list of pocket-finding methods from literature review conducted in 2023. See attached table.

**Table S2.**
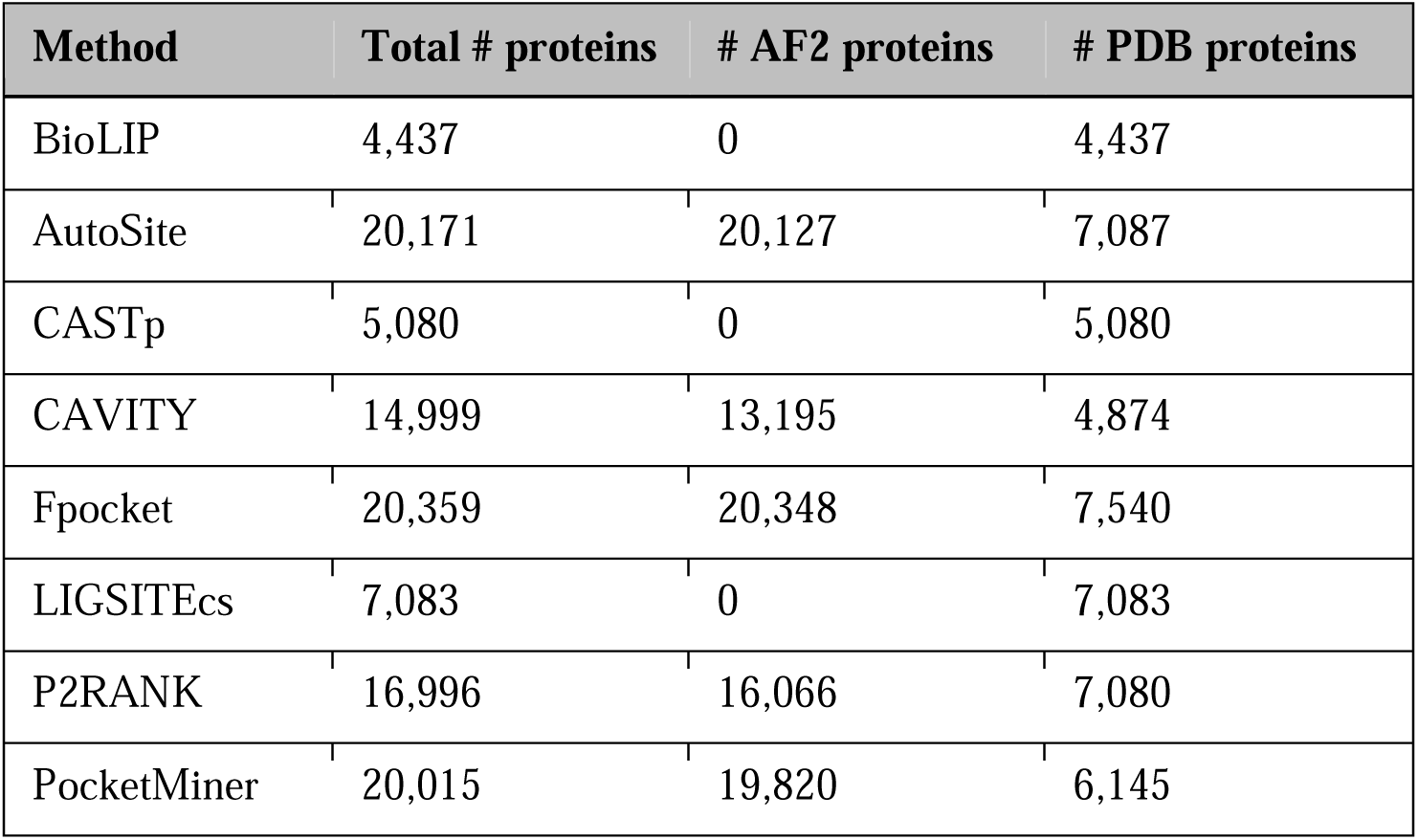
Breakdown of proteins with experimentally-determined protein structures (“PDB”) and proteins with computationally-predicted protein structures (“AF2”) for which pocket annotations and predictions were generated.

**Table S3.**
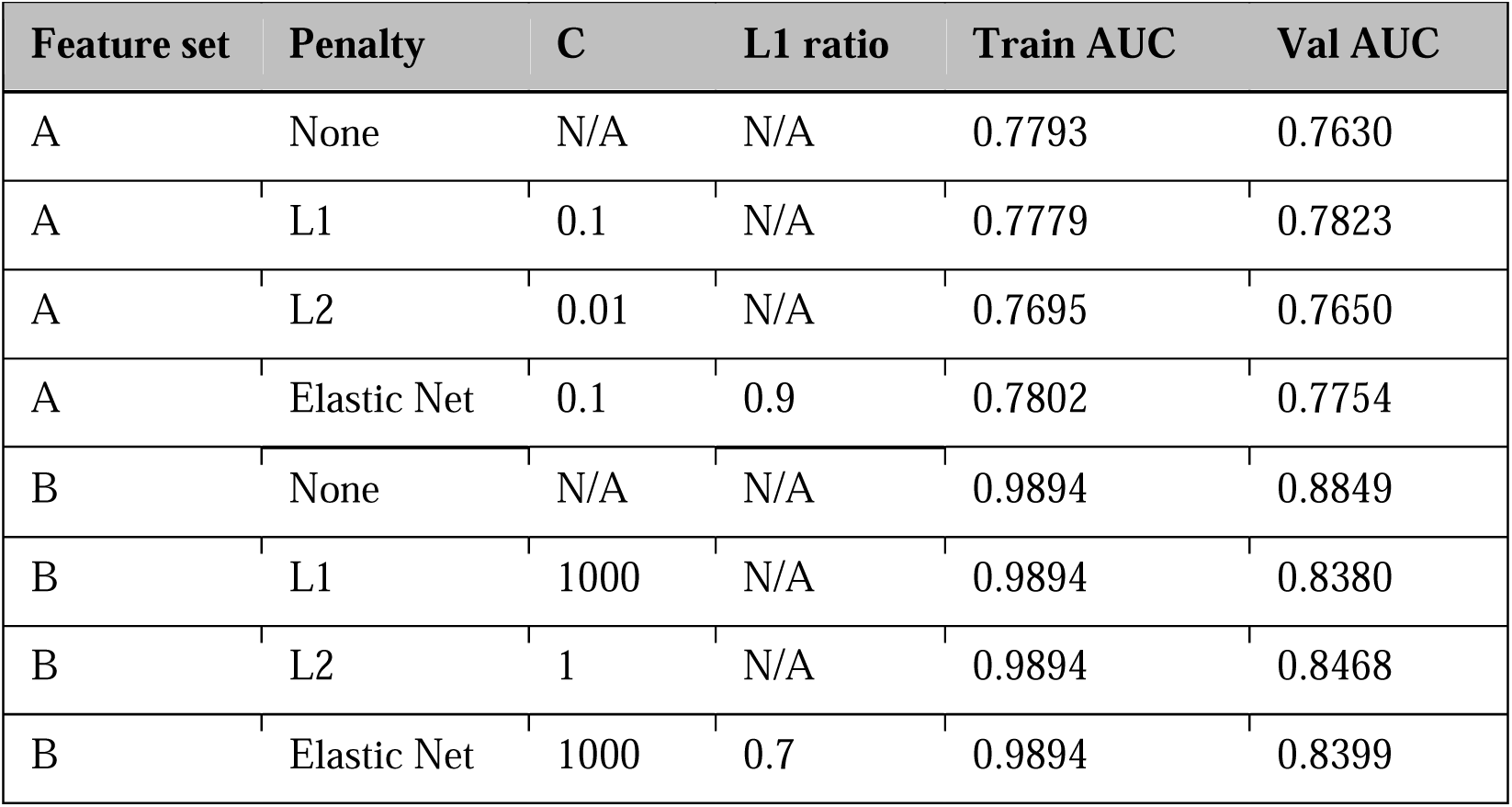
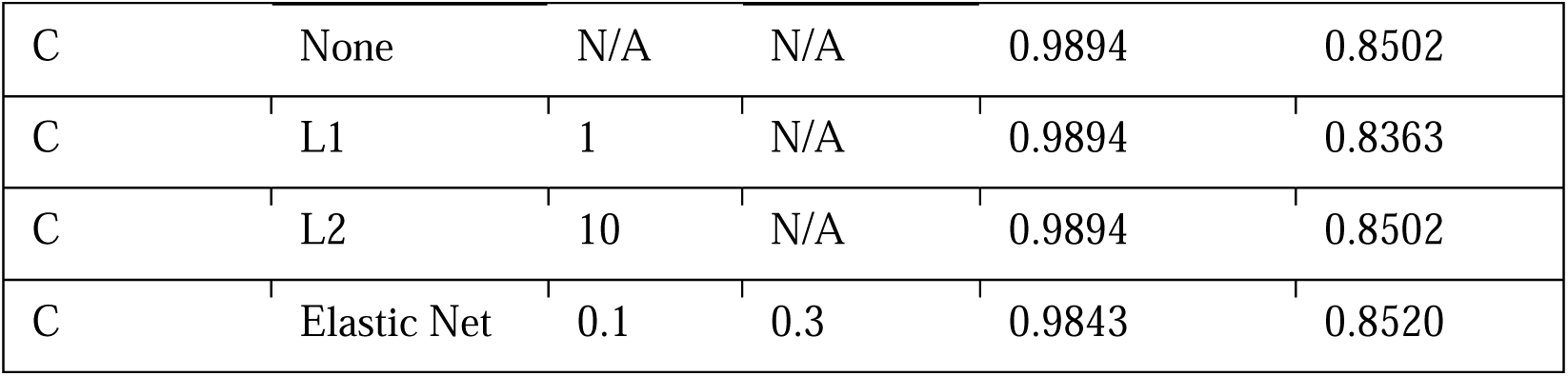
Logistic regression hyperparameter tuning. For each combination of feature set and logistic regression penalty, the C value, L1 ratio, training set AUC, and validation set AUC are shown for the model with the best validation AUC out of all hyperparameter combinations assessed. Models were trained until convergence or a maximum of 500 iterations. There was no variation in performance across different random seeds. Feature set A is the per-residue pocket predictions from each of the constituent methods; Feature set B is the per-residue ESM2 embeddings; Feature set C is both the per-residue pocket predictions and per-residue ESM2 embeddings concatenated together.

**Table S4.**
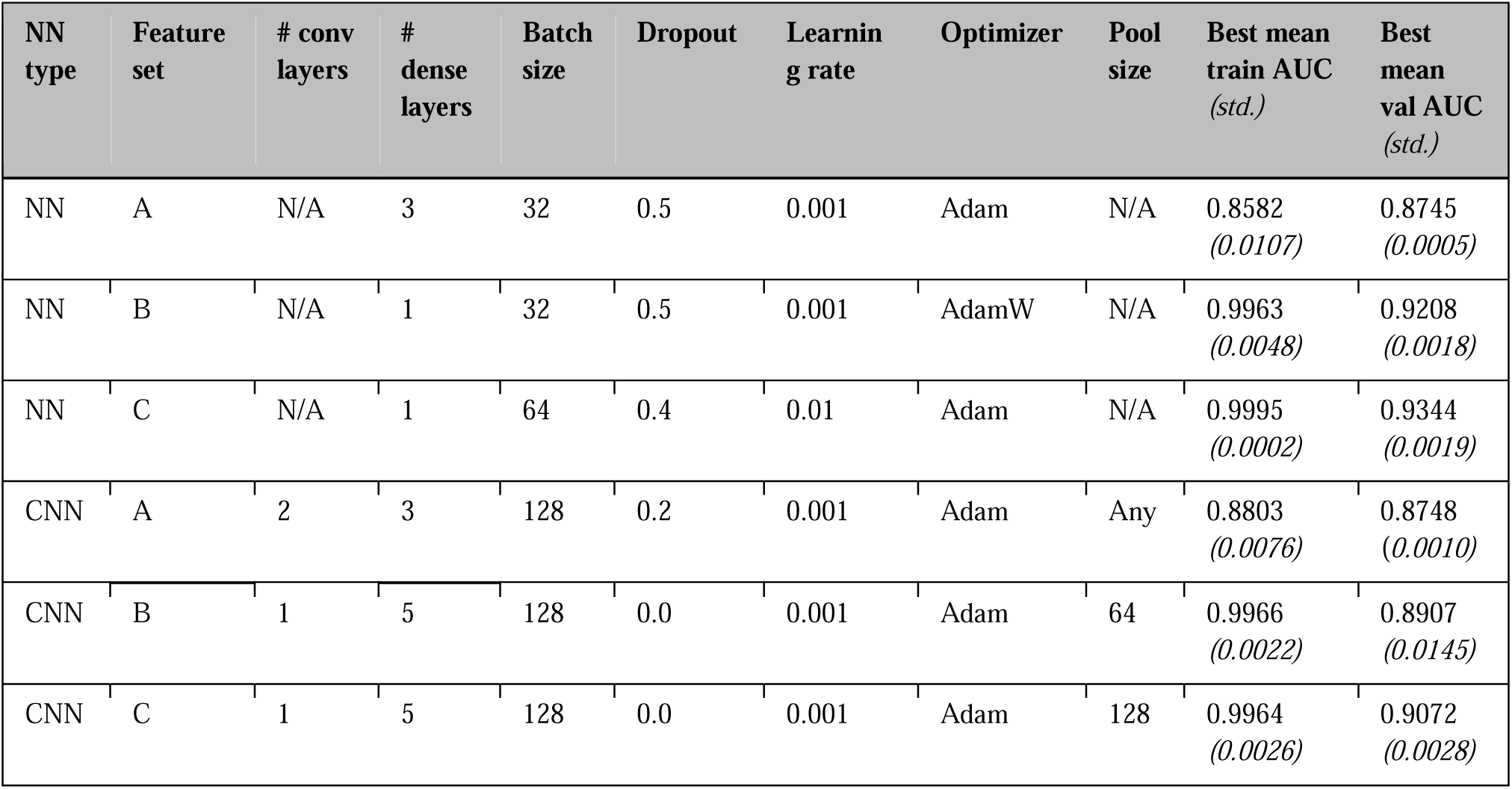
Neural network (NN and CNN) hyperparameter tuning. For each combination of feature set and neural network type (feedforward NN or convolutional NN), the number of convolutional layers, number of feedforward layers, batchsize, dropout probability, learning rate, optimizer, pooling size, training set AUC, and validation set AUC are shown for the model with the best validation AUC out of all hyperparameter combinations assessed. The AUCs shown are averaged over three identical models trained with different random seeds and are from the epoch with the highest validation AUC achieved in xx epochs. Feature set A is the per-residue pocket predictions from each of the constituent methods; Feature set B is the per-residue ESM2 embeddings; Feature set C is both the per-residue pocket predictions and per-residue ESM2 embeddings concatenated together.

**Table S5.**
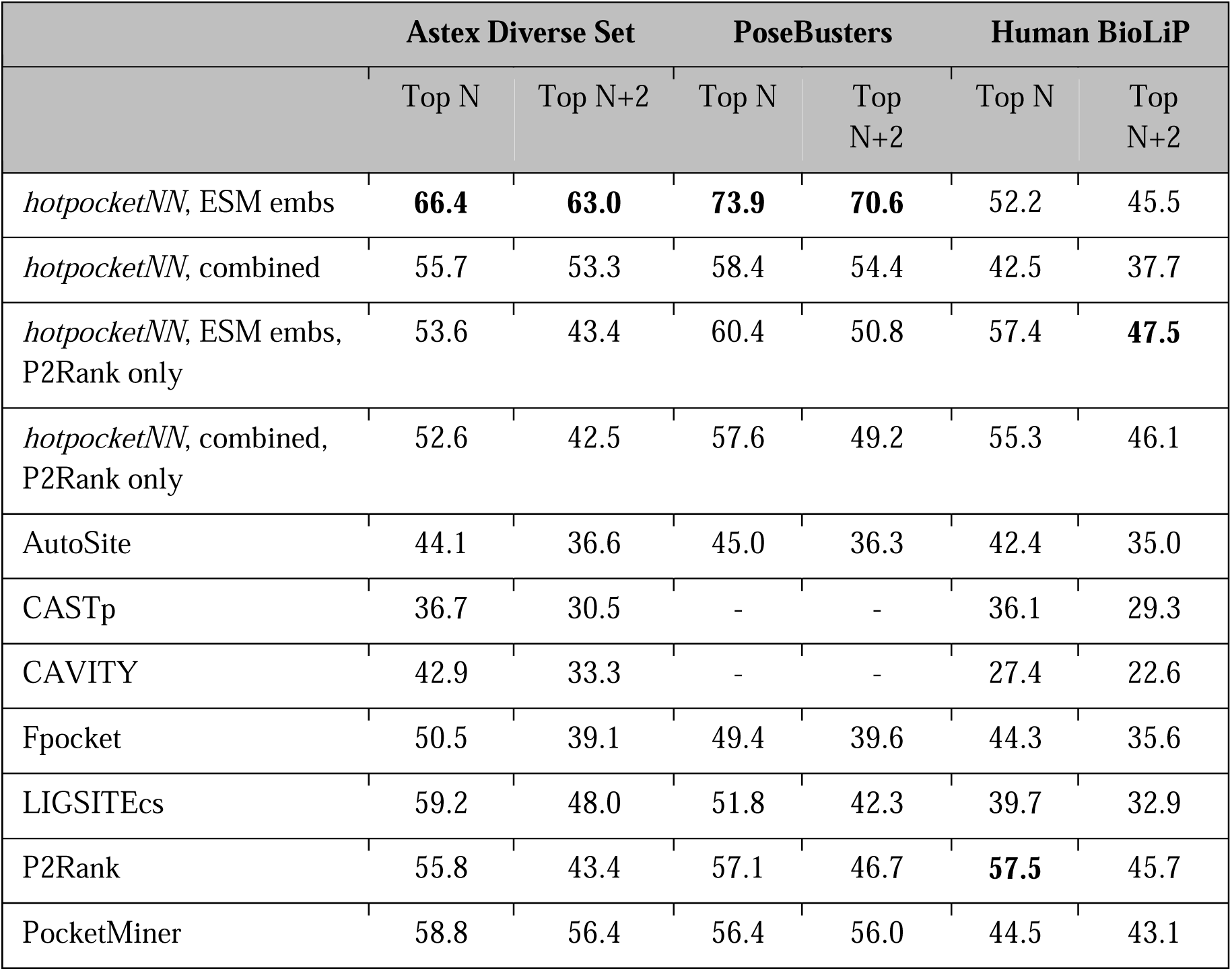
DCCcriterion using a threshold of 8A for top LJ_D_ and top LJ_D_+2 pockets generated by each method, where LJ_D_ is the number of biologically-relevant ligands in each structure. DCCcriterion is defined as the percentage of pockets that have a center of mass within the threshold distance (here, 8A) of a biologically-relevant ligand; its values range from 0 to 100, with higher values indicating better performance. Any structures present in the *hotpocketNN* training or validation sets were omitted (401 structures removed from Human BioLiP dataset, 1 structure removed from Astex Diverse Set, 0 structures removed from PoseBusters dataset).

|  | Astex Diverse Set |  | PoseBusters |  | Human BioLiP |  |
| --- | --- | --- | --- | --- | --- | --- |
|  | Top N | Top N+2 | Top N | Top N+2 | Top N | Top N+2 |
| *hotpocketNN*, ESM embs | **66.4** | **63.0** | **73.9** | **70.6** | 52.2 | 45.5 |
| *hotpocketNN*, combined | 55.7 | 53.3 | 58.4 | 54.4 | 42.5 | 37.7 |
| *hotpocketNN*, ESM embs, P2Rank only | 53.6 | 43.4 | 60.4 | 50.8 | 57.4 | **47.5** |
| *hotpocketNN*, combined, P2Rank only | 52.6 | 42.5 | 57.6 | 49.2 | 55.3 | 46.1 |
| AutoSite | 44.1 | 36.6 | 45.0 | 36.3 | 42.4 | 35.0 |
| CASTp | 36.7 | 30.5 | - | - | 36.1 | 29.3 |
| CAVITY | 42.9 | 33.3 | - | - | 27.4 | 22.6 |
| Fpocket | 50.5 | 39.1 | 49.4 | 39.6 | 44.3 | 35.6 |
| LIGSITEcs | 59.2 | 48.0 | 51.8 | 42.3 | 39.7 | 32.9 |
| P2Rank | 55.8 | 43.4 | 57.1 | 46.7 | **57.5** | 45.7 |
| PocketMiner | 58.8 | 56.4 | 56.4 | 56.0 | 44.5 | 43.1 |

**Table S6.**
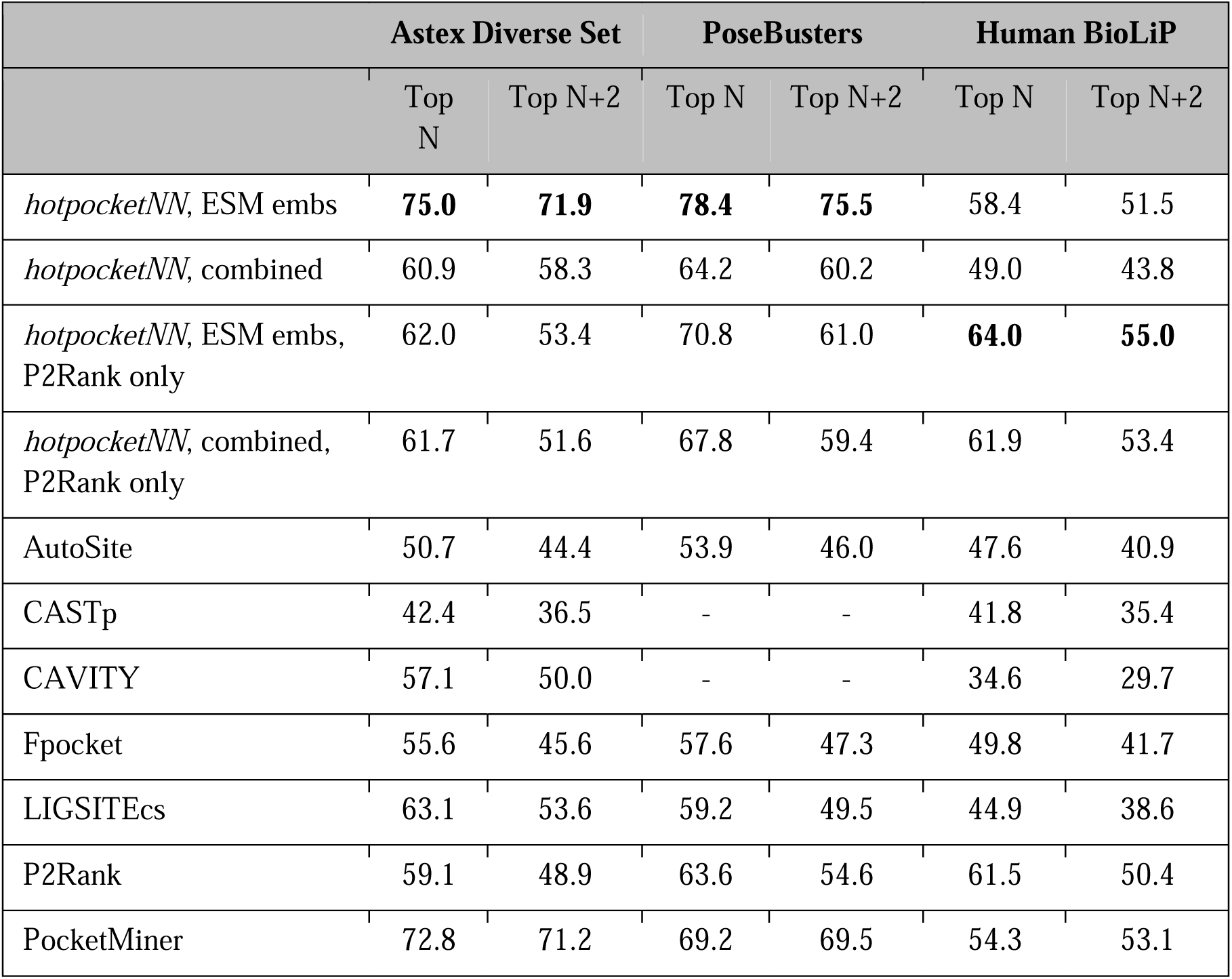
DCCcriterion using a threshold of 10A for top LJ_D_ and top LJ_D_+2 pockets generated by each method, where LJ_D_ is the number of biologically-relevant ligands in each structure. DCCcriterion is defined as the percentage of pockets that have a center of mass within the threshold distance (here, 10A) of a biologically-relevant ligand; its values range from 0 to 100, with higher values indicating better performance. Any structures present in the *hotpocketNN* training or validation sets were omitted (401 structures removed from Human BioLiP dataset, 1 structure removed from Astex Diverse Set, 0 structures removed from PoseBusters dataset).

**Table S7.**
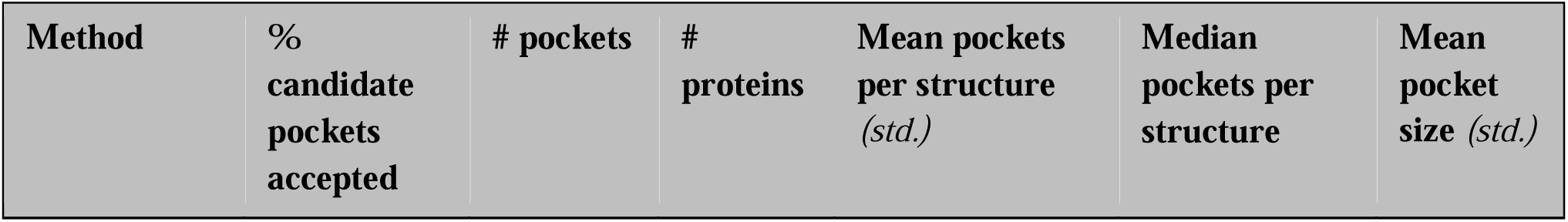
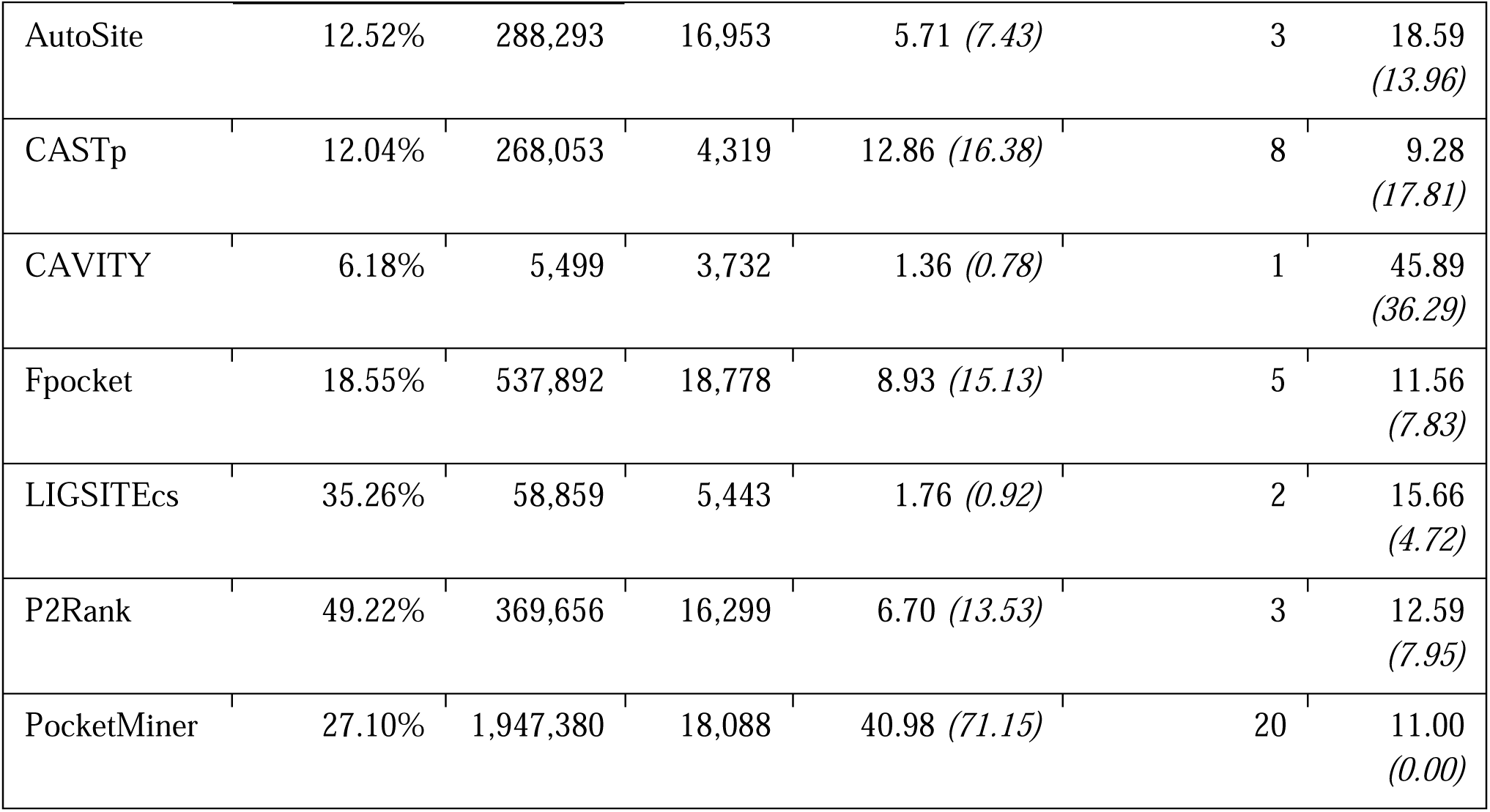
Summary of proteome-wide pockets after passing through the *hotpocketNN* filter (using Feature Set C: ESM2 embeddings and per-residue constituent method predictions) and the low-confidence AlphaFold2 filter.

**Table S8.**
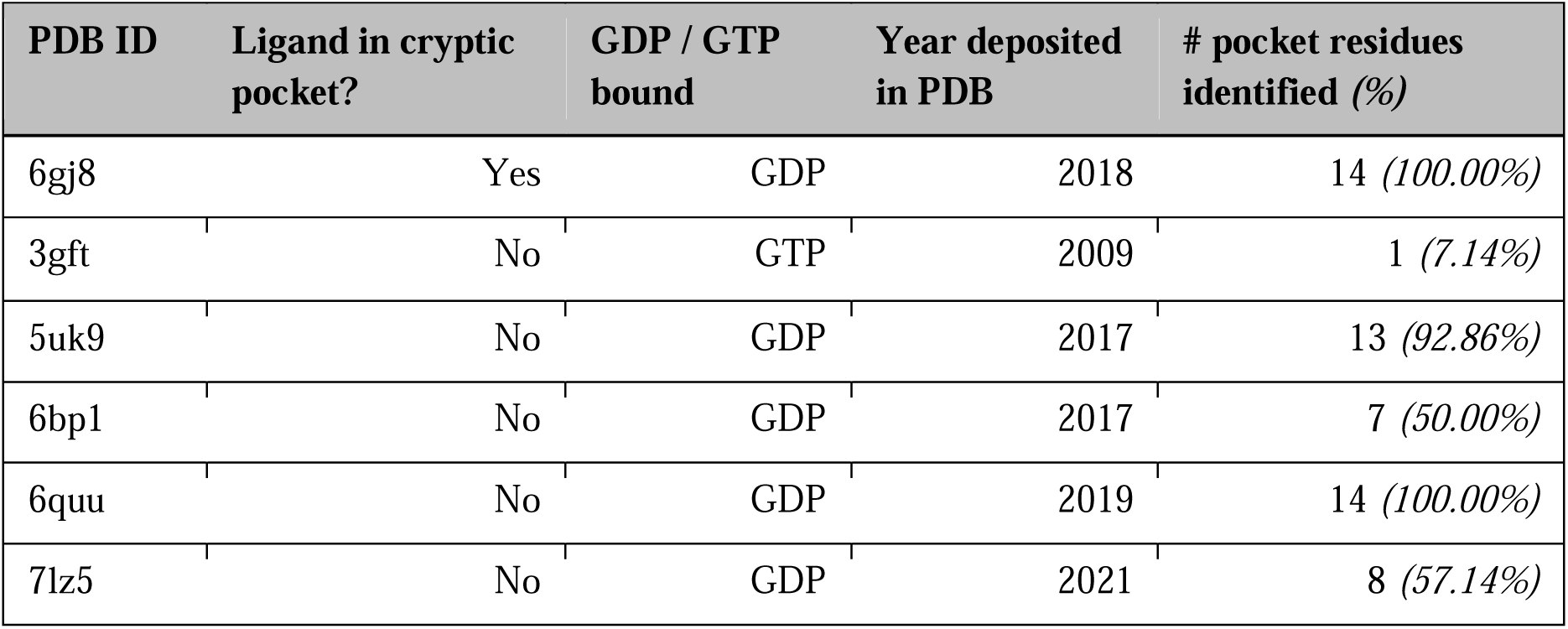
Recovery of switch I/II cryptic pocket on various KRAS structures when predicted by *hotpocketNN*.

**Table S9.**
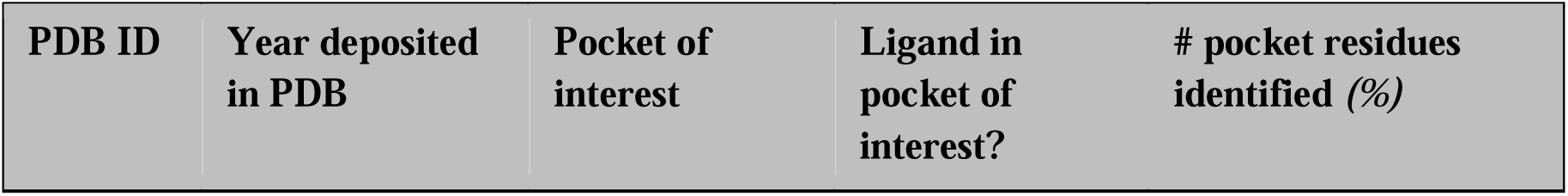
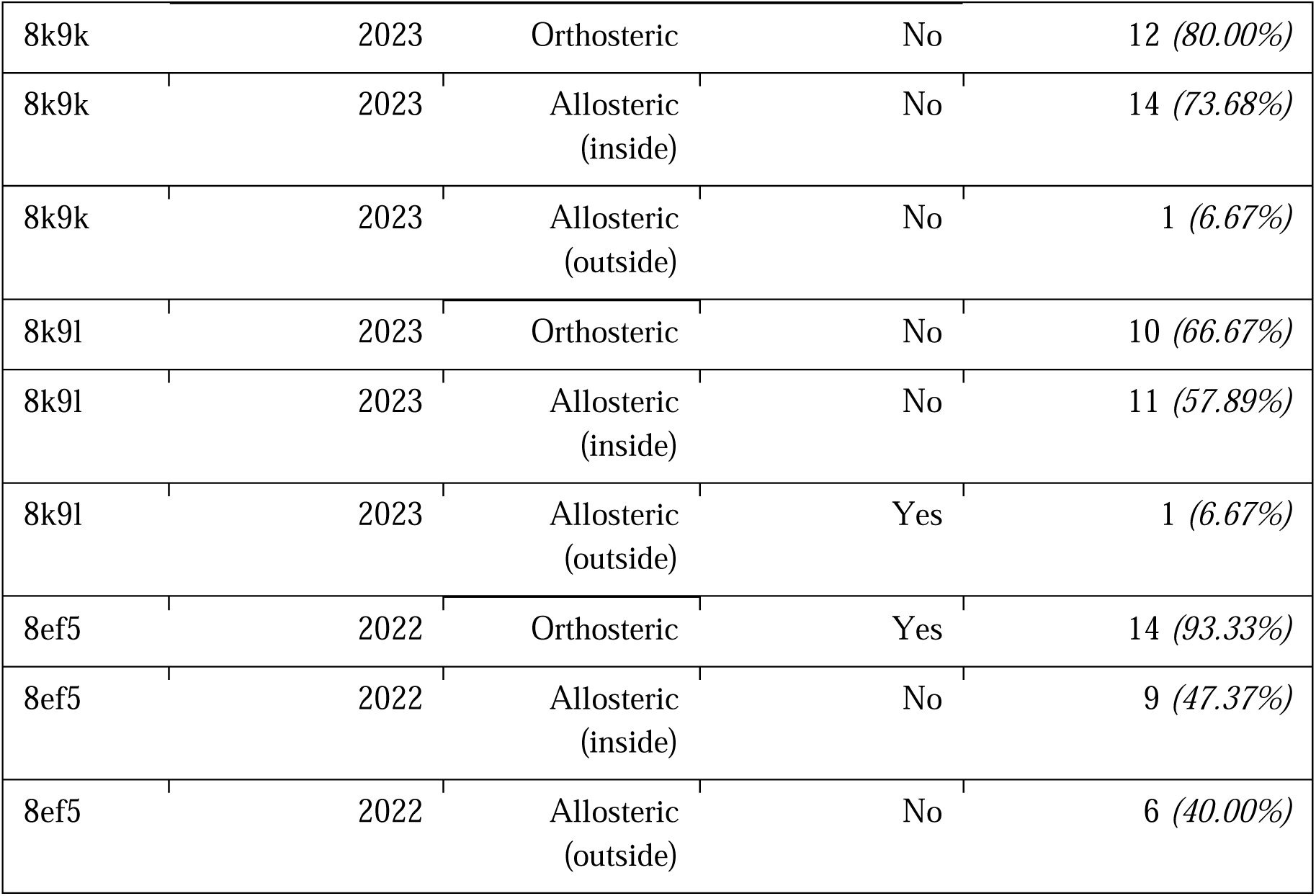
Recovery of orthosteric and allosteric pockets on various mOR structures when predicted by *hotpocketNN*. “Allosteric (inside)” is the allosteric pocket shown in 9bjk; “allosteric (outside)” is the allosteric pocket shown in 8k9l.

**Figure S1.**
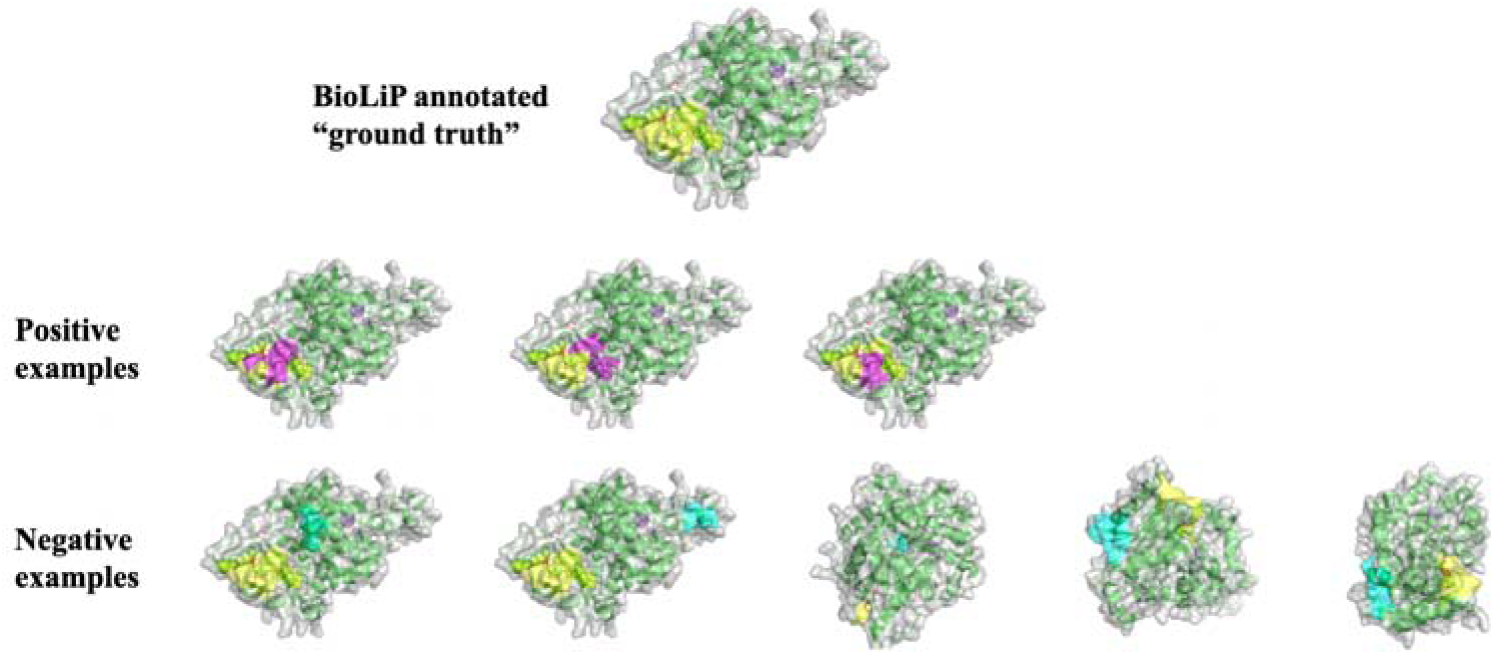
Schematic depicting process for generating positive and negative pocket examples from BioLiP known binding pocket annotations, to train, validate, and test the ML filtering method.

**Figure S2.**
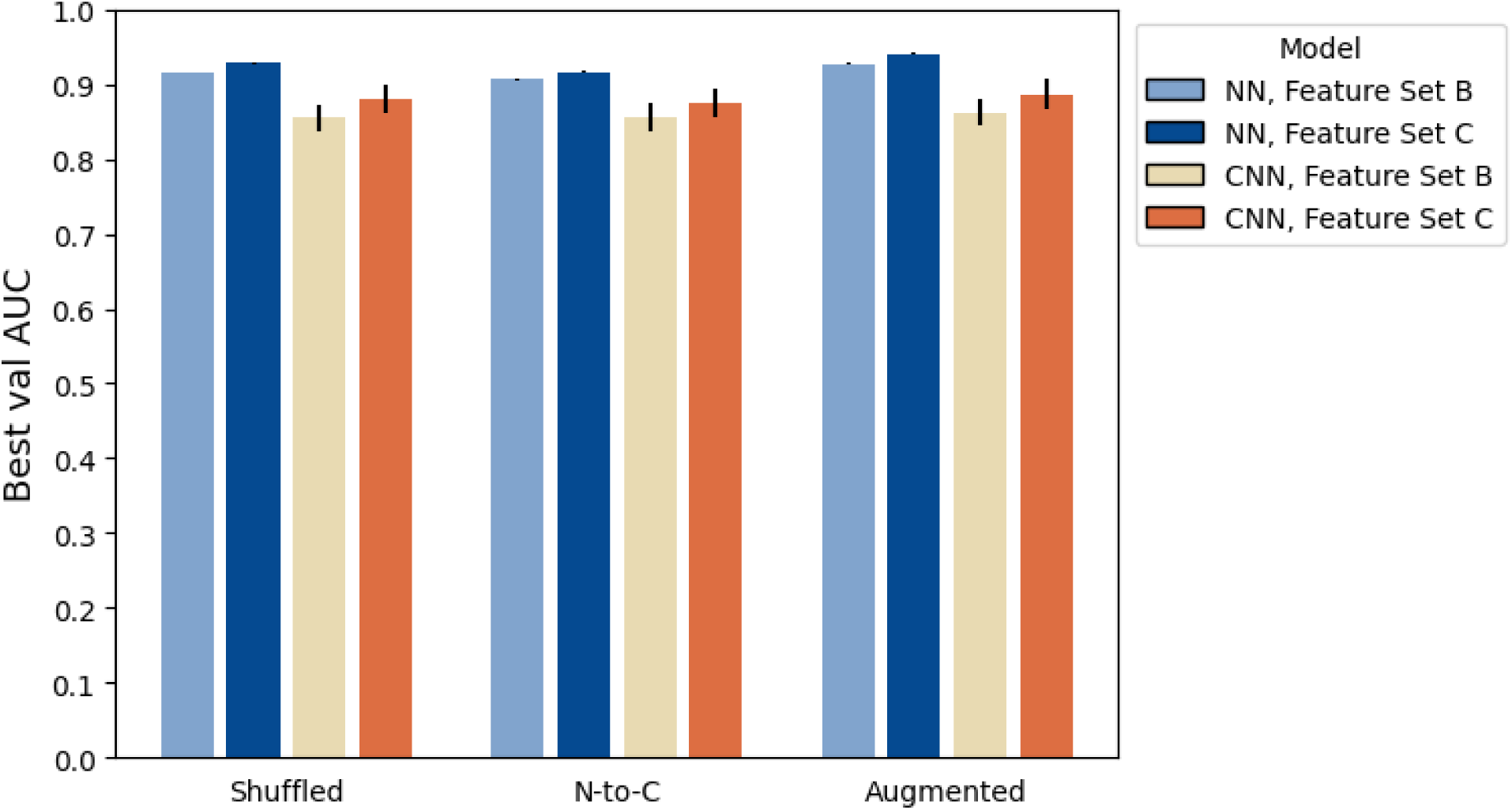
Performance of different pocket residue ordering schemes for both NNs and CNNs, and Feature Sets B and C. Average performance across different hyperparameter settings is shown with standard error.

**Figure S3.**
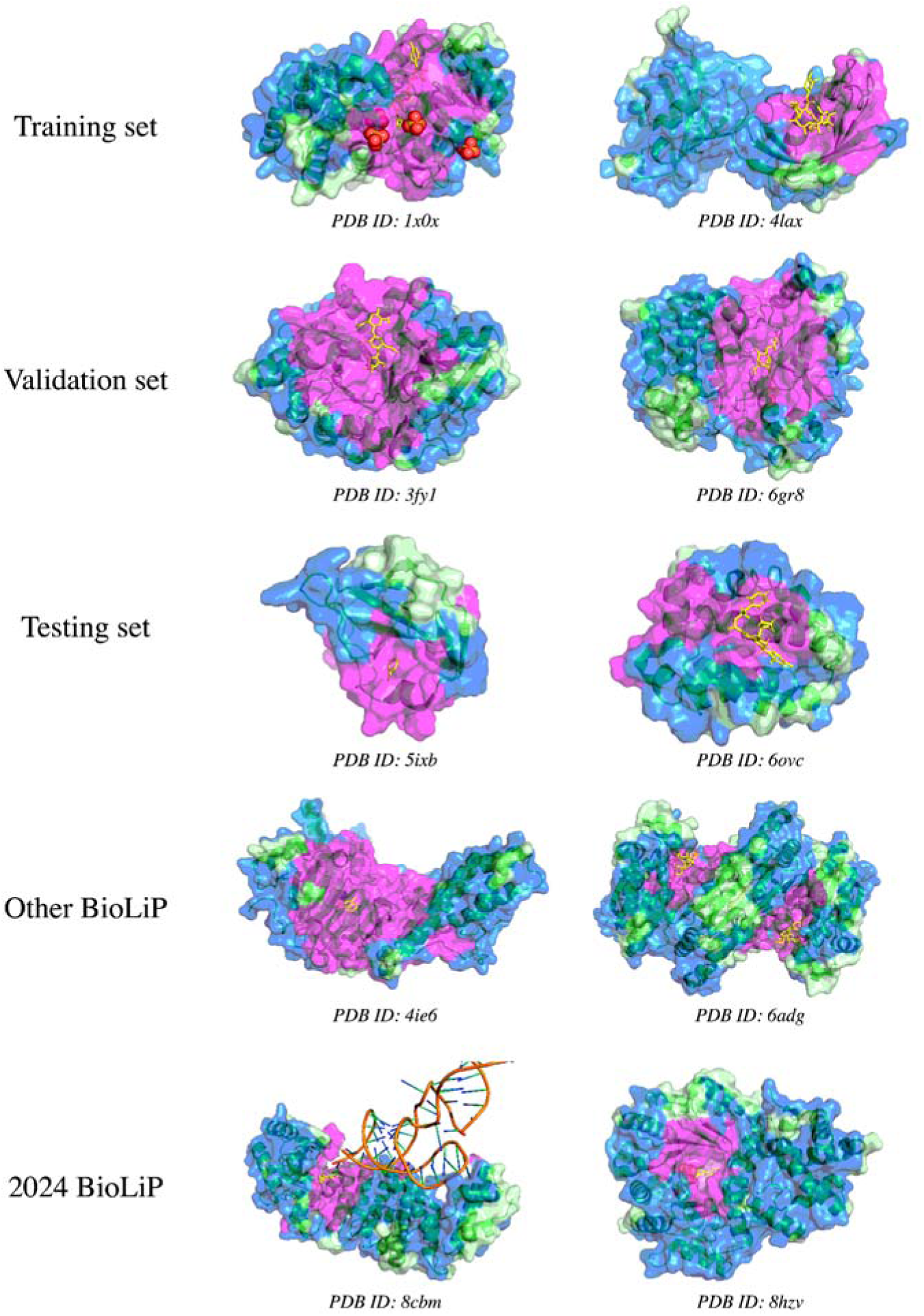
Visualizations of accepted and rejected candidate pockets on experimentally-determined structures from the PDB, using ESM2 embeddings as features for *hotpocketNN* ensembling and filtering method. From top row to bottom row, structures are taken from: the *hotpocketNN* training set, the *hotpocketNN* validation set, the *hotpocketNN* testing set, human protein structures with BioLiP annotations not included in the *hotpocketNN* train/val/test sets, and human protein structures with BioLiP annotations released in 2024 with low sequence identity to previously-seen structures. The surface of the protein structure is colored as follows: magenta for residues that are part of an accepted candidate pocket accepted by *hotpocketNN*, blue for residues that are part of a candidate pocket but not an accepted candidate pocket, and light green for residues that are not part of any candidate pockets. Biologically-relevant ligands are visualized in the structure as yellow sticks; non-biologically-relevant ligands are omitted. For each structure, we show only a single biological assembly; for the “2024 BioLiP” structures, we show only a single chain to better see the novel protein-ligand interaction.

**Figure S4.**
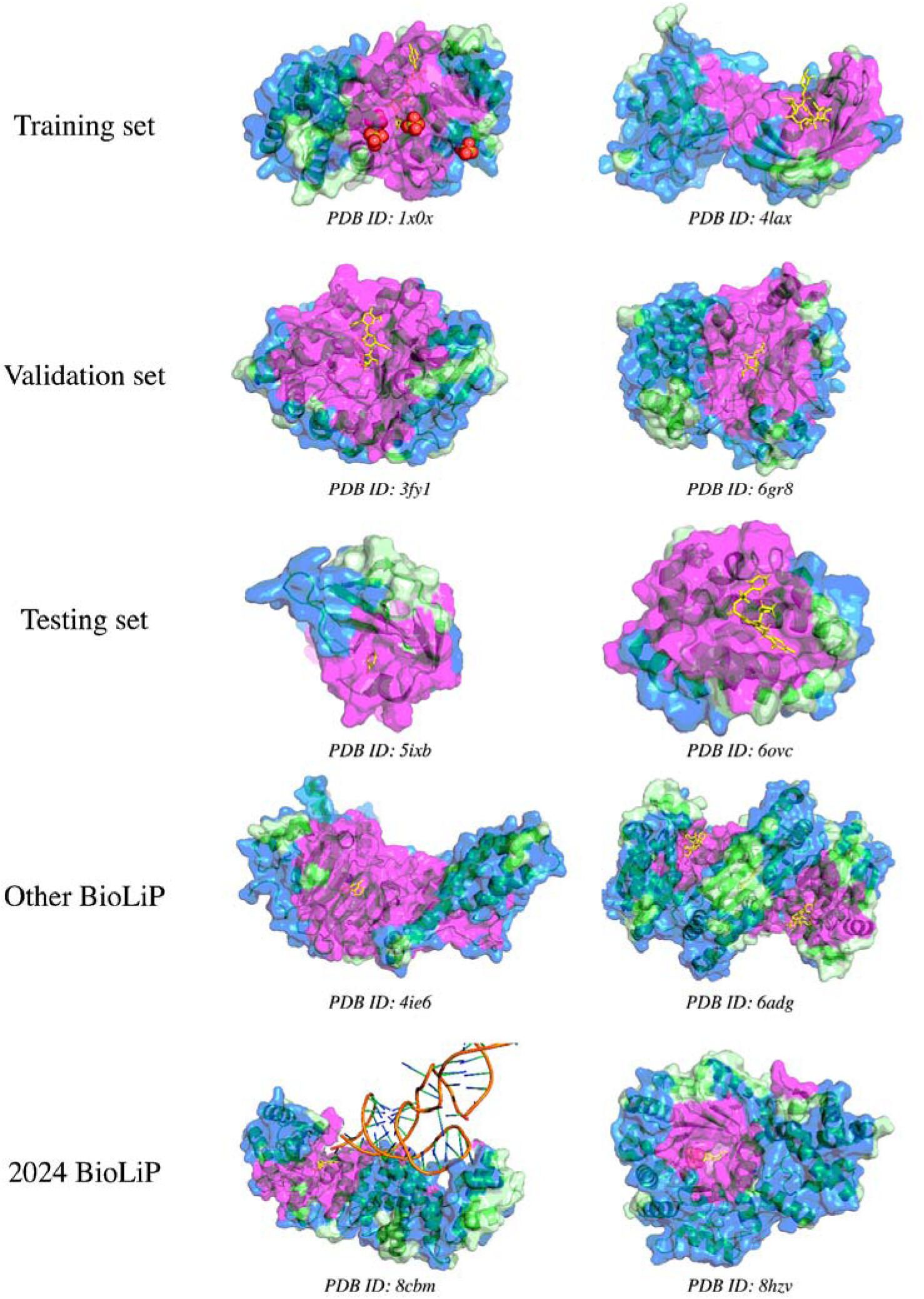
Visualizations of accepted and rejected candidate pockets on experimentally-determined structures from the PDB, using constituent method predictions and ESM2 embeddings as features for *hotpocketNN* ensembling and filtering method. From top row to bottom row, structures are taken from: the *hotpocketNN* training set, the *hotpocketNN* validation set, the *hotpocketNN* testing set, human protein structures with BioLiP annotations not included in the *hotpocketNN* train/val/test sets, and human protein structures with BioLiP annotations released in 2024 with low sequence identity to previously-seen structures. The surface of the protein structure is colored as follows: magenta for residues that are part of an accepted candidate pocket accepted by *hotpocketNN*, blue for residues that are part of a candidate pocket but not an accepted candidate pocket, and light green for residues that are not part of any candidate pockets. Biologically-relevant ligands are visualized in the structure as yellow sticks; non-biologically-relevant ligands are omitted. For each structure, we show only a single biological assembly; for the “2024 BioLiP” structures, we show only a single chain to better see the novel protein-ligand interaction.

**Figure S5.**
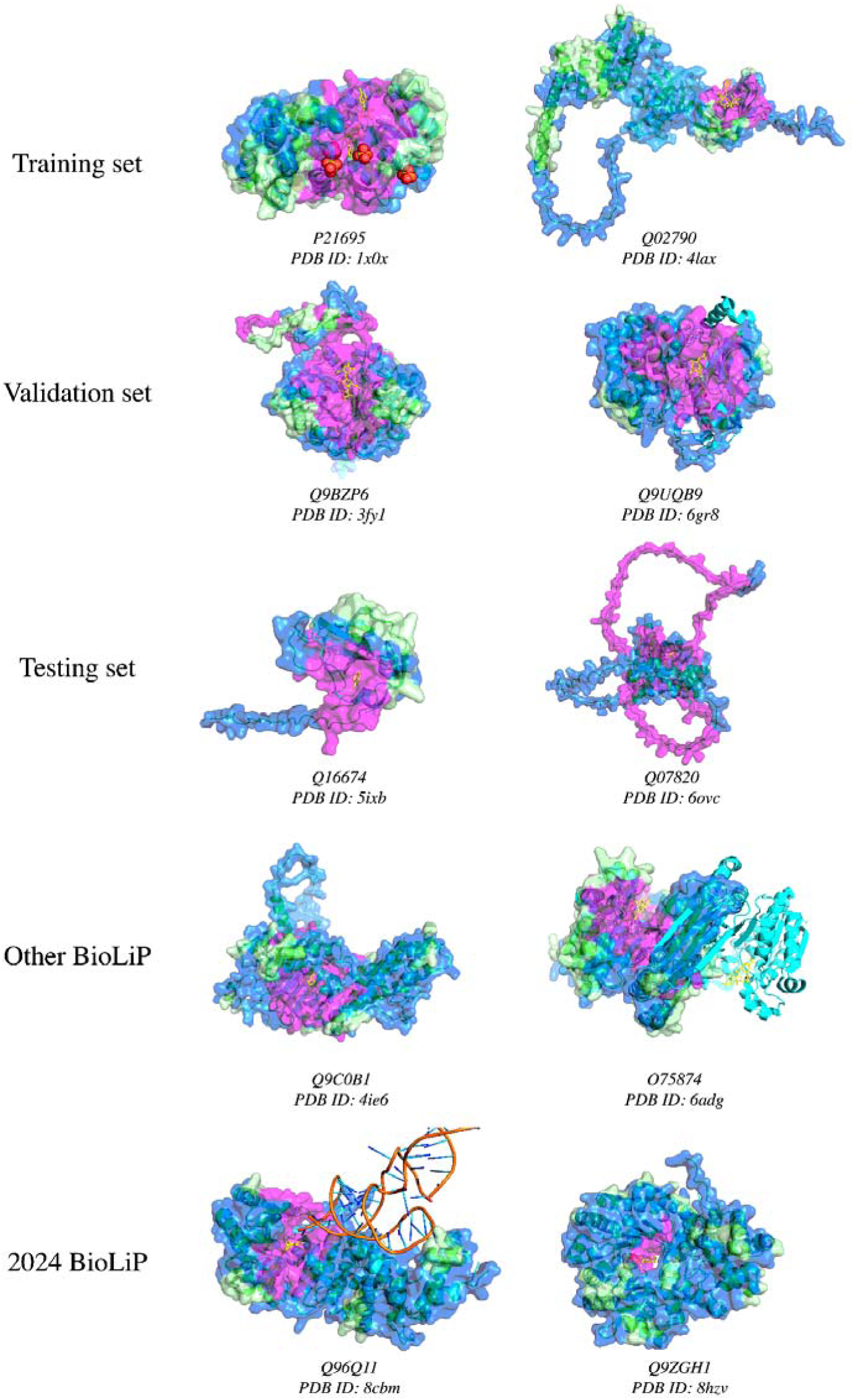
Visualizations of accepted and rejected candidate pockets on AlphaFold2-predicted protein structures, using ESM2 embeddings as features for *hotpocketNN* ensembling and filtering method. AlphaFold2-predicted structures (green ribbons) are shown aligned with their experimentally-determined PDB counterparts (cyan ribbons). Only the surface of the predicted structure is shown and pocket predictions were made on the predicted structure. The aligned ligands from the experimentally-determined structure are shown in yellow; these ligands are not a part of the AlphaFold2-predicted structures. The chains of the experimentally-determined structures are the same as in Figure 5. From top row to bottom row, structures are taken from: the *hotpocketNN* training set, the *hotpocketNN* validation set, the *hotpocketNN* testing set, human protein structures with BioLiP annotations not included in the *hotpocketNN* train/val/test sets, and human protein structures with BioLiP annotations released in 2024 with low sequence identity to previously-seen structures. The surface of the protein structure is colored as follows: magenta for residues that are part of an accepted candidate pocket accepted by *hotpocketNN*, blue for residues that are part of a candidate pocket but not an accepted candidate pocket, and light green for residues that are not part of any candidate pockets.

**Figure S6.**
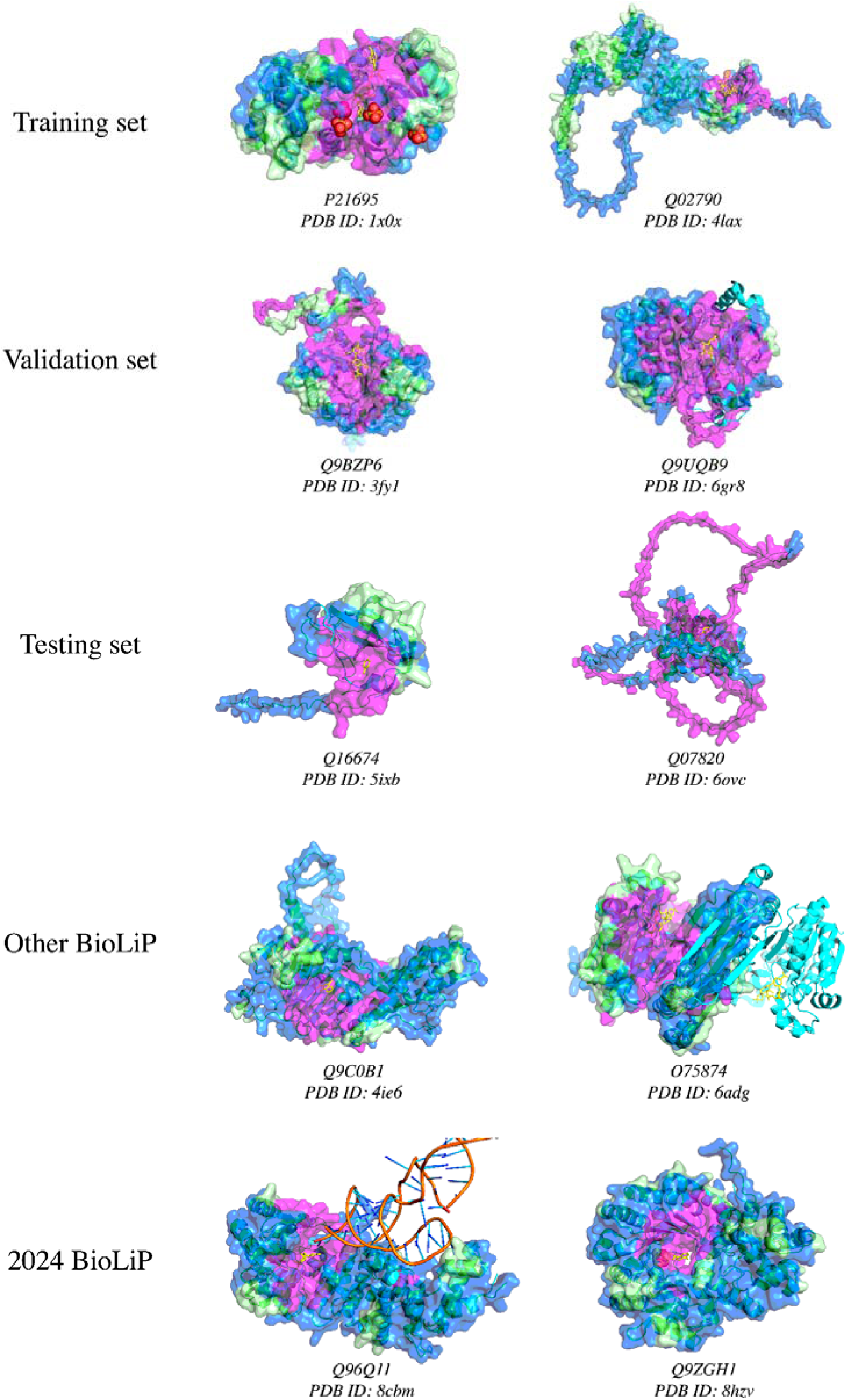
Visualizations of accepted and rejected candidate pockets on AlphaFold2-predicted protein structures, using constituent method predictions and ESM2 embeddings as features for *hotpocketNN* ensembling and filtering method. AlphaFold2-predicted structures (green ribbons) are shown aligned with their experimentally-determined PDB counterparts (cyan ribbons). Only the surface of the predicted structure is shown and pocket predictions were made on the predicted structure. The aligned ligands from the experimentally-determined structure are shown in yellow; these ligands are not a part of the AlphaFold2-predicted structures. The chains of the experimentally-determined structures are the same as in **Figure S3**. From top row to bottom row, structures are taken from: the *hotpocketNN* training set, the *hotpocketNN* validation set, the *hotpocketNN* testing set, human protein structures with BioLiP annotations not included in the *hotpocketNN* train/val/test sets, and human protein structures with BioLiP annotations released in 2024 with low sequence identity to previously-seen structures. The surface of the protein structure is colored as follows: magenta for residues that are part of an accepted candidate pocket accepted by *hotpocketNN*, blue for residues that are part of a candidate pocket but not an accepted candidate pocket, and light green for residues that are not part of any candidate pockets.

**Figure S7.**
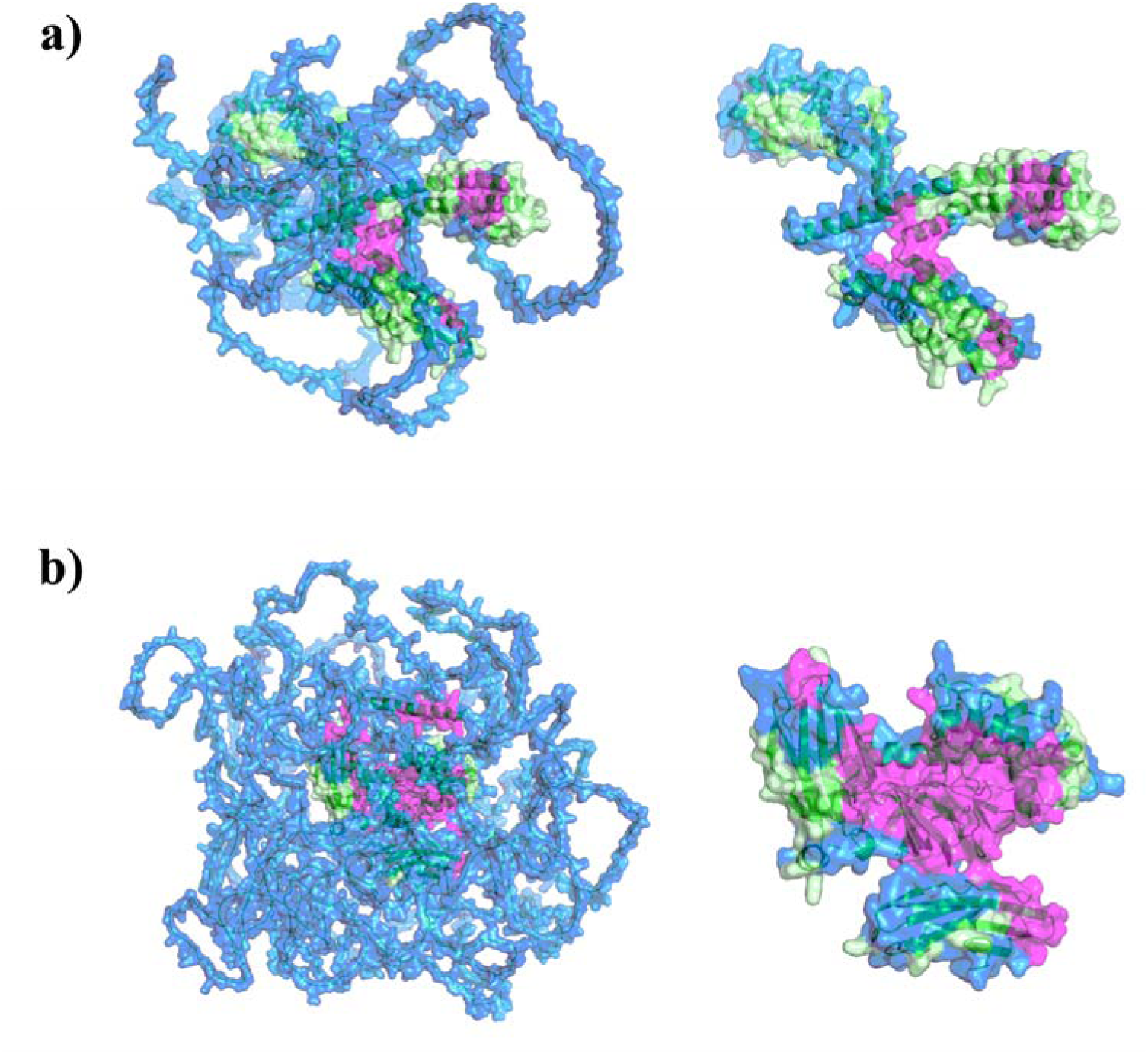
Visualizations of accepted and rejected candidate pockets on AlphaFold2-predicted protein structures with an average per-residue confidence that is **a)** “low” (average pLDDT between 50 and 70; UniProt ID: Q92953), or **b)** “very low” (average pLDDT less than 50; UniProt ID: Q86TB3), using ESM2 embeddings as features for *hotpocketNN* ensembling and filtering method. Neither protein has an experimentally-determined protein structure. The surface of the protein structure is colored as follows: magenta for residues that are part of an accepted candidate pocket accepted by *hotpocketNN*, blue for residues that are part of a candidate pocket but not an accepted candidate pocket, and light green for residues that are not part of any candidate pockets. Both the full predicted protein structure (left) and the subset of the structure that is not low-confidence (pLDDT greater than 70) (right) are shown.

**Figure S8.**
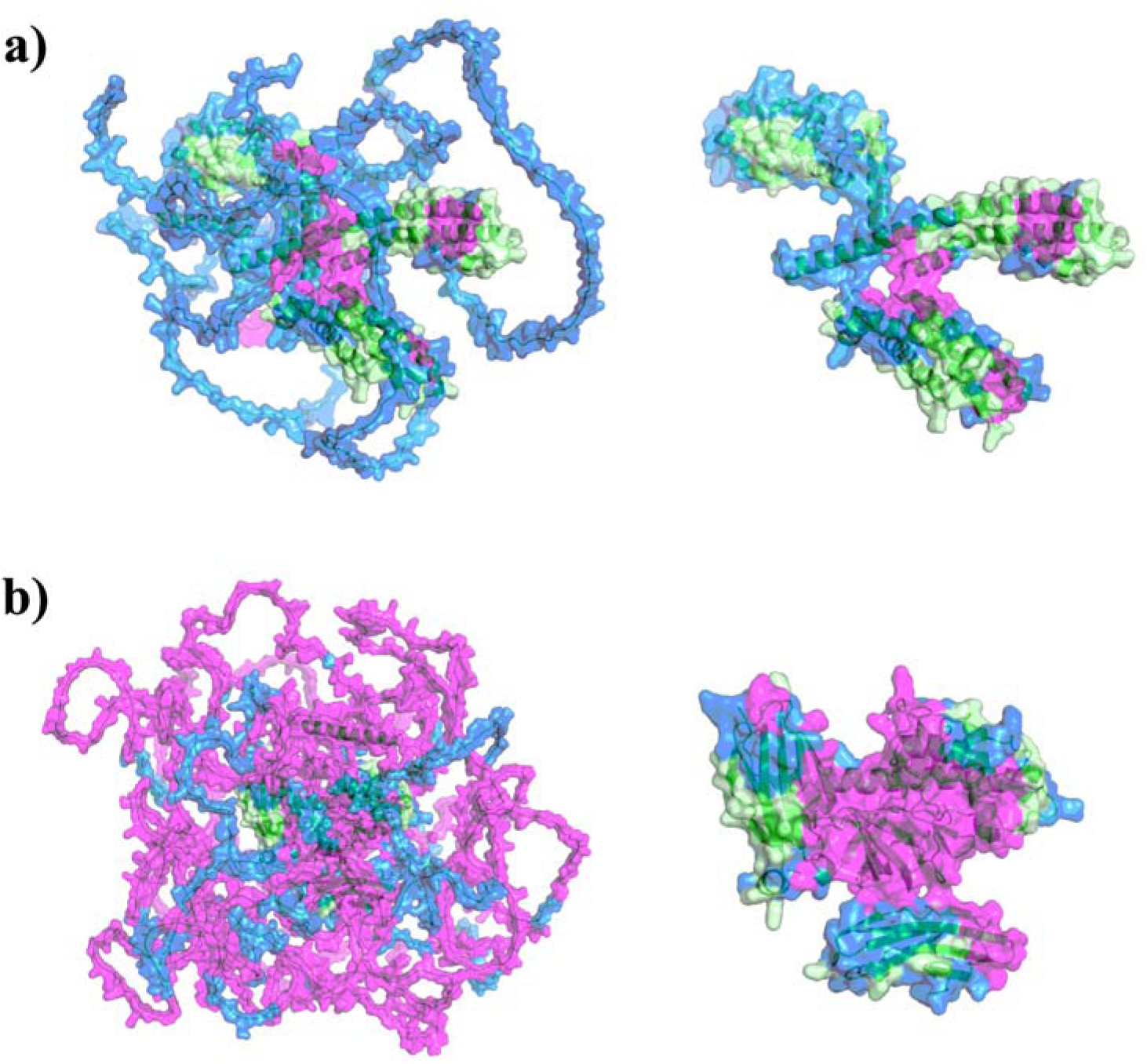
Visualizations of accepted and rejected candidate pockets on AlphaFold2-predicted protein structures with an average per-residue confidence that is **a)** “low” (average pLDDT between 50 and 70; UniProt ID: Q92953), or **b)** “very low” (average pLDDT less than 50; UniProt ID: Q86TB3), using constituent method predictions and ESM2 embeddings as features for *hotpocketNN* ensembling and filtering method. Neither protein has an experimentally-determined protein structure. The surface of the protein structure is colored as follows: magenta for residues that are part of an accepted candidate pocket accepted by *hotpocketNN*, blue for residues that are part of a candidate pocket but not an accepted candidate pocket, and light green for residues that are not part of any candidate pockets. Both the full predicted protein structure (left) and the subset of the structure that is not low-confidence (pLDDT greater than 70) (right) are shown.

**Figure S9.**
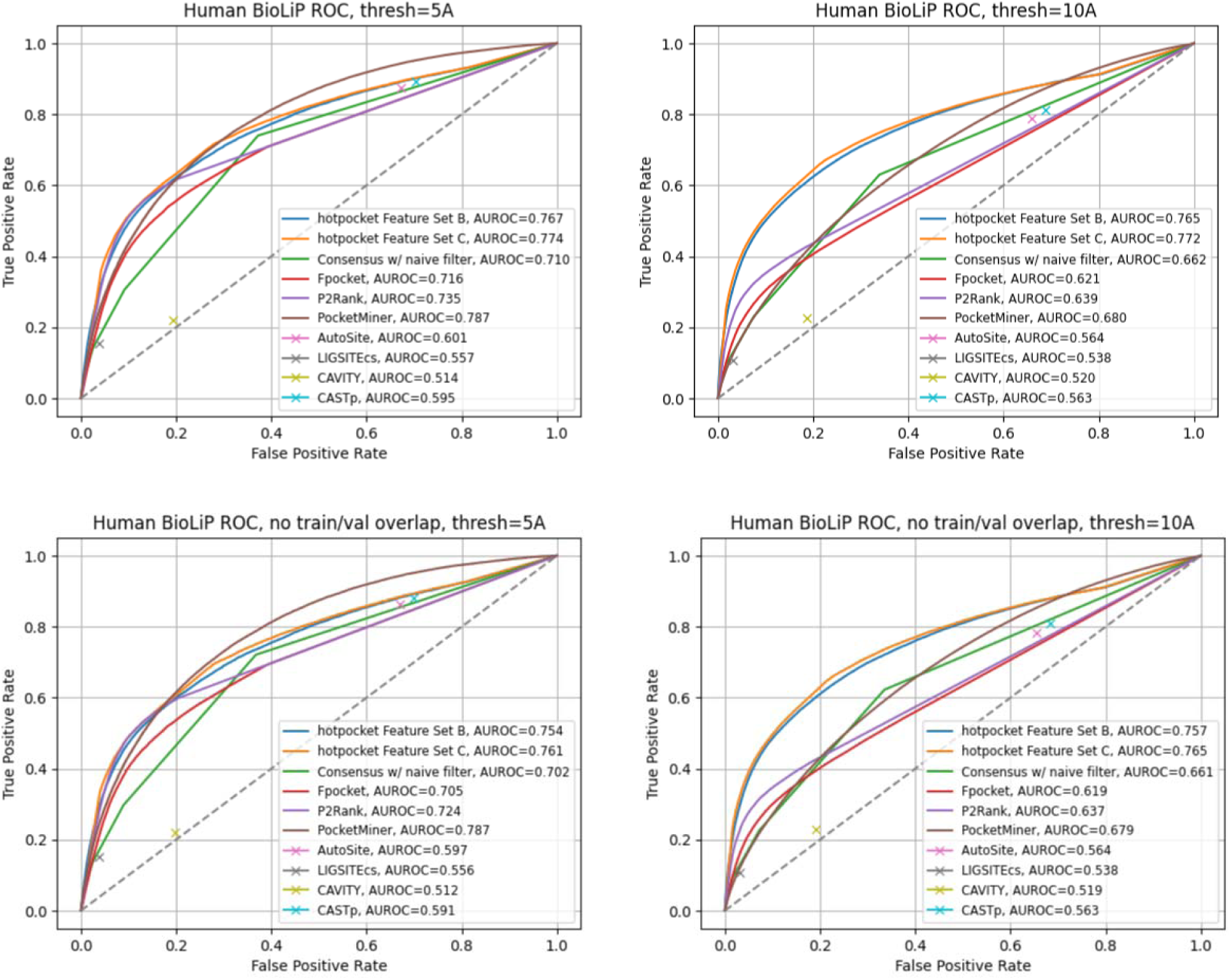
ROC curves and AUROCs for per-residue scoring performance of *hotpocketNN*, naive filter across union of constituent methods, and each constituent pocket-finding method individually across the Human BioLiP dataset. Each residue was labeled as a binding residue if it was within 5 angstroms (left) or 10 angstroms (right) of a relevant ligand. For the *hotpocketNN* models, Fpocket, P2Rank, and PocketMiner, each residue’s predicted score for being a member of a pocket was the maximum of all predicted pockets of which it was a member. AutoSite, LIGSITEcs, CAVITY, and CASTp do not provide pocket scores; for these methods, each residue’s predicted score for being a member of a pocket was 1 if it is part of any predicted pocket for the structure, or 0 otherwise. The top row shows results when all structures in all datasets are included, including structures present in the training and validation sets for the *hotpocketNN*. The bottom row shows results when structures present in the training and validation sets for the *hotpocketNN* are excluded. CAVITY and CASTp did not have predictions available for any PoseBusters structures. Feature Set B is the per-residue ESM2 embeddings; Feature Set C is both the per-residue pocket predictions and per-residue ESM2 embeddings concatenated together.

**Figure S10.**
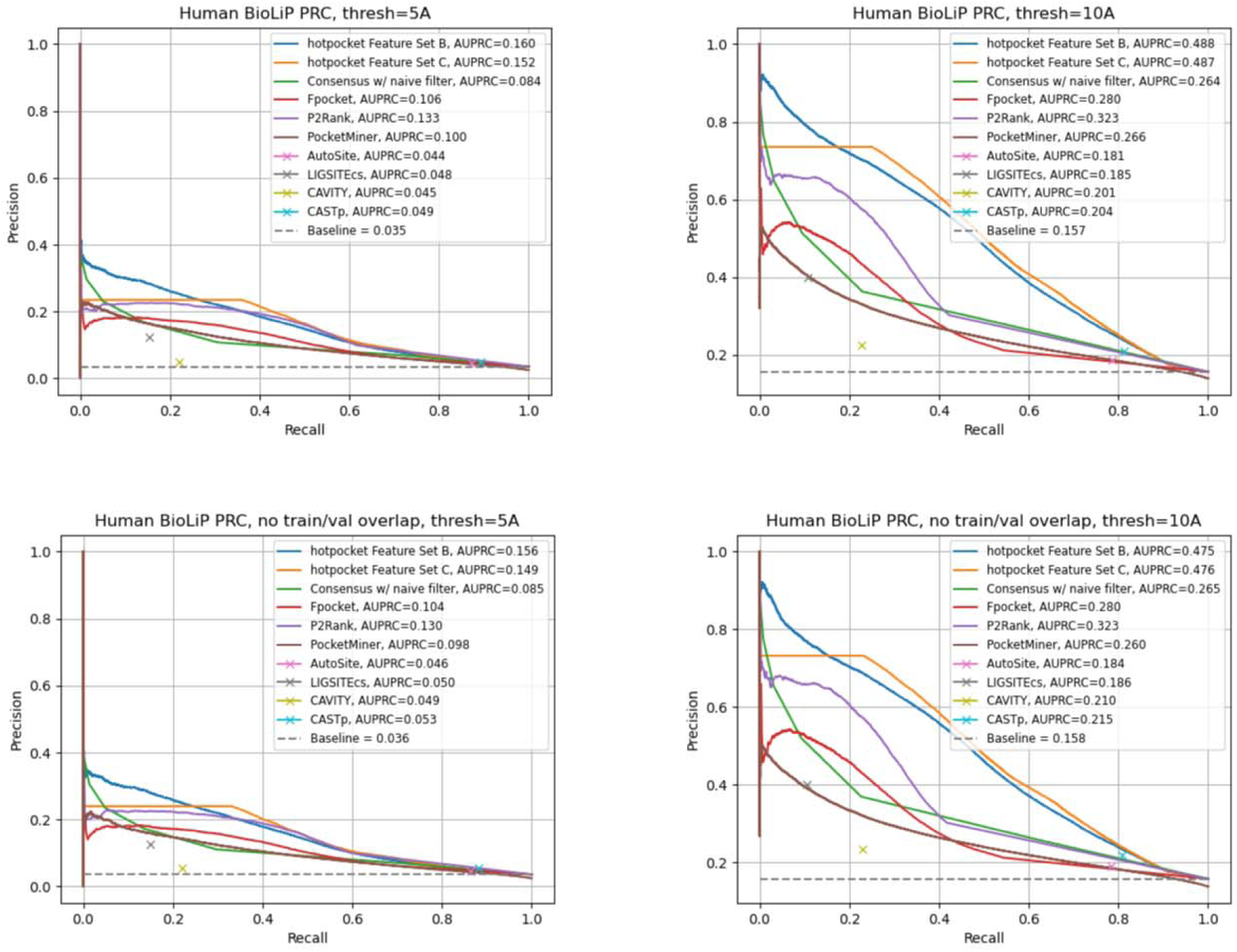
PRC curves and AUPRCs for per-residue scoring performance of *hotpocketNN*, naive filter across union of constituent methods, and each constituent pocket-finding method individually across the Human BioLiP dataset. Each residue was labeled as a binding residue if it was within 5 angstroms (left) or 10 angstroms (right) of a relevant ligand. For the *hotpocketNN* models, Fpocket, P2Rank, and PocketMiner, each residue’s predicted score for being a member of a pocket was the maximum of all predicted pockets of which it was a member. AutoSite, LIGSITEcs, CAVITY, and CASTp do not provide pocket scores; for these methods, each residue’s predicted score for being a member of a pocket was 1 if it is part of any predicted pocket for the structure, or 0 otherwise. The top row shows results when all structures in all datasets are included, including structures present in the training and validation sets for the *hotpocketNN*. The bottom row shows results when structures present in the training and validation sets for the *hotpocketNN* are excluded. CAVITY and CASTp did not have predictions available for any PoseBusters structures. Feature Set B is the per-residue ESM2 embeddings; Feature Set C is both the per-residue pocket predictions and per-residue ESM2 embeddings concatenated together.

## Footnotes

^1^https://ccsb.scripps.edu/autosite/downloads/

^2^http://sts.bioe.uic.edu/castp/index.html

^3^http://www.pkumdl.cn:8000/cavityspace/downloads

^4^https://github.com/Discngine/fpocket

^5^https://github.com/ludlows/BALL

^6^https://github.com/rdk/p2rank

^7^https://github.com/Mickdub/gvp/tree/pocket_pred

^8^https://zhanggroup.org/BioLiP/download.html

