## Supplementary file for "The Human Omnibus of Targetable Pockets"

### **Supplement**

**Text S1.** List of excluded BioLiP ligand identifiers for “druglike ligand” filter to include only organic small molecules.

dna, rna, peptide, ACT, CA, CO, CO3, CU, FE, K, MG, MN, PO4, SO3, SO4, ZN

**Text S2.** Naive pocket filtering criteria.

AutoSite

Built-in score: None provided besides ranked pocket order.

Filter: Include only first five pockets.

CASTp

Built-in score: None provided besides ranked pocket order.

Filter: Include only first five pockets.

CavitySpace

Built-in score: Categorical pocket druggability classes.

Filter: Include only pockets labeled with “Strong” druggability.

Fpocket

Built-in score: Pocket druggability score ranging from 0 to 1.

Filter: Include only pockets with druggability score of at least 0.5.

LIGSITEcs

Built-in score: None provided besides ranked pocket order.

Filter: Include only first five pockets.

P2Rank

Built-in score: Pocket probability scores ranging from 0 to 1.

Filter: Include only pockets with probability score of at least 0.5.

PocketMiner

Built-in score: Per-residue pocket likelihood score ranging from 0 to 1; aggregated score cutoff of 0.7 already accounted for in method for generating predicted pockets.

Filter: Include only pockets with aggregated pocket likelihood score of at least 0.8.

**Table S1.** Non-comprehensive list of pocket-finding methods from literature review conducted in 2023. See attached table.

**Table S2.** Breakdown of proteins with experimentally-determined protein structures (“PDB”) and proteins with computationally-predicted protein structures (“AF2”) for which pocket annotations and predictions were generated.

| **Method** | **Total # proteins** | **# AF2 proteins** | **# PDB proteins** |
| --- | --- | --- | --- |
| BioLIP | 4,437 | 0 | 4,437 |
| AutoSite | 20,171 | 20,127 | 7,087 |
| CASTp | 5,080 | 0 | 5,080 |
| CAVITY | 14,999 | 13,195 | 4,874 |
| Fpocket | 20,359 | 20,348 | 7,540 |
| LIGSITEcs | 7,083 | 0 | 7,083 |
| P2RANK | 16,996 | 16,066 | 7,080 |
| PocketMiner | 20,015 | 19,820 | 6,145 |

**Table S3.** Logistic regression hyperparameter tuning. For each combination of feature set and logistic regression penalty, the C value, L1 ratio, training set AUC, and validation set AUC are shown for the model with the best validation AUC out of all hyperparameter combinations assessed. Models were trained until convergence or a maximum of 500 iterations. There was no variation in performance across different random seeds. Feature set A is the per-residue pocket predictions from each of the constituent methods; Feature set B is the per-residue ESM2 embeddings; Feature set C is both the per-residue pocket predictions and per-residue ESM2 embeddings concatenated together.

| **Feature set** | **Penalty** | **C** | **L1 ratio** | **Train AUC** | **Val AUC** |
| --- | --- | --- | --- | --- | --- |
| A | None | N/A | N/A | 0.7793 | 0.7630 |
| A | L1 | 0.1 | N/A | 0.7779 | 0.7823 |
| A | L2 | 0.01 | N/A | 0.7695 | 0.7650 |
| A | Elastic Net | 0.1 | 0.9 | 0.7802 | 0.7754 |
| B | None | N/A | N/A | 0.9894 | 0.8849 |
| B | L1 | 1000 | N/A | 0.9894 | 0.8380 |
| B | L2 | 1 | N/A | 0.9894 | 0.8468 |
| B | Elastic Net | 1000 | 0.7 | 0.9894 | 0.8399 |
| C | None | N/A | N/A | 0.9894 | 0.8502 |
| C | L1 | 1 | N/A | 0.9894 | 0.8363 |
| C | L2 | 10 | N/A | 0.9894 | 0.8502 |
| C | Elastic Net | 0.1 | 0.3 | 0.9843 | 0.8520 |

**Table S4.** Neural network (NN and CNN) hyperparameter tuning. For each combination of feature set and neural network type (feedforward NN or convolutional NN), the number of convolutional layers, number of feedforward layers, batchsize, dropout probability, learning rate, optimizer, pooling size, training set AUC, and validation set AUC are shown for the model with the best validation AUC out of all hyperparameter combinations assessed. The AUCs shown are averaged over three identical models trained with different random seeds and are from the epoch with the highest validation AUC achieved in xx epochs. Feature set A is the per-residue pocket predictions from each of the constituent methods; Feature set B is the per-residue ESM2 embeddings; Feature set C is both the per-residue pocket predictions and per-residue ESM2 embeddings concatenated together.

| **NN type** | **Feature set** | **# conv layers** | **# dense layers** | **Batchsize** | **Dropout** | **Learning rate** | **Optimizer** | **Pool size** | **Best mean train AUC** *(std.)* | **Best mean val AUC** *(std.)* |
| --- | --- | --- | --- | --- | --- | --- | --- | --- | --- | --- |
| NN | A | N/A | 3 | 32 | 0.5 | 0.001 | Adam | N/A | 0.8582 *(0.0107)* | 0.8745 *(0.0005)* |
| NN | B | N/A | 1 | 32 | 0.5 | 0.001 | AdamW | N/A | 0.9963 *(0.0048)* | 0.9208 *(0.0018)* |
| NN | C | N/A | 1 | 64 | 0.4 | 0.01 | Adam | N/A | 0.9995 *(0.0002)* | 0.9344 *(0.0019)* |
| CNN | A | 2 | 3 | 128 | 0.2 | 0.001 | Adam | Any | 0.8803 *(0.0076)* | 0.8748 (*0.0010)* |
| CNN | B | 1 | 5 | 128 | 0.0 | 0.001 | Adam | 64 | 0.9966 *(0.0022)* | 0.8907 *(0.0145)* |
| CNN | C | 1 | 5 | 128 | 0.0 | 0.001 | Adam | 128 | 0.9964 *(0.0026)* | 0.9072 *(0.0028)* |

**Table S5.** DCCcriterion using a threshold of 8A for top $N_{L}$ and top $N_{L}$+2 pockets generated by each method, where $N_{L}$ is the number of biologically-relevant ligands in each structure. DCCcriterion is defined as the percentage of pockets that have a center of mass within the threshold distance (here, 8A) of a biologically-relevant ligand; its values range from 0 to 100, with higher values indicating better performance. Any structures present in the *hotpocketNN* training or validation sets were omitted (401 structures removed from Human BioLiP dataset, 1 structure removed from Astex Diverse Set, 0 structures removed from PoseBusters dataset).

|  | **Astex Diverse Set** | | **PoseBusters** | | **Human BioLiP** | |
| --- | --- | --- | --- | --- | --- | --- |
|  | Top N | Top N+2 | Top N | Top N+2 | Top N | Top N+2 |
| *hotpocketNN*, ESM embs | **66.4** | **63.0** | **73.9** | **70.6** | 52.2 | 45.5 |
| *hotpocketNN*, combined | 55.7 | 53.3 | 58.4 | 54.4 | 42.5 | 37.7 |
| *hotpocketNN*, ESM embs, P2Rank only | 53.6 | 43.4 | 60.4 | 50.8 | 57.4 | **47.5** |
| *hotpocketNN*, combined, P2Rank only | 52.6 | 42.5 | 57.6 | 49.2 | 55.3 | 46.1 |
| AutoSite | 44.1 | 36.6 | 45.0 | 36.3 | 42.4 | 35.0 |
| CASTp | 36.7 | 30.5 | - | - | 36.1 | 29.3 |
| CAVITY | 42.9 | 33.3 | - | - | 27.4 | 22.6 |
| Fpocket | 50.5 | 39.1 | 49.4 | 39.6 | 44.3 | 35.6 |
| LIGSITEcs | 59.2 | 48.0 | 51.8 | 42.3 | 39.7 | 32.9 |
| P2Rank | 55.8 | 43.4 | 57.1 | 46.7 | **57.5** | 45.7 |
| PocketMiner | 58.8 | 56.4 | 56.4 | 56.0 | 44.5 | 43.1 |

**Table S6.** DCCcriterion using a threshold of 10A for top $N_{L}$ and top $N_{L}$+2 pockets generated by each method, where $N_{L}$ is the number of biologically-relevant ligands in each structure. DCCcriterion is defined as the percentage of pockets that have a center of mass within the threshold distance (here, 10A) of a biologically-relevant ligand; its values range from 0 to 100, with higher values indicating better performance. Any structures present in the *hotpocketNN* training or validation sets were omitted (401 structures removed from Human BioLiP dataset, 1 structure removed from Astex Diverse Set, 0 structures removed from PoseBusters dataset).

|  | **Astex Diverse Set** | | **PoseBusters** | | **Human BioLiP** | |
| --- | --- | --- | --- | --- | --- | --- |
|  | Top N | Top N+2 | Top N | Top N+2 | Top N | Top N+2 |
| *hotpocketNN*, ESM embs | **75.0** | **71.9** | **78.4** | **75.5** | 58.4 | 51.5 |
| *hotpocketNN*, combined | 60.9 | 58.3 | 64.2 | 60.2 | 49.0 | 43.8 |
| *hotpocketNN*, ESM embs, P2Rank only | 62.0 | 53.4 | 70.8 | 61.0 | **64.0** | **55.0** |
| *hotpocketNN*, combined, P2Rank only | 61.7 | 51.6 | 67.8 | 59.4 | 61.9 | 53.4 |
| AutoSite | 50.7 | 44.4 | 53.9 | 46.0 | 47.6 | 40.9 |
| CASTp | 42.4 | 36.5 | - | - | 41.8 | 35.4 |
| CAVITY | 57.1 | 50.0 | - | - | 34.6 | 29.7 |
| Fpocket | 55.6 | 45.6 | 57.6 | 47.3 | 49.8 | 41.7 |
| LIGSITEcs | 63.1 | 53.6 | 59.2 | 49.5 | 44.9 | 38.6 |
| P2Rank | 59.1 | 48.9 | 63.6 | 54.6 | 61.5 | 50.4 |
| PocketMiner | 72.8 | 71.2 | 69.2 | 69.5 | 54.3 | 53.1 |

**Table S7.** Summary of proteome-wide pockets after passing through the *hotpocketNN* filter (using Feature Set C: ESM2 embeddings and per-residue constituent method predictions) and the low-confidence AlphaFold2 filter.

| **Method** | **% candidate pockets accepted** | **# pockets** | **# proteins** | **Mean pockets per structure** *(std.)* | **Median pockets per structure** | **Mean pocket size** *(std.)* |
| --- | --- | --- | --- | --- | --- | --- |
| AutoSite | 12.52% | 288,293 | 16,953 | 5.71 *(7.43)* | 3 | 18.59 *(13.96)* |
| CASTp | 12.04% | 268,053 | 4,319 | 12.86 *(16.38)* | 8 | 9.28 *(17.81)* |
| CAVITY | 6.18% | 5,499 | 3,732 | 1.36 *(0.78)* | 1 | 45.89 *(36.29)* |
| Fpocket | 18.55% | 537,892 | 18,778 | 8.93 *(15.13)* | 5 | 11.56 *(7.83)* |
| LIGSITEcs | 35.26% | 58,859 | 5,443 | 1.76 *(0.92)* | 2 | 15.66 *(4.72)* |
| P2Rank | 49.22% | 369,656 | 16,299 | 6.70 *(13.53)* | 3 | 12.59 *(7.95)* |
| PocketMiner | 27.10% | 1,947,380 | 18,088 | 40.98 *(71.15)* | 20 | 11.00 *(0.00)* |

**Table S8.** Recovery of switch I/II cryptic pocket on various KRAS structures when predicted by *hotpocketNN*.

| **PDB ID** | **Ligand in cryptic pocket?** | **GDP / GTP bound** | **Year deposited in PDB** | **# pocket residues identified *(%)*** |
| --- | --- | --- | --- | --- |
| 6gj8 | Yes | GDP | 2018 | 14 *(100.00%)* |
| 3gft | No | GTP | 2009 | 1 *(7.14%)* |
| 5uk9 | No | GDP | 2017 | 13 *(92.86%)* |
| 6bp1 | No | GDP | 2017 | 7 *(50.00%)* |
| 6quu | No | GDP | 2019 | 14 *(100.00%)* |
| 7lz5 | No | GDP | 2021 | 8 *(57.14%)* |

**Table S9.** Recovery of orthosteric and allosteric pockets on various mOR structures when predicted by *hotpocketNN*. “Allosteric (inside)” is the allosteric pocket shown in 9bjk; “allosteric (outside)” is the allosteric pocket shown in 8k9l.

| **PDB ID** | **Year deposited in PDB** | **Pocket of interest** | **Ligand in pocket of interest?** | **# pocket residues identified *(%)*** |
| --- | --- | --- | --- | --- |
| 8k9k | 2023 | Orthosteric | No | 12 *(80.00%)* |
| 8k9k | 2023 | Allosteric (inside) | No | 14 *(73.68%)* |
| 8k9k | 2023 | Allosteric (outside) | No | 1 *(6.67%)* |
| 8k9l | 2023 | Orthosteric | No | 10 *(66.67%)* |
| 8k9l | 2023 | Allosteric (inside) | No | 11 *(57.89%)* |
| 8k9l | 2023 | Allosteric (outside) | Yes | 1 *(6.67%)* |
| 8ef5 | 2022 | Orthosteric | Yes | 14 *(93.33%)* |
| 8ef5 | 2022 | Allosteric (inside) | No | 9 *(47.37%)* |
| 8ef5 | 2022 | Allosteric (outside) | No | 6 *(40.00%)* |

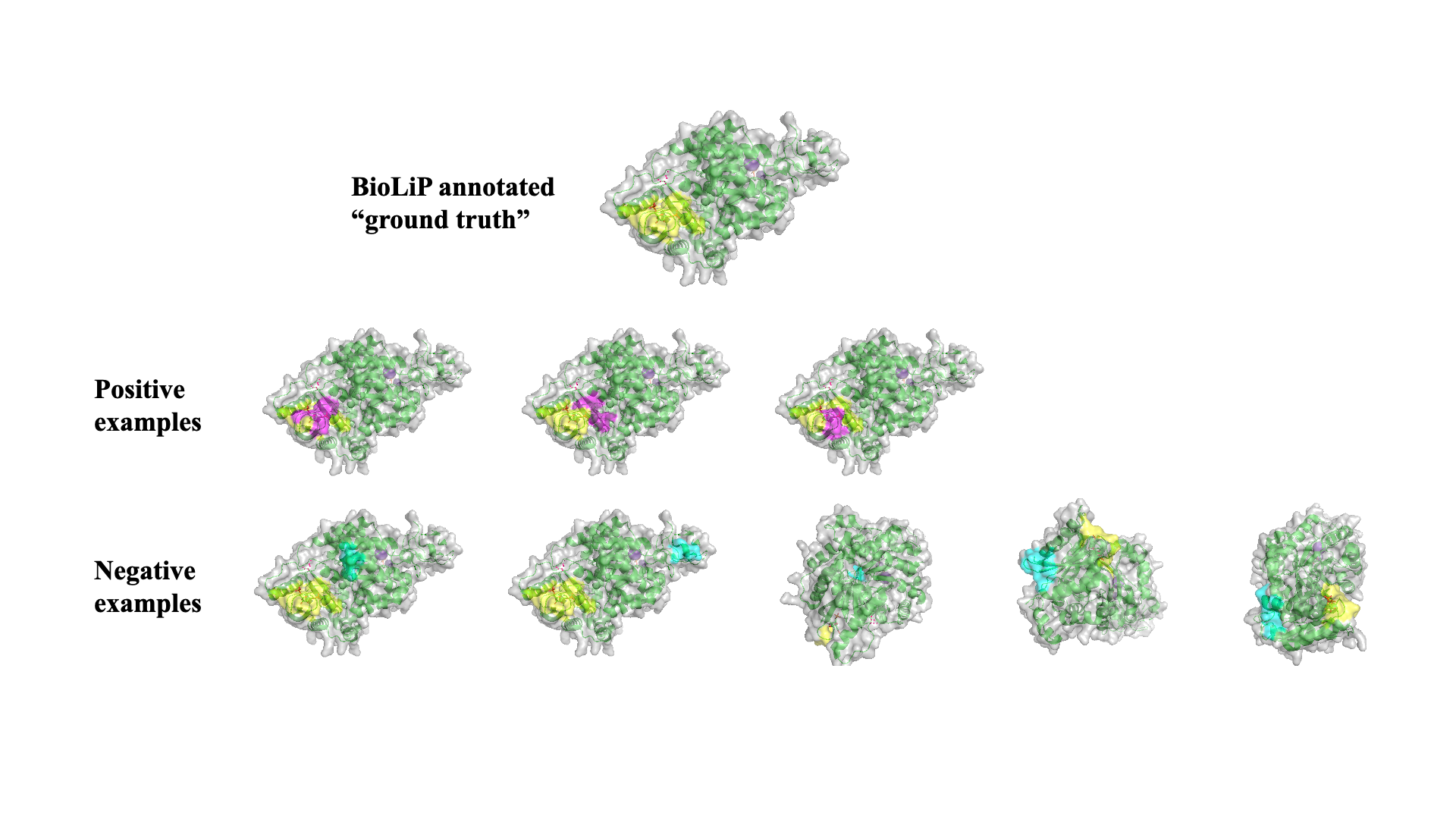

**Figure S1.** Schematic depicting process for generating positive and negative pocket examples from BioLiP known binding pocket annotations, to train, validate, and test the ML filtering method.

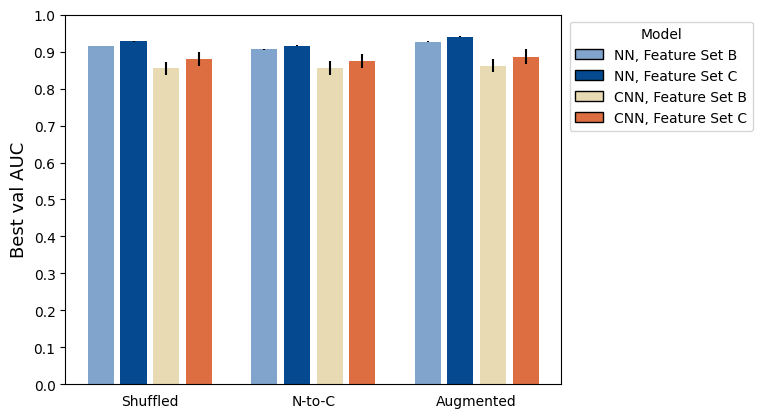

**Figure S2.** Performance of different pocket residue ordering schemes for both NNs and CNNs, and Feature Sets B and C. Average performance across different hyperparameter settings is shown with standard error.

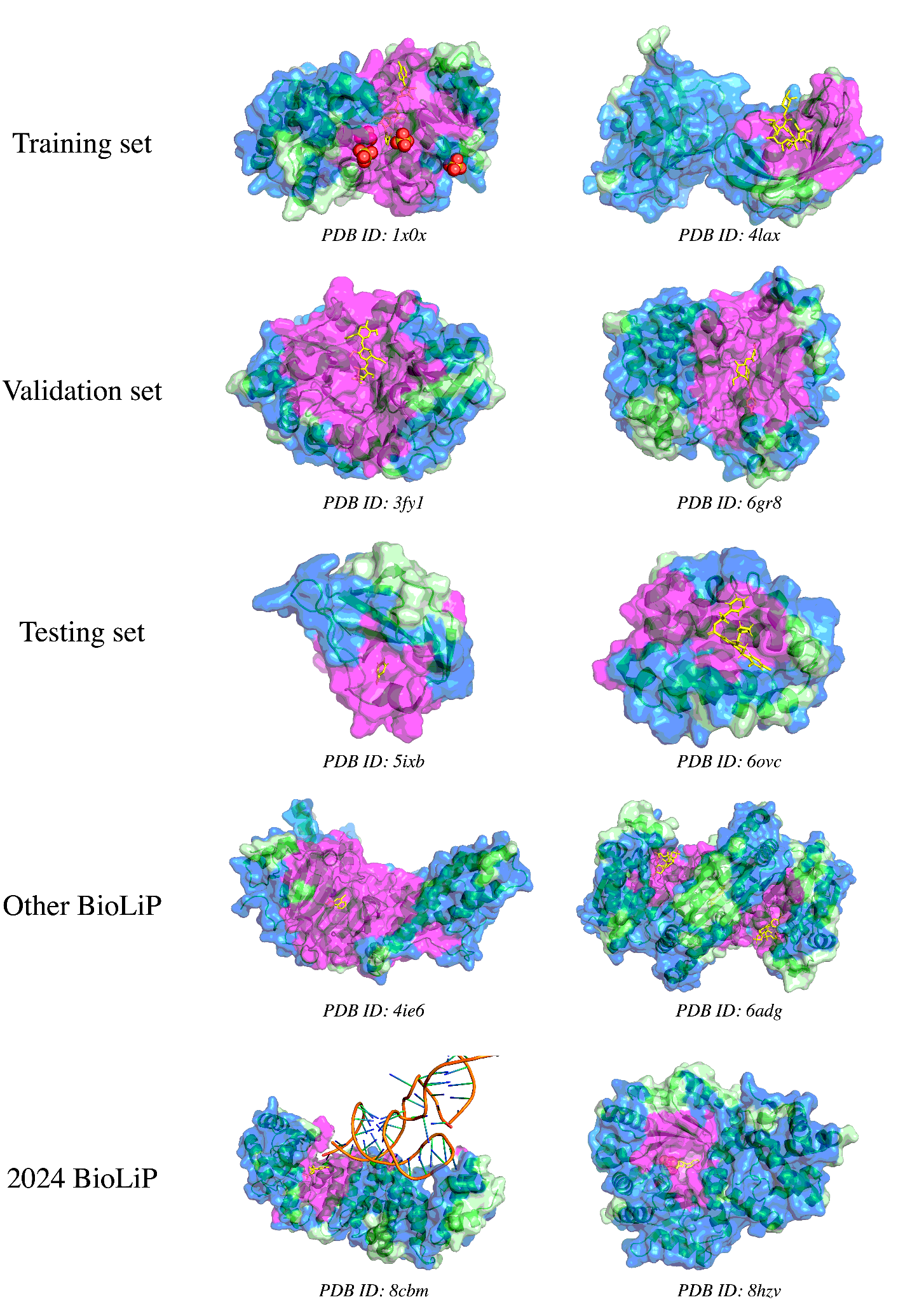

**Figure S3.** Visualizations of accepted and rejected candidate pockets on experimentally-determined structures from the PDB, using ESM2 embeddings as features for *hotpocketNN* ensembling and filtering method. From top row to bottom row, structures are taken from: the *hotpocketNN* training set, the *hotpocketNN* validation set, the *hotpocketNN* testing set, human protein structures with BioLiP annotations not included in the *hotpocketNN* train/val/test sets, and human protein structures with BioLiP annotations released in 2024 with low sequence identity to previously-seen structures. The surface of the protein structure is colored as follows: magenta for residues that are part of an accepted candidate pocket accepted by *hotpocketNN*, blue for residues that are part of a candidate pocket but not an accepted candidate pocket, and light green for residues that are not part of any candidate pockets. Biologically-relevant ligands are visualized in the structure as yellow sticks; non-biologically-relevant ligands are omitted. For each structure, we show only a single biological assembly; for the “2024 BioLiP” structures, we show only a single chain to better see the novel protein-ligand interaction.

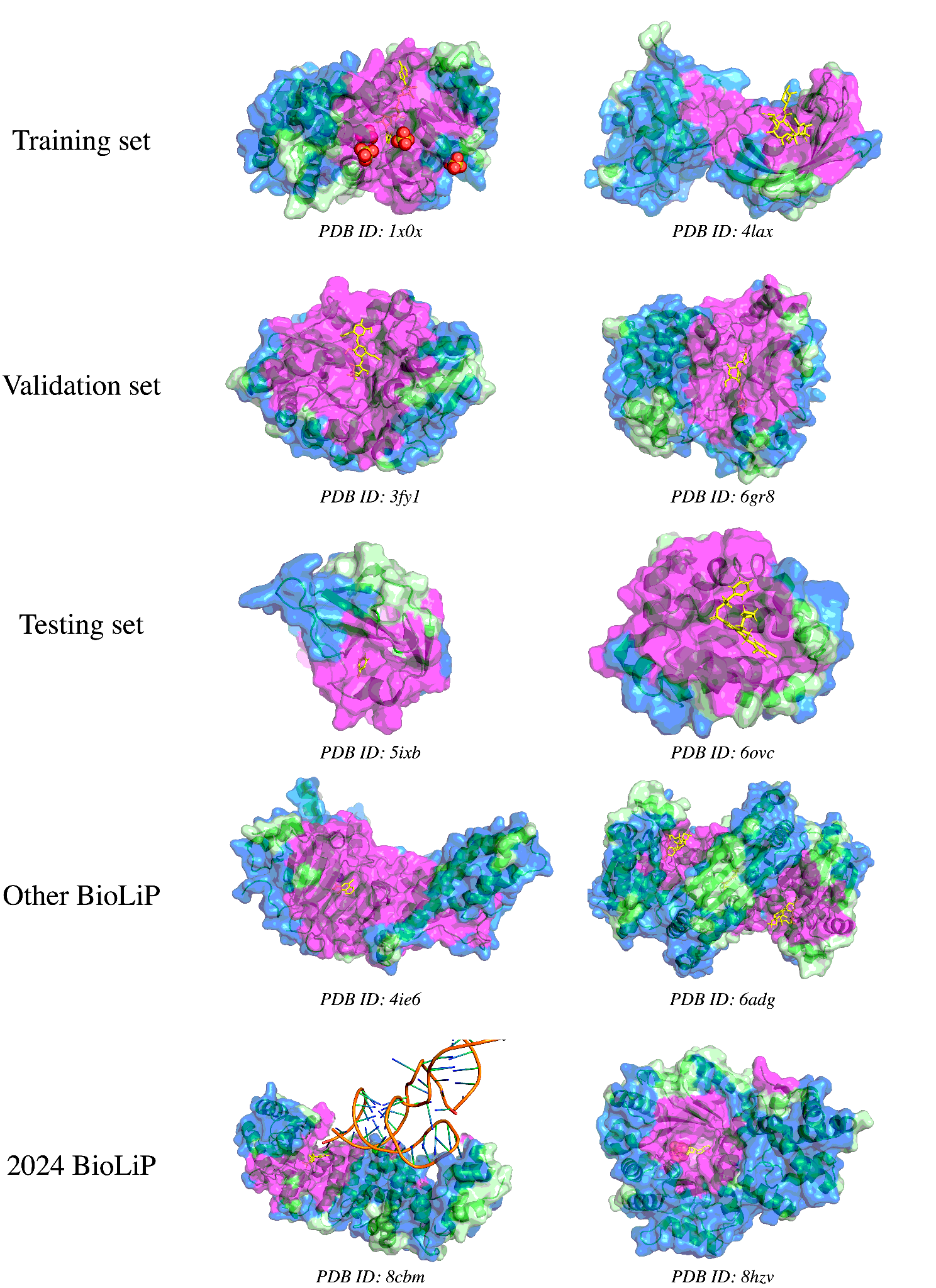

**Figure S4.** Visualizations of accepted and rejected candidate pockets on experimentally-determined structures from the PDB, using constituent method predictions and ESM2 embeddings as features for *hotpocketNN* ensembling and filtering method. From top row to bottom row, structures are taken from: the *hotpocketNN* training set, the *hotpocketNN* validation set, the *hotpocketNN* testing set, human protein structures with BioLiP annotations not included in the *hotpocketNN* train/val/test sets, and human protein structures with BioLiP annotations released in 2024 with low sequence identity to previously-seen structures. The surface of the protein structure is colored as follows: magenta for residues that are part of an accepted candidate pocket accepted by *hotpocketNN*, blue for residues that are part of a candidate pocket but not an accepted candidate pocket, and light green for residues that are not part of any candidate pockets. Biologically-relevant ligands are visualized in the structure as yellow sticks; non-biologically-relevant ligands are omitted. For each structure, we show only a single biological assembly; for the “2024 BioLiP” structures, we show only a single chain to better see the novel protein-ligand interaction.

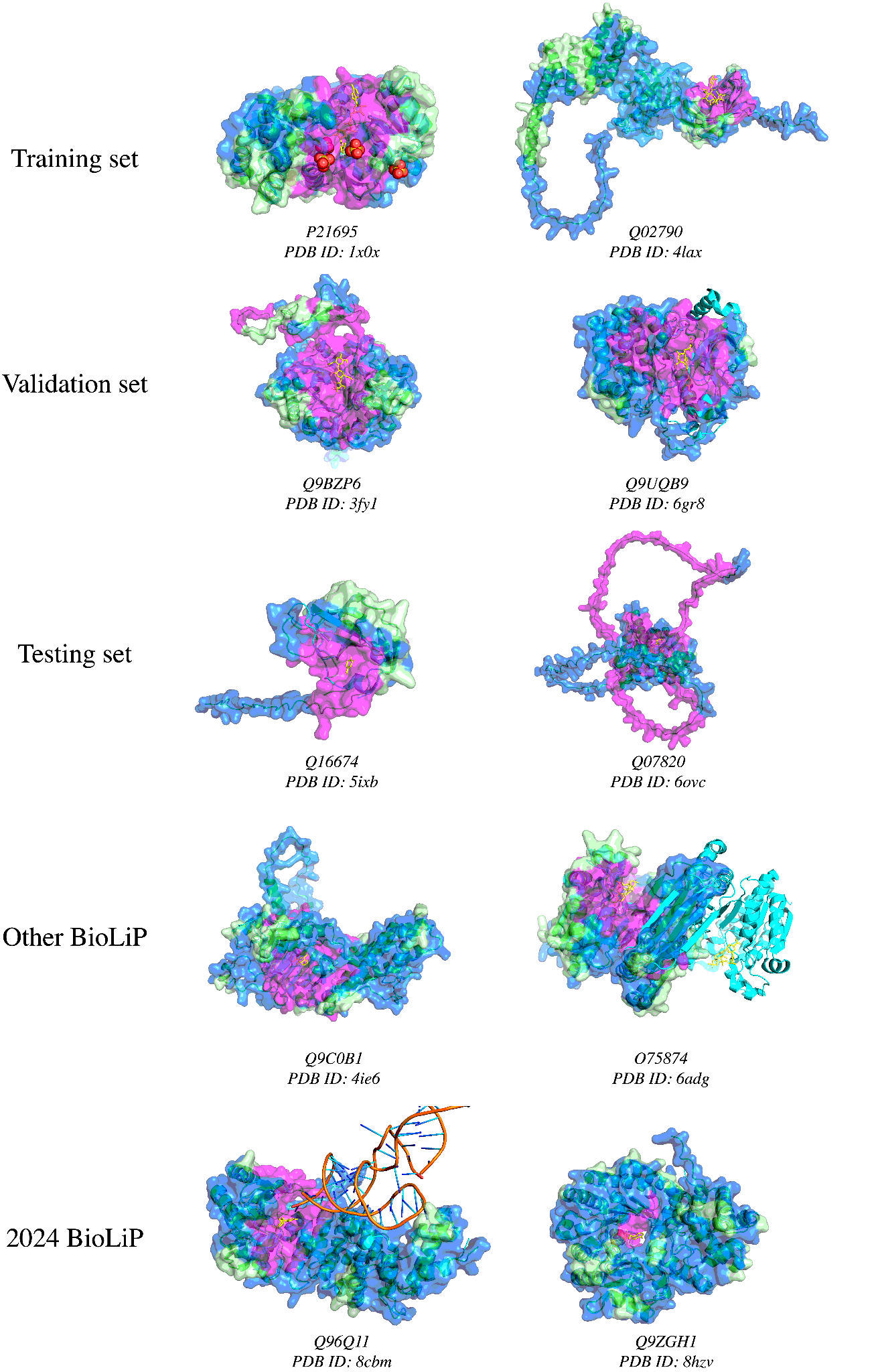

**Figure S5.** Visualizations of accepted and rejected candidate pockets on AlphaFold2-predicted protein structures, using ESM2 embeddings as features for *hotpocketNN* ensembling and filtering method. AlphaFold2-predicted structures (green ribbons) are shown aligned with their experimentally-determined PDB counterparts (cyan ribbons). Only the surface of the predicted structure is shown and pocket predictions were made on the predicted structure. The aligned ligands from the experimentally-determined structure are shown in yellow; these ligands are not a part of the AlphaFold2-predicted structures. The chains of the experimentally-determined structures are the same as in **Figure 5**. From top row to bottom row, structures are taken from: the *hotpocketNN* training set, the *hotpocketNN* validation set, the *hotpocketNN* testing set, human protein structures with BioLiP annotations not included in the *hotpocketNN* train/val/test sets, and human protein structures with BioLiP annotations released in 2024 with low sequence identity to previously-seen structures. The surface of the protein structure is colored as follows: magenta for residues that are part of an accepted candidate pocket accepted by *hotpocketNN*, blue for residues that are part of a candidate pocket but not an accepted candidate pocket, and light green for residues that are not part of any candidate pockets.

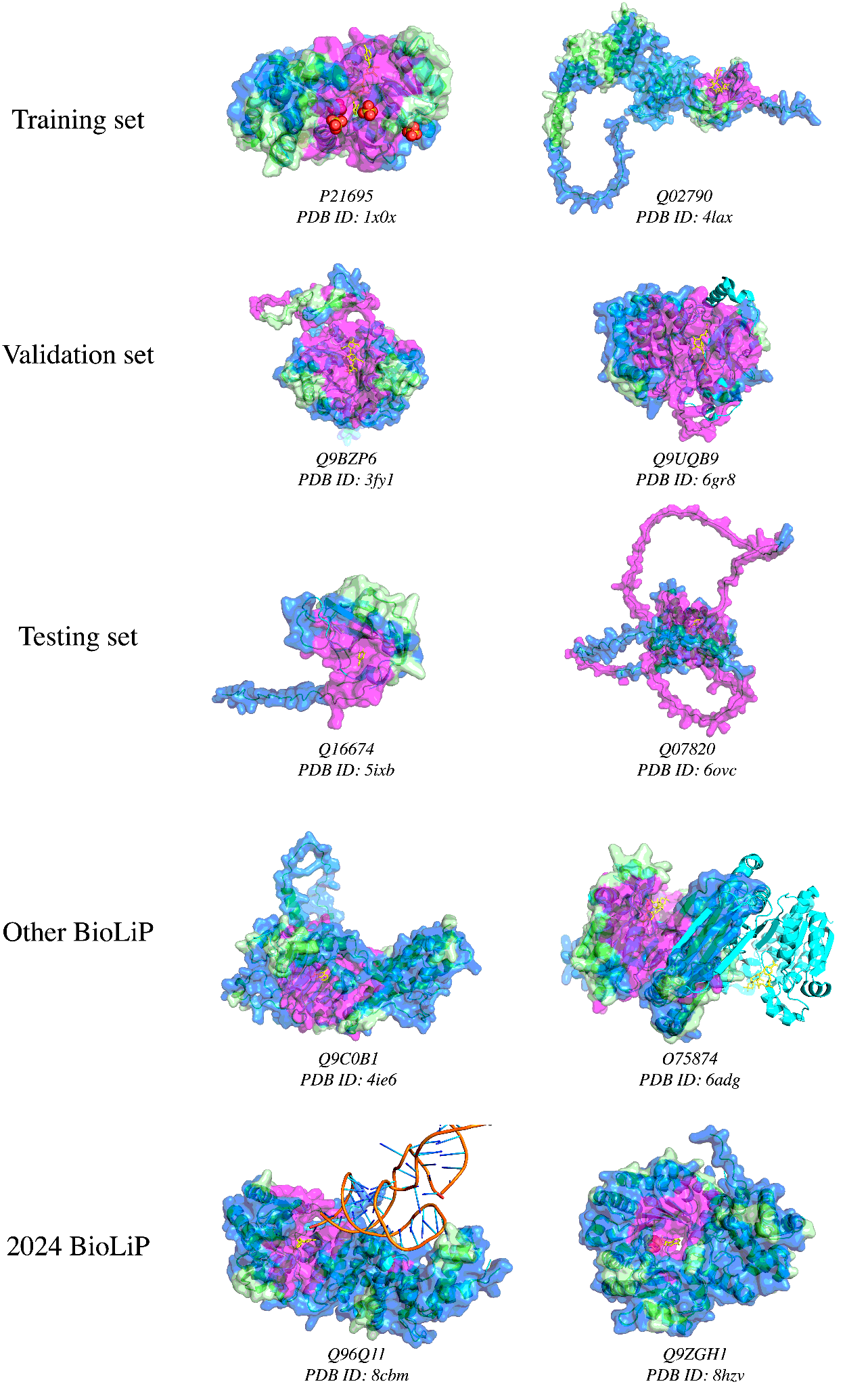

**Figure S6.** Visualizations of accepted and rejected candidate pockets on AlphaFold2-predicted protein structures, using constituent method predictions and ESM2 embeddings as features for *hotpocketNN* ensembling and filtering method. AlphaFold2-predicted structures (green ribbons) are shown aligned with their experimentally-determined PDB counterparts (cyan ribbons). Only the surface of the predicted structure is shown and pocket predictions were made on the predicted structure. The aligned ligands from the experimentally-determined structure are shown in yellow; these ligands are not a part of the AlphaFold2-predicted structures. The chains of the experimentally-determined structures are the same as in **Figure S3**. From top row to bottom row, structures are taken from: the *hotpocketNN* training set, the *hotpocketNN* validation set, the *hotpocketNN* testing set, human protein structures with BioLiP annotations not included in the *hotpocketNN* train/val/test sets, and human protein structures with BioLiP annotations released in 2024 with low sequence identity to previously-seen structures. The surface of the protein structure is colored as follows: magenta for residues that are part of an accepted candidate pocket accepted by *hotpocketNN*, blue for residues that are part of a candidate pocket but not an accepted candidate pocket, and light green for residues that are not part of any candidate pockets.

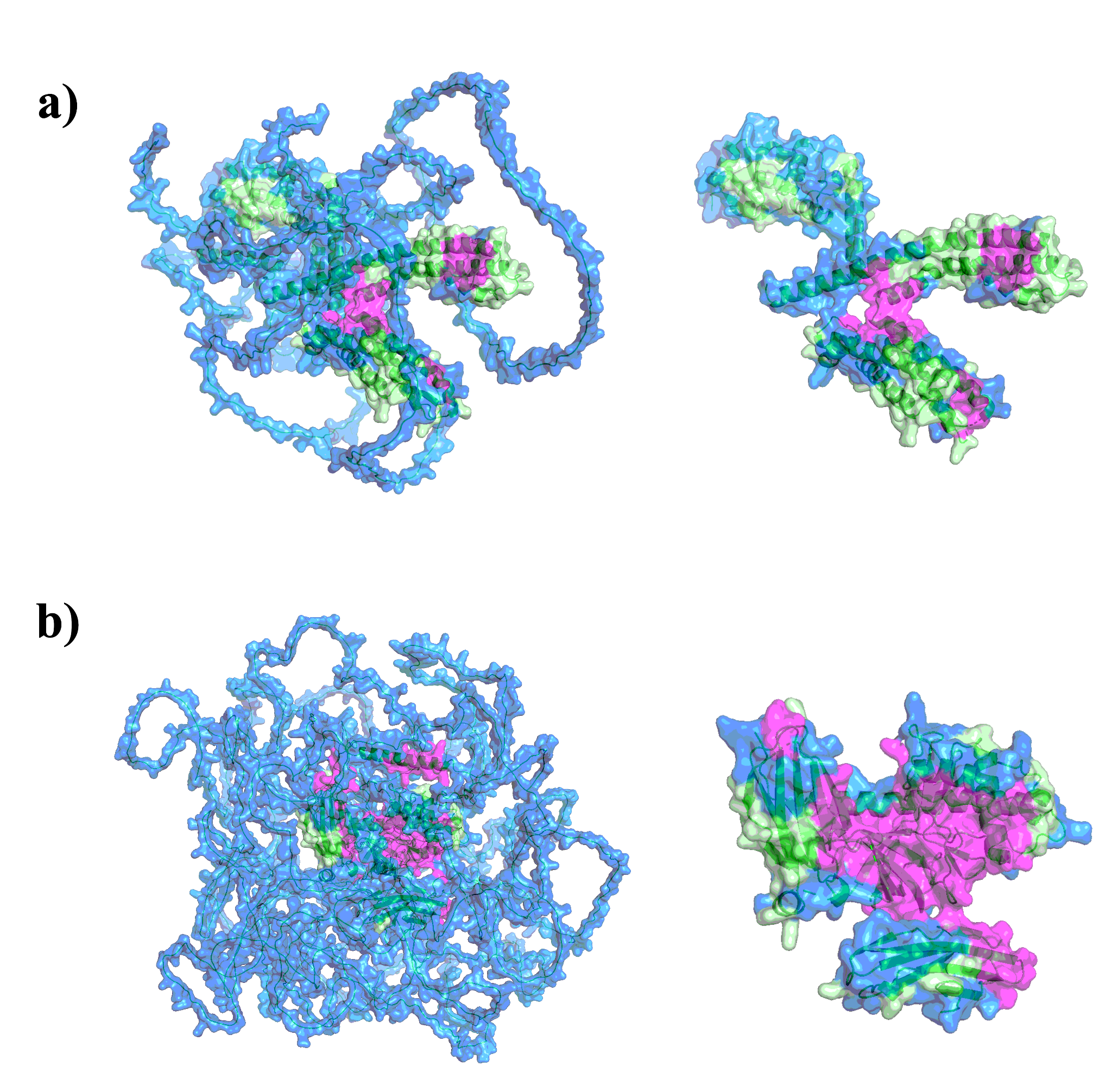

**Figure S7.** Visualizations of accepted and rejected candidate pockets on AlphaFold2-predicted protein structures with an average per-residue confidence that is **a)** “low” (average pLDDT between 50 and 70; UniProt ID: Q92953), or **b)** “very low” (average pLDDT less than 50; UniProt ID: Q86TB3), using ESM2 embeddings as features for *hotpocketNN* ensembling and filtering method. Neither protein has an experimentally-determined protein structure. The surface of the protein structure is colored as follows: magenta for residues that are part of an accepted candidate pocket accepted by *hotpocketNN*, blue for residues that are part of a candidate pocket but not an accepted candidate pocket, and light green for residues that are not part of any candidate pockets. Both the full predicted protein structure (left) and the subset of the structure that is not low-confidence (pLDDT greater than 70) (right) are shown.

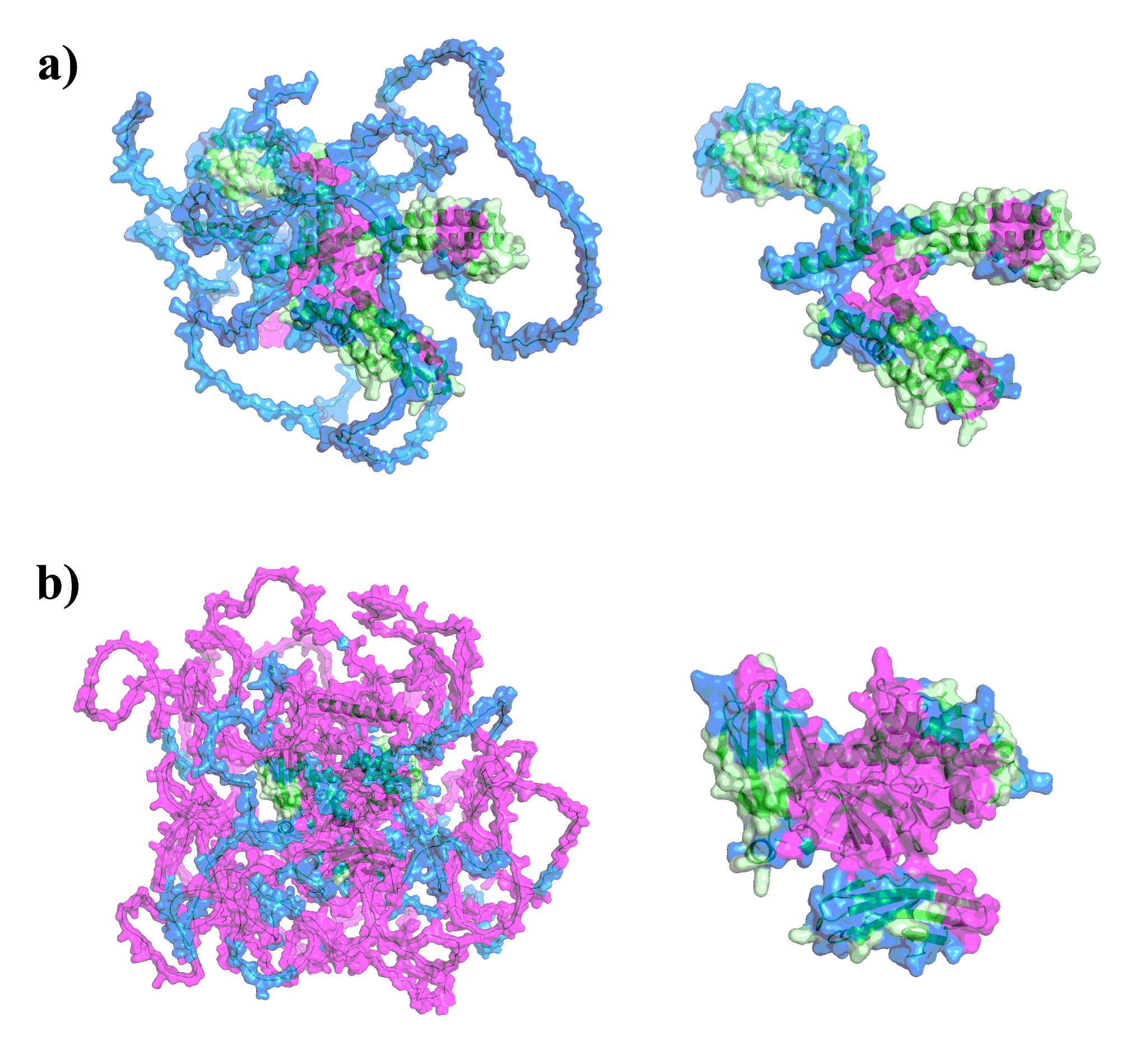

**Figure S8.** Visualizations of accepted and rejected candidate pockets on AlphaFold2-predicted protein structures with an average per-residue confidence that is **a)** “low” (average pLDDT between 50 and 70; UniProt ID: Q92953), or **b)** “very low” (average pLDDT less than 50; UniProt ID: Q86TB3), using constituent method predictions and ESM2 embeddings as features for *hotpocketNN* ensembling and filtering method. Neither protein has an experimentally-determined protein structure. The surface of the protein structure is colored as follows: magenta for residues that are part of an accepted candidate pocket accepted by *hotpocketNN*, blue for residues that are part of a candidate pocket but not an accepted candidate pocket, and light green for residues that are not part of any candidate pockets. Both the full predicted protein structure (left) and the subset of the structure that is not low-confidence (pLDDT greater than 70) (right) are shown.

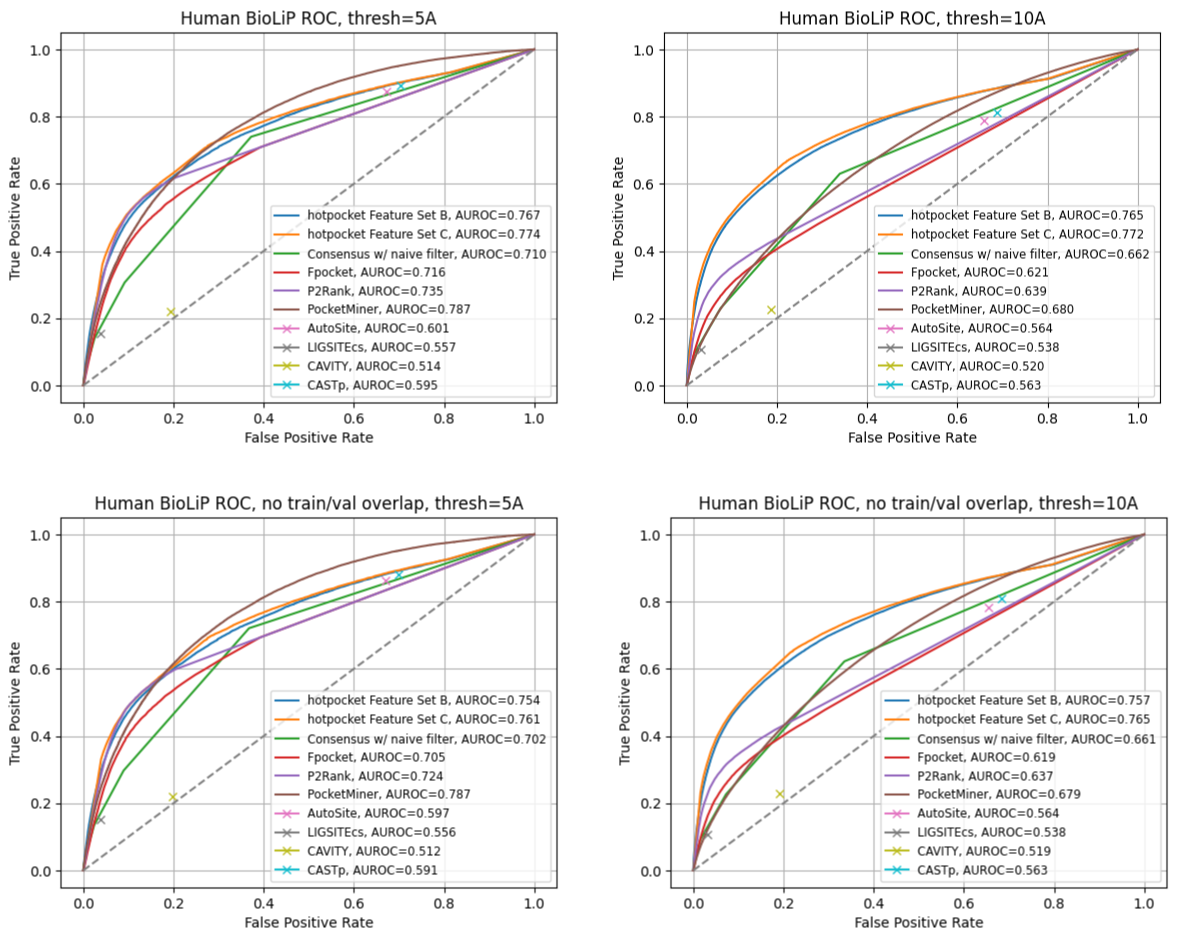

**Figure S9.** ROC curves and AUROCs for per-residue scoring performance of *hotpocketNN*, naive filter across union of constituent methods, and each constituent pocket-finding method individually across the Human BioLiP dataset. Each residue was labeled as a binding residue if it was within 5 angstroms (left) or 10 angstroms (right) of a relevant ligand. For the *hotpocketNN* models, Fpocket, P2Rank, and PocketMiner, each residue’s predicted score for being a member of a pocket was the maximum of all predicted pockets of which it was a member. AutoSite, LIGSITEcs, CAVITY, and CASTp do not provide pocket scores; for these methods, each residue’s predicted score for being a member of a pocket was 1 if it is part of any predicted pocket for the structure, or 0 otherwise. The top row shows results when all structures in all datasets are included, including structures present in the training and validation sets for the *hotpocketNN*. The bottom row shows results when structures present in the training and validation sets for the *hotpocketNN* are excluded. CAVITY and CASTp did not have predictions available for any PoseBusters structures. Feature Set B is the per-residue ESM2 embeddings; Feature Set C is both the per-residue pocket predictions and per-residue ESM2 embeddings concatenated together.

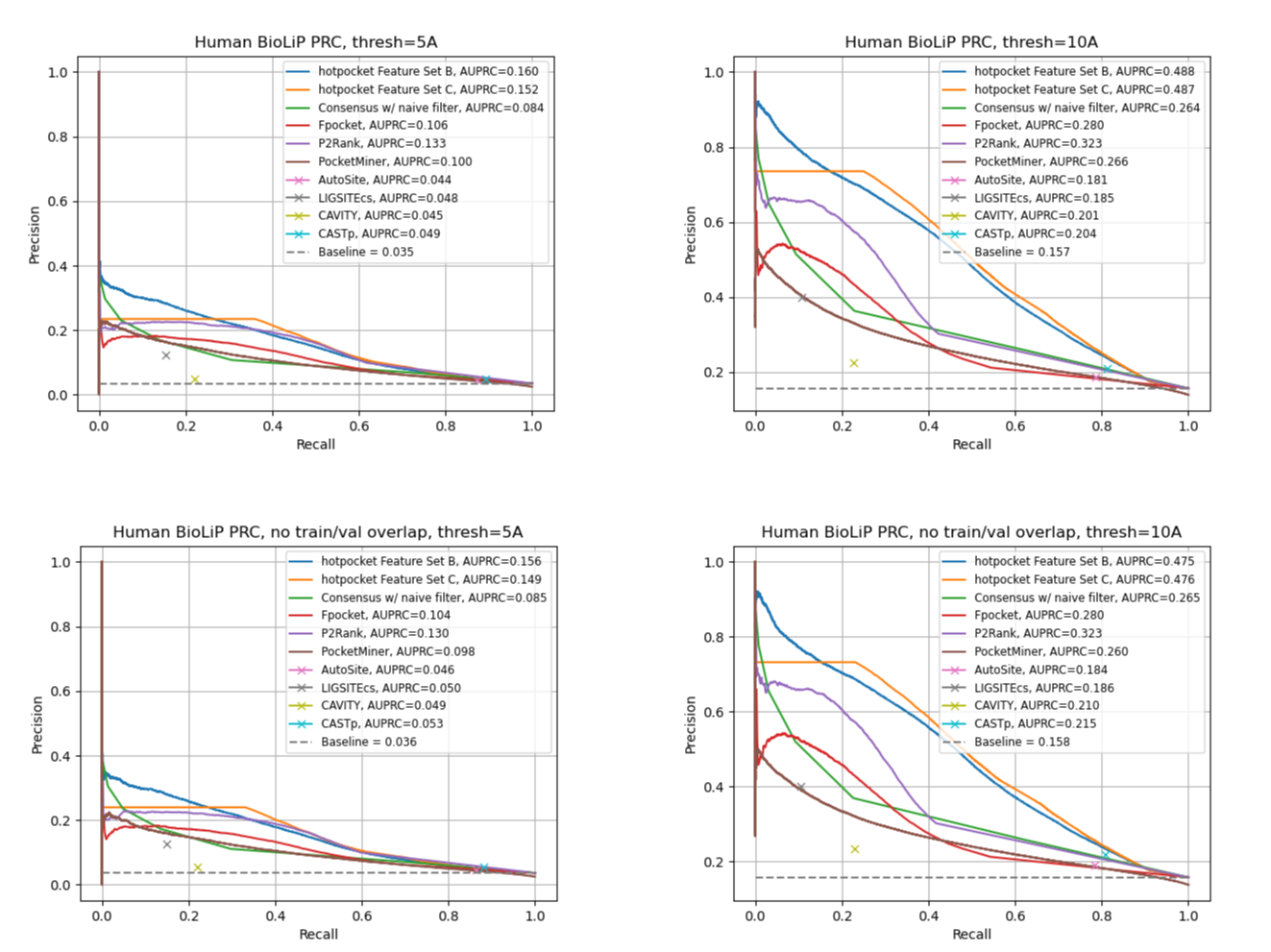

**Figure S10.** PRC curves and AUPRCs for per-residue scoring performance of *hotpocketNN*, naive filter across union of constituent methods, and each constituent pocket-finding method individually across the Human BioLiP dataset. Each residue was labeled as a binding residue if it was within 5 angstroms (left) or 10 angstroms (right) of a relevant ligand. For the *hotpocketNN* models, Fpocket, P2Rank, and PocketMiner, each residue’s predicted score for being a member of a pocket was the maximum of all predicted pockets of which it was a member. AutoSite, LIGSITEcs, CAVITY, and CASTp do not provide pocket scores; for these methods, each residue’s predicted score for being a member of a pocket was 1 if it is part of any predicted pocket for the structure, or 0 otherwise. The top row shows results when all structures in all datasets are included, including structures present in the training and validation sets for the *hotpocketNN*. The bottom row shows results when structures present in the training and validation sets for the *hotpocketNN* are excluded. CAVITY and CASTp did not have predictions available for any PoseBusters structures. Feature Set B is the per-residue ESM2 embeddings; Feature Set C is both the per-residue pocket predictions and per-residue ESM2 embeddings concatenated together.
